# Analysis of human visual experience data

**DOI:** 10.1101/2025.08.11.669764

**Authors:** Johannes Zauner, Aaron Nicholls, Lisa A. Ostrin, Manuel Spitschan

## Abstract

Exposure to the optical environment — often referred to as *visual experience* — profoundly influences human physiology and behavior across multiple time scales. In controlled laboratory settings, stimuli can be held constant or manipulated parametrically. However, such exposures rarely replicate real-world conditions, which are inherently complex and dynamic, generating high-dimensional datasets that demand rigorous and flexible analysis strategies. This tutorial presents an analysis pipeline for visual experience datasets, with a focus on reproducible workflows for human chronobiology and myopia research. Light exposure and its retinal encoding affect human physiology and behavior across multiple time scales. Here we provide step-by-step instructions for importing, visualizing, and processing viewing distance and light exposure data. This includes time-series analyses for working distance, biologically relevant light metrics, and spectral characteristics. The tasks are standardized through the open-source R package <u>LightLogR</u>. By leveraging a modular approach, the tutorial supports researchers in building flexible and robust pipelines that accommodate diverse experimental paradigms and measurement systems.

## Introduction

Exposure to the optical environment — often referred to as *visual experience* — profoundly influences human physiology and behavior across multiple time scales. Two notable examples, from distinct research domains, can be understood through a common retinally-referenced framework.

The first example relates to the non-visual effects of light on human circadian and neuroendocrine physiology. The light–dark cycle entrains the circadian clock, and light exposure at night suppresses melatonin production (Brown et al. 2022; Blume, Garbazza, and Spitschan 2019). The second example concerns the influence of visual experience on ocular development, particularly myopia. Time spent outdoors — which features distinct optical environments — has been consistently associated with protective effects on ocular growth and health outcomes (Dahlmann-Noor et al. 2025).

In controlled laboratory settings, light exposure can be held constant or manipulated parametrically. In contrast, real-world conditions are inherently complex and dynamic, and cannot be captured by single spot measurements. As people move in and between spaces (indoors and outdoors) and move their body, head, and eyes, exposure to the optical environment varies significantly (Webler et al. 2019) and is modulated by behavior (Biller, Balakrishnan, and Spitschan 2024). Wearable devices for measuring light exposure have thus emerged as vital tools to capture the ecological visual experience. These tools generate high-dimensional datasets that demand rigorous and flexible analysis strategies.

Starting in the 1980s (Okudaira, Kripke, and Webster 1983), technology to measure optical exposure has matured, with miniaturized illuminance sensors now (in 2025) very common in consumer wearables (van Duijnhoven et al. 2025). In research, several devices are available that differ in functionality, ranging from small pins measuring ambient illuminance (Mohamed et al. 2021) to head-mounted multi-modal devices capturing nearly all relevant aspects of visual experience (Gibaldi et al. 2024). Increased capabilities in wearables bring complex, dense datasets. These go hand-in-hand with a proliferation of metrics, as highlighted by recent review papers in both circadian and myopia research (Hönekopp and Weigelt 2023; Hartmeyer and Andersen 2023).

At present, the analysis processes to derive metrics are often implemented on a per-laboratory or even per-researcher basis. This fragmentation is a potential source of errors and inconsistencies between studies, consumes considerable researcher time (Hartmeyer, Webler, and Andersen 2022), and these bespoke processes and formats hinder harmonization or meta-analysis across multiple studies. It is very common that more time is spent preparing data than gaining insights through rigorous statistical analysis. These preparation tasks are best handled, or at least facilitated, by standardized, transparent, community-based analysis pipelines (Zauner, Udovicic, and Spitschan 2024).

In circadian research, the R package LightLogR was developed to address this need (Zauner, Hartmeyer, and Spitschan 2025). LightLogR is an open-source, MIT-licensed, community-driven package specifically designed for data from wearable light loggers and optical radiation dosimeters. It contains functions to calculate over sixty different metrics used in the field (Hartmeyer and Andersen 2023). The package functions come with light-related defaults, but they remain fundamentally agnostic to modality. As a result, parameters like viewing distance and light spectra, both highly relevant to myopia research (Hönekopp and Weigelt 2023), can easily be handled.

In this article, we demonstrate that LightLogR’s analysis pipelines and metric functions apply broadly across the field of visual experience research, not just to circadian rhythms and chronobiology. Our approach is modular and extensible, allowing researchers to adapt it to a variety of devices and research questions. Emphasis is placed on clarity, transparency, and reproducibility, aligning with best practices in scientific computing and open science. We use example data from two devices (worn by different individuals and at different times) to showcase the LightLogR workflow with metrics relevant to myopia research, covering working distance, (day)light exposure, and spectral analysis. Readers are encouraged to recreate the analysis using the provided code. All necessary data and code are openly available in the <u>GitHub repository</u>.

### Scope

This article focuses on workflows for deriving condensed metrics from time-series data collected with wearable devices in the visual-experience domain. Specifically, we address illuminance, viewing distance, and spectral irradiance. Example datasets from two types of wearable devices are used for illustration.

Many relevant considerations arise when collecting data with wearable devices. This article covers only a subset of these. In particular, it does not address:

- device selection (see, e.g., van Duijnhoven et al. 2025; Zauner et al. 2025a)
- measurement accuracy or device calibration
- auxiliary data such as sleep/wake information (see, e.g., Zauner et al. 2025b; Guidolin et al. 2024)

More information on those aspects can be found in the *Technical guide for wearable optical radiation dosimetry and visual experience assessment* (Zauner et al. 2025c).

To demonstrate the workflows, this article uses expert-informed definitions of metrics and metric parameters (see, e.g., **Table 1**, **Table 2**, and the non-wear detection rules based on activity data described in <u>Supplement 1</u>). These definitions and thresholds should not be interpreted as universal standards, nor are they hard-coded into the software package. For any application, parameter choices must be tailored to the research domain, study context and design, and the specifications of the wearable device.

**Table 1:** Overview of metrics. In all cases, the averages for weekday, weekend, and the mean daily value are calculated. Definitions are shown in Table 2.

| No. | Name | Implementation<sup>1</sup> |
| --- | --- | --- |
| **Distance** |  |  |
| 1 | Total wear time daily | durations() * |
| 2 | Duration of<br>Near work,<br>Intermediate Work,<br>Near + Intermediate Work, or<br>per each *Distance range* | filter for distance range +<br>durations() * (for single ranges)<br>or<br>grouping by distance range +<br>durations() * (for all ranges) |
|  | (10cm steps) |  |
| 3 | Frequency of Continuous near work | extract_clusters()* + summarize_numeric()* |
| 4 | Frequency, duration, and distances of *Near Work episodes* | extract_clusters()* + extract_metric()* + summarize_numeric()* |
| 5 | Frequency and duration of *Visual breaks* | extract_clusters()* + filter |
| Light |  |  |
| 6 | Light exposure (in lux) | summarize_numeric()* |
| 7 | Duration per *Outdoor range* | grouping by Outdoor range + durations()* |
| 8 | The number of times light level changes from indoor (<1000 lx) to outdoor (>1000 lx) | extract_states()* + summarize_numeric()* |
| 9 | Longest period above 1000 lx | period_above_threshold()* |
| Spectrum |  |  |
| 10 | Ratio of short vs. long wavelength light | spectral_integration()* + summarize_numeric()* |
| 11 | Melanopic daylight efficacy ratio (MDER) | spectral_integration()* + summarize_numeric()* |
| 12 | Short-wavelength light at certain times of day | spectral_integration()* + filter_Time()* (for defined times) or cut_Datetime()* (for regular time intervals) or add_photoperiod()* (for solar times) + grouping by time state + summarize_numeric()* |

<sup>1</sup> Functions from `LightLogR` are presented with an \*. Non-asterisk functions refer to tidyverse functions.

**Table 2:**
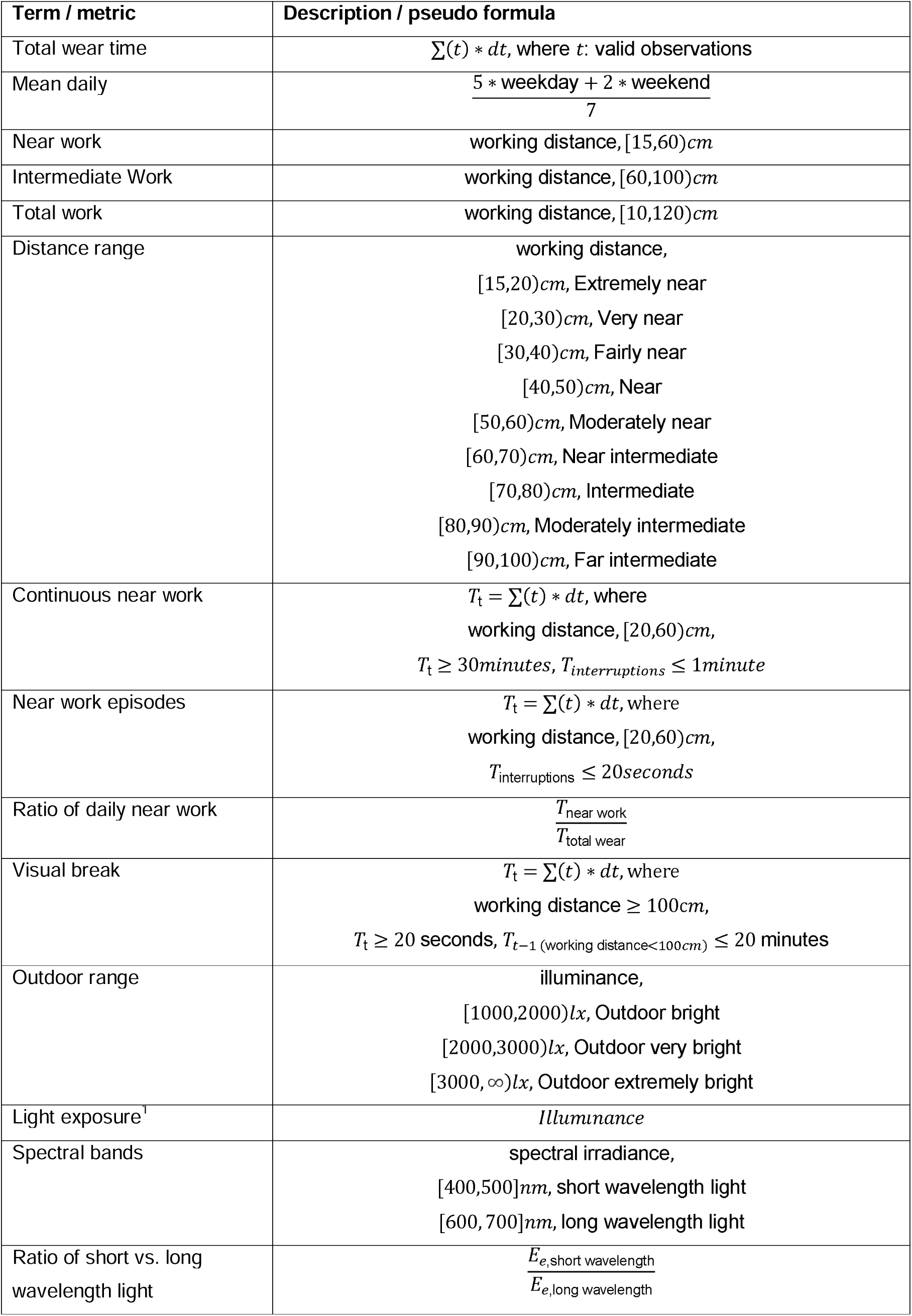
Definitions of terms and metrics

Further, the article is split up in the main analysis part, where all metrics are calculated, and the <u>Supplement 1</u>, where data is imported, screened, and prepared. Thus, the reader is referred to <u>Supplement 1</u> for all aspects regarding data formats, preparation steps, and handling of gaps, i.e., missing data.

Lastly, the example data used in the article do not stem from a controlled experimental data collection but consist of pilot data gathered in an ecological setting without a fixed protocol. Given the substantial interindividual differences in visual experience metrics, and because the analyses focus on one participant at a time, the reported results should be interpreted as illustrative rather than representative of typical or population-level values.

## Method and Materials

### Software

This tutorial was built with Quarto, an open-source scientific and technical publishing system that integrates text, code, and code output into a single document. The source code to reproduce all results is included and accessible via the Quarto document’s code tool menu. All analyses were conducted in *R* (version 4.5.0, “How About a Twenty-Six”) using *LightLogR* (version 0.10.0 “High noon”). We also used the *tidyverse* suite (version 2.0.0) for data manipulation (which LightLogR follows in its design), and the *gt* package (version 1.1.0) for generating summary tables. A comprehensive overview of the R computing environment is provided in the session info (see <u>Session info</u> section of the online tutorial).

### Metric selection and definitions

In March 2025, two workshops with myopia researchers — initiated by the Research Data Alliance (RDA) Working Group on Optical Radiation Exposure and Visual Experience Data — focused on current needs and future opportunities in data analysis, including the development and standardization of metrics. Based on expert input from these workshops, the authors of this tutorial compiled a list of visual experience metrics, shown in **Table 1**. These include many currently used metrics and definitions (Wen et al. 2020, 2019; Bhandari and Ostrin 2020; Williams et al. 2019), as well as new metrics enabled by spectrally-resolved measurements. While they are not derived by a formal consensus process, they are expert-informed and used in current scientific research, and thus will serve as example-definitions for metrics and thresholds throughout this article.

Table 2 provides definitions for the terms used in **Table 1**. Note that specific definitions may vary depending on the research question or device capabilities.

It should be noted that although daylight levels can far exceed the thresholds defined in **Table 2** - and may reach or even exceed 10^5 lux - empirical daylight levels measured at eye level are much lower, typically around 10^3 lux, especially when considering aggregated time-series data over minutes to hours.

### Devices

Data from two wearable devices are used in this analysis:

· Clouclip: A wearable device that measures viewing distance and ambient light simultaneously [Glasson Technology Co., Ltd, Hangzhou, China; Wen et al. (2021); Wen et al. (2020)]. The Clouclip provides a simple data output with only distance (working distance, in centimeters) and illuminance (ambient light, in lux). Data in our example were recorded at 5-second intervals. Approximately one week of data (∼120,960 observations) is about 1.6 MB in size.
· Visual Environment Evaluation Tool (VEET): A head-mounted multi-modal device that logs multiple data streams [Reality Labs Research, Menlo Park, CA, USA; Sah, Narra, and Ostrin (2025); Sullivan et al. (2024)]. The VEET dataset used here contains simultaneous measurements of distance (via a time-of-flight sensor), ambient light (illuminance), activity (accelerometer & gyroscope), and spectral irradiance (multi-channel light sensor). Data were recorded at 2-second intervals, yielding a very dense dataset (∼270 MB per week).

### Data processing summary

The <u>Results</u> section uses imported and pre-processed data from the two devices to calculate metrics. <u>Supplement 1</u> contains the annotated code and description for the steps involved, which are summarized as follows, and as shown in Figure 1. Please refer to the supplement for details.

**Figure 1:**
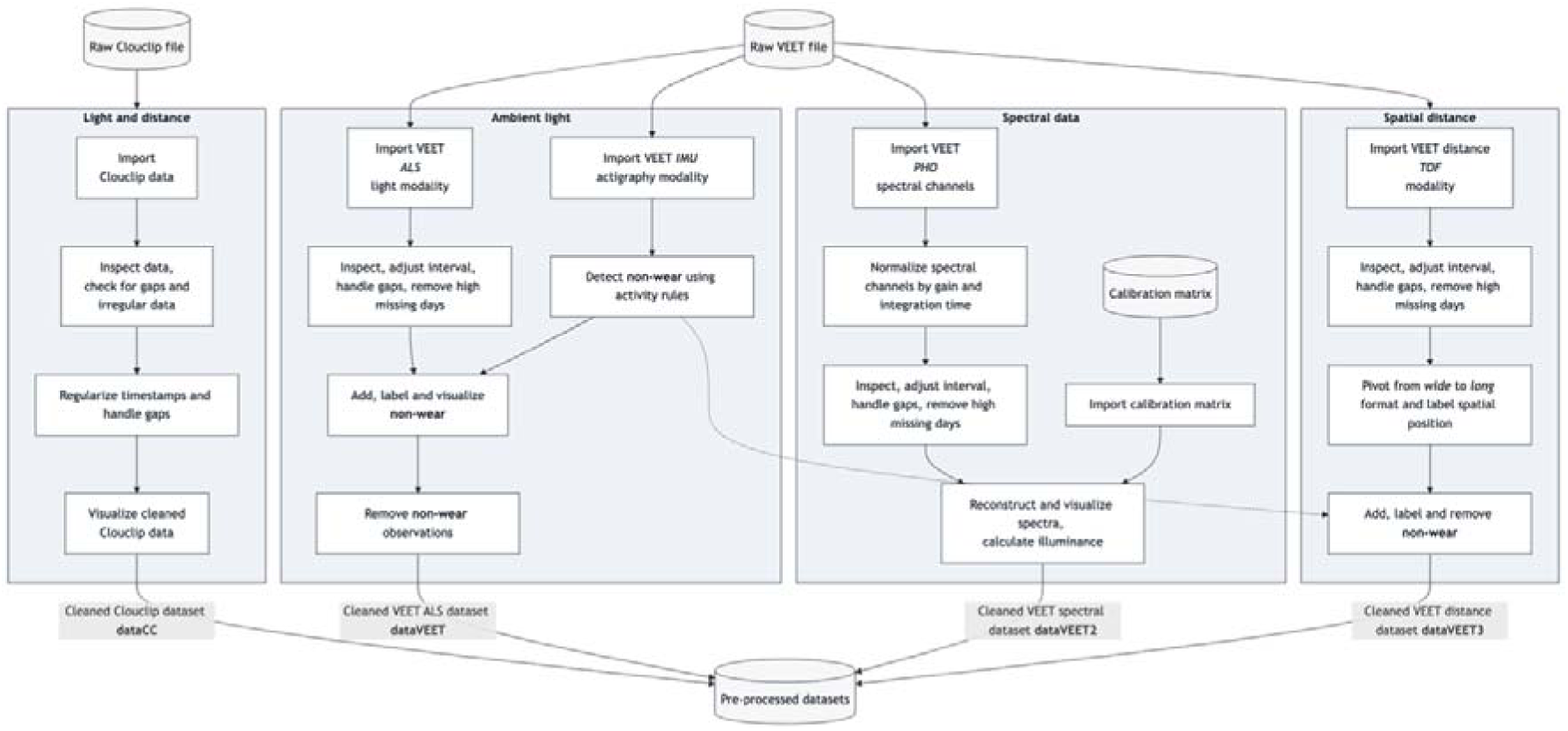
Pre-processing steps in the supplement document

**Data import:** We imported raw data from the Clouclip and VEET devices using LightLogR’s built-in import functions, which automatically handle device-specific formats and idiosyncrasies.

The Clouclip export file (provided as a tab-delimited text file) contains timestamped records of distance (cm) and illuminance (lux). LightLogR’s import$Clouclip function reads this file, after specifying the device’s recording timezone, and converts device-specific sentinel codes into proper missing values. For instance, the Clouclip uses special numeric codes to indicate when it is in “sleep mode” or when a reading is out of the sensor’s range, rather than recording a normal value. LightLogR identifies -1 (for both distance and lux) as indicating the device’s sleep mode and 204 (for distance) as indicating the object was beyond the measurable range, replacing these with NA and logging their status in separate columns. The import routine also provides an initial summary of the dataset, including start and end times and any irregular sampling intervals or gaps.

For the VEET device, data were provided as CSV logs (zipped on Github, due to size). We focused on the ambient light sensor modality first. Using import$VEET(…, modality = "ALS"), we extracted the illuminance (Lux) data stream and its timestamps. The raw VEET data similarly contains irregular intervals and can contain missing periods (e.g., if the device stopped recording or was reset); the import summary flags these issues.

Besides the Clouclip and VEET, LightLogR 0.10.0 contains import functions for 18 more wearable devices. The package further supports versions due to evolving data formats, and includes documentation for both <u>code-based</u> and <u>code-less</u> additions of new device import-functions.

### Irregular intervals, gaps, non-wear times

Both datasets showed irregular timing and missing data, i.e., gaps. Irregular data means that some observations did not align to the nominal sampling interval (e.g., slight timing drift or pauses in recording). For the Clouclip 5-second data, we detected irregular timestamps spanning all but the first and last day of the recording. Handling such irregularities is important because many downstream analyses assume a regular time series. We evaluated strategies to address this, including:

- Removing an initial portion of data if irregularities occur mainly during device start-up.
- Rounding all timestamps to the nearest regular interval (5 s in this case).
- Aggregating to a coarser time interval (with some loss of temporal resolution).

Based on the import summary and visual inspection of the time gaps, we chose to round the observation times to the nearest 5-second mark, as this addressed the minor offsets without significant data loss. After rounding timestamps, we added an explicit date column for convenient grouping by day.

We then generated a summary of missing data for each day. Implicit gaps (intervals where the device should have recorded data but did not) were converted into explicit missing entries using LightLogR’s gap-handling functions. We also removed days that had very little data to focus on days with substantial wear time. In our Clouclip example, days with <1 hour of recordings were dropped. This threshold should be adjusted based on how much complete days matter for a given analysis at hand.

E.g., in circadian science, the metrics of *interdaily stability* and *intradaily variation* require measurements for each hour of the day.

After these preprocessing steps, the Clouclip dataset had no irregular timestamps remaining and contained explicit markers for all periods of missing data (e.g., times when the device was off or not worn). The distance and illuminance values were now ready for metric calculations. Because the device was put in sleep mode when not worn, there are no measurements during non-wear times.

The VEET illuminance data underwent a similar cleaning procedure. To make the VEET’s 2-second illuminance data more comparable to the Clouclip’s and to reduce computational load, we aggregated the illuminance time series to 5-second intervals. Aggregation was performed with the arithmetic mean of values in a 5-second bin.

We then inserted explicit missing entries for each whole day and removed days with more than one hour of missing illuminance data. After cleaning, six days of VEET illuminance data with good coverage remained for analysis (see <u>Supplement 1</u> for details).

Finally, for spectral analysis, we imported the VEET’s spectral sensor modality, and, for the distance analysis, the time-of-flight modality. This required additional processing: the raw spectral data consists of counts from nine wavelength-specific channels (approximately 415 nm through 940 nm, unequally spaced between 30 and 50 nm, plus one broadband clear channel covering the whole range of individual channels, another broadband channel for flicker detection, and a dark channel) along with a sensor gain setting. We aggregated the spectral data to 5-minute intervals to focus on broader trends and reduce data volume. Each channel’s counts were normalized by the appropriate gain. Using a calibration matrix provided by the manufacturer (specific to the spectral sensor model), we reconstructed full spectral power distributions for each 5-minute interval. The end result is a list-column in the dataset where each entry is the estimated spectral irradiance across wavelengths for that time interval. Detailed spectral preprocessing steps, including the calibration and normalization, are provided in the <u>Supplement 1</u>. After spectral reconstruction, the dataset was ready for calculating example spectrum-based metrics.

Similarly, the time-of-flight modality contains 256 values per observation, encoding an 8x8 grid of distance and confidence measurements for up to two objects (8x8 grid, times two objects, times distance + confidence column for each object and grid point -> 256 values). For computational reasons, only the first object was kept.

These data were pivoted into a long format, where each row contains the distance and confidence data for a given position in the grid and a given datetime. After pivoting and converting grid positions into a deviation angle from central view, the dataset was ready to be used for distance analysis.

Because the VEET devices record even when not worn, a non-wear detection using the devices’ actigraphy modality was implemented. This process used the standard deviation of a linear motion sensor in a 5-minute bin with a visually derived threshold to separate wear from non-wear time. Measurements of illuminance and distance were consequently removed during the calculated non-wear times.

This tutorial will start by importing a Clouclip dataset and providing an overview of the data. The Clouclip export is considerably simpler compared to the VEET export, only containing Distance and Illuminance measurements. The VEET dataset will be imported later for the spectrum related metrics.

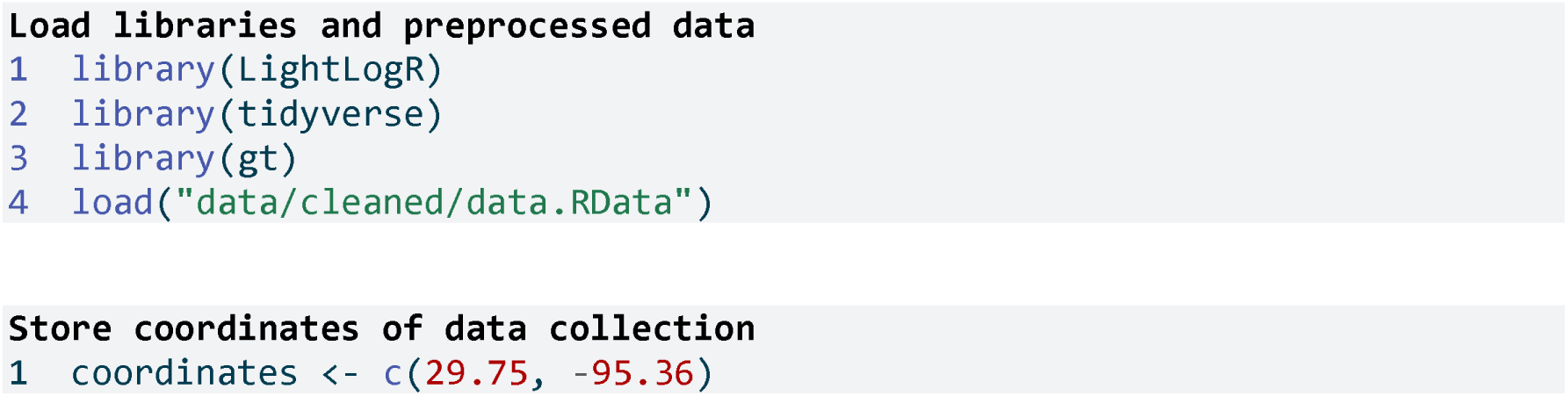

**Line 1** Coordinates for Houston, Texas; coordinates are important to calculate and visualize photoperiods later

## Results

Figure 2 shows an overview of the covered workflows in the results section.

**Figure 2:**
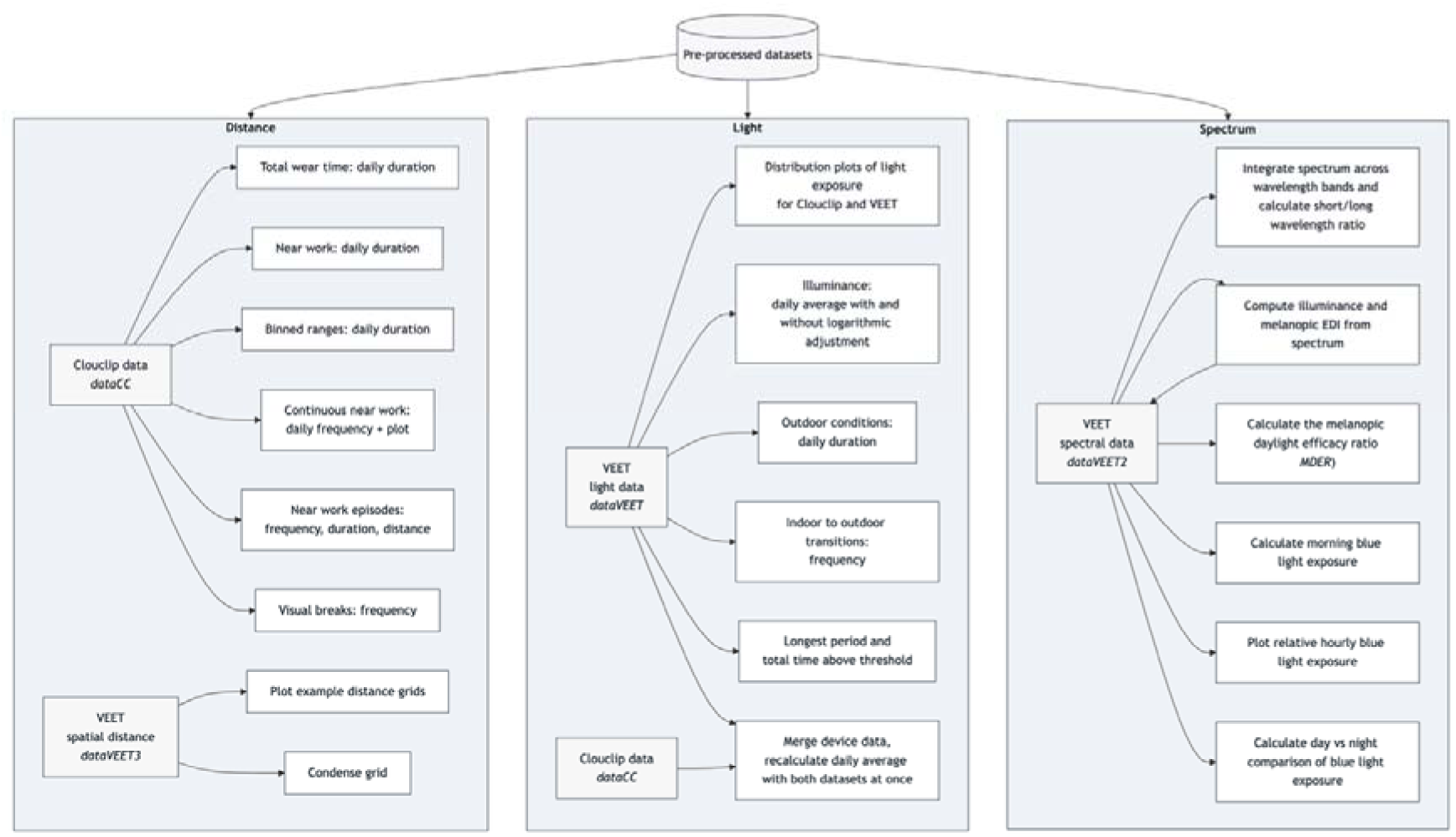
Workflows that are covered in the results section. The pre-processed datasets are covered in detail in the supplement

We first examine metrics related to viewing distance, using the processed Clouclip dataset. Many distance-based metrics are computed for each day and then averaged over weekdays, weekends, or across all days. To facilitate this, we define a helper function that will take daily metric values and calculate the mean values for weekdays, weekends, and the overall daily average:

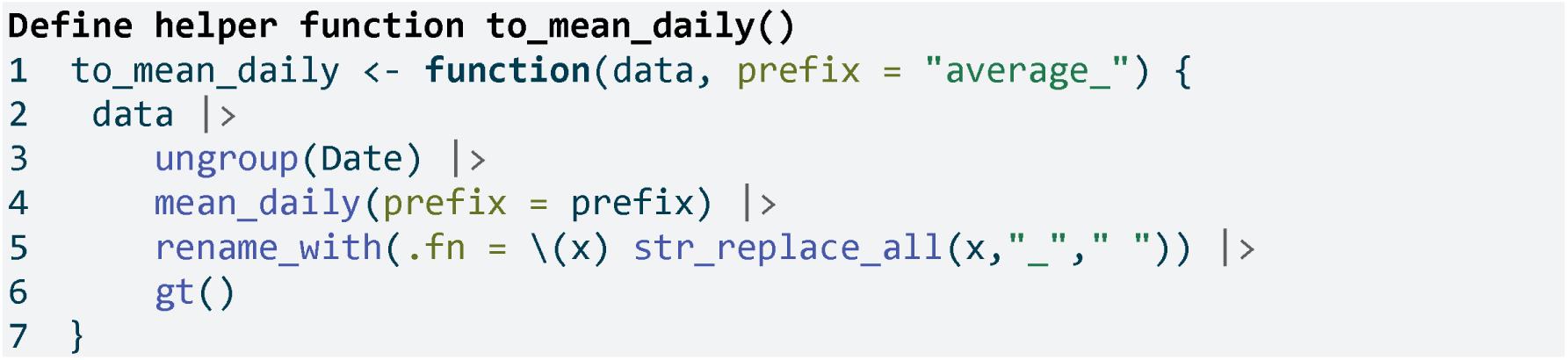

**Line 3** Ungroup by days

**Line 4** Calculate the averages per grouping

**Line 5** Remove underscores in names

**Line 6** Format as a gt table for display

### Total wear time daily

*Total wear time daily* refers to the amount of time the device was actively collecting distance data each day (i.e. the time the device was worn and operational). We compute this by summing all intervals where a valid distance measurement is present, ignoring periods where data are missing or the device was off. The results are shown in **Table 3**.

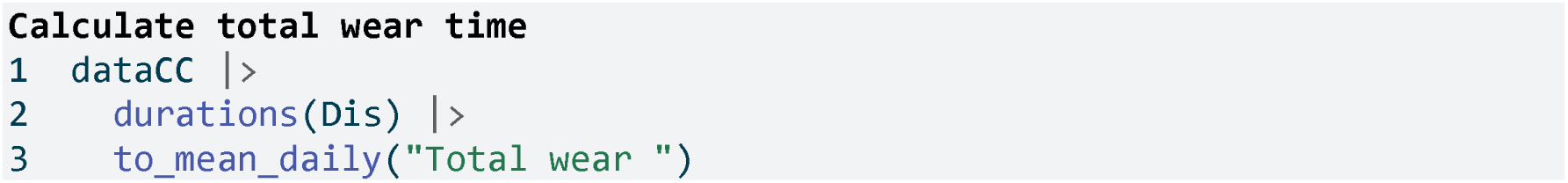

**Table 3:** Total wear time per day (average across days)

|  | Total wear duration |
| --- | --- |
| Mean daily | 31448s (~8.74 hours) |
| Weekday | 34460s (~9.57 hours) |
| Weekend | 23918s (~6.64 hours) |

**Line 2** Calculate total duration of data per day

**Line 3** Using the helper function defined above

### Duration within distance ranges

Many myopia-relevant metrics concern the time spent at certain viewing distances (e.g., “near work” vs. intermediate or far distances). We calculate the duration of time spent in specific distance ranges. **Table 4** shows the average daily *duration of near work*, defined here as time viewing at 15–60 cm. **Table 5** provides a more detailed breakdown across multiple distance bands.

**Table 4:**
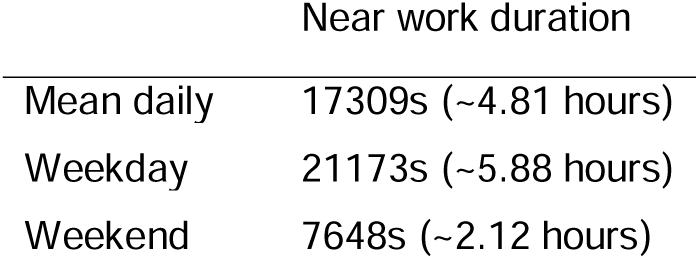
Daily duration of near work (15–60 cm viewing distance)

**Table 5:** Daily duration in each viewing distance range in minutes. For better readability, the table was transposed (compared to the code results)

|  | Mean daily | Weekday | Weekend |
| --- | --- | --- | --- |
| Far | 16m | 20m | 5m |
| Far intermediate | 11m | 14m | 2m |
| Moderately intermediate | 5m | 6m | 3m |
| Intermediate | 4m | 6m | 1m |
| Near intermediate | 7m | 7m | 8m |
| Moderately near | 13m | 16m | 5m |
| Near | 27m | 36m | 5m |
| Fairly near | 46m | 60m | 12m |
| Very near | 102m | 128m | 38m |
| Extremely near | 74m | 88m | 40m |

### Duration of near work

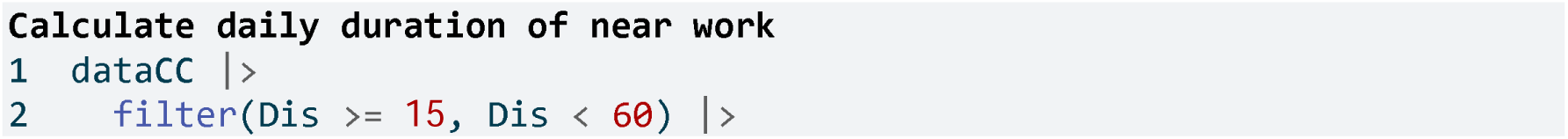
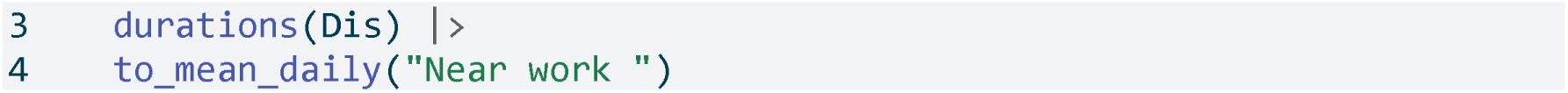

**Line 2** Consider only distances in [15, 60) cm

**Line 3** Total duration in that range per day

### Duration within distance ranges

First, we define a set of distance breakpoints and descriptive labels for each range:

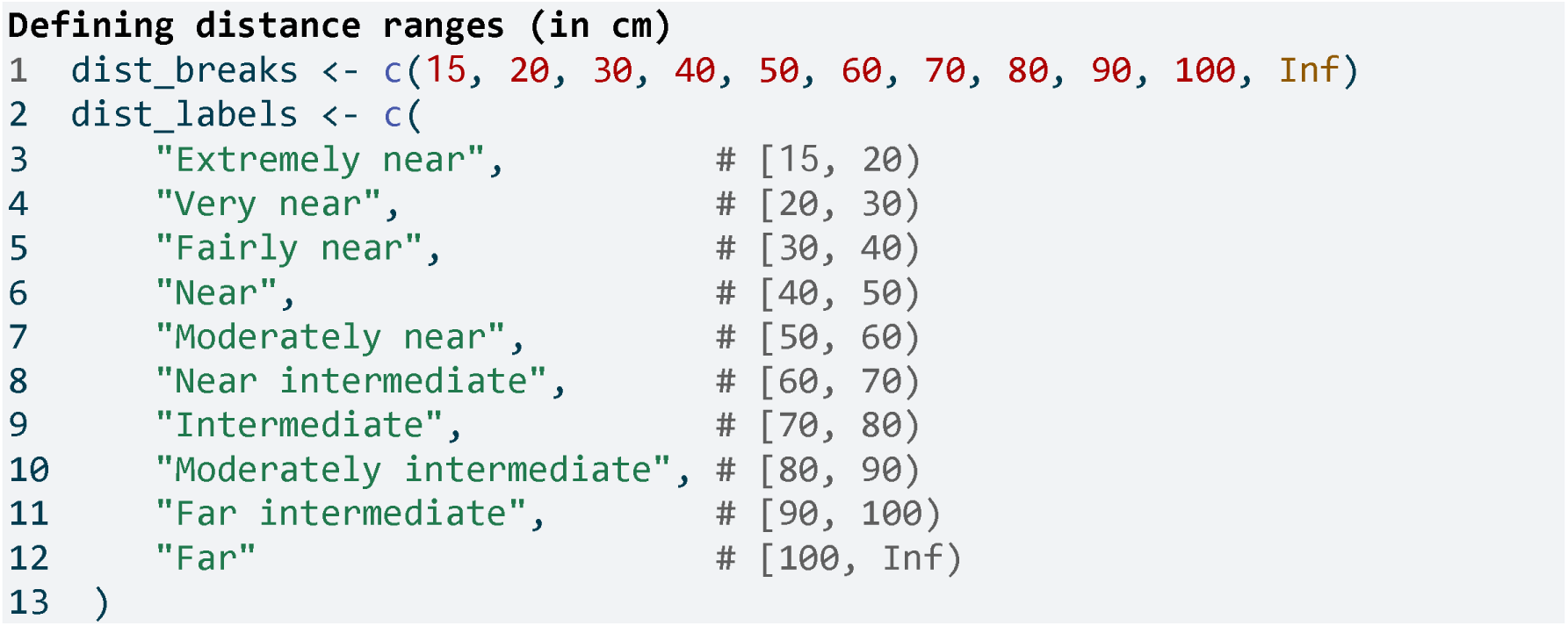

Now we cut the distance data into these ranges and compute the daily duration spent in each range:

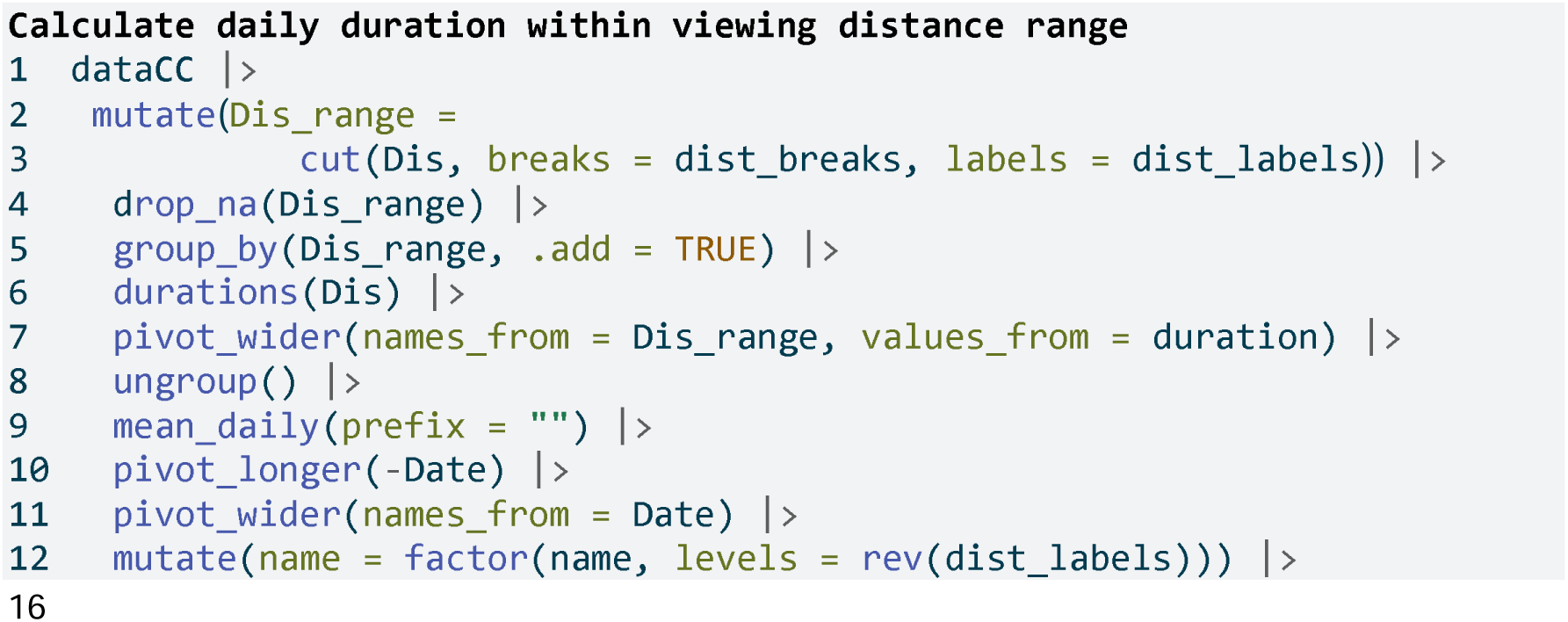
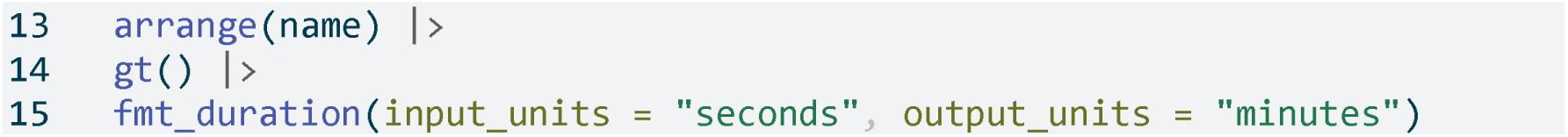

**Line 2-3** Categorize distances

**Line 4** Remove intervals with no data

**Line 5** Group by distance range (in addition to the date)

**Line 6** Duration per range per day

**Line 7** Pivot data from long to wide format (ranges as columns)

**Line 15** Convert seconds to minutes

To visualize this, **Figure 3** illustrates the relative proportion of time spent in each distance range:

**Figure 3:**
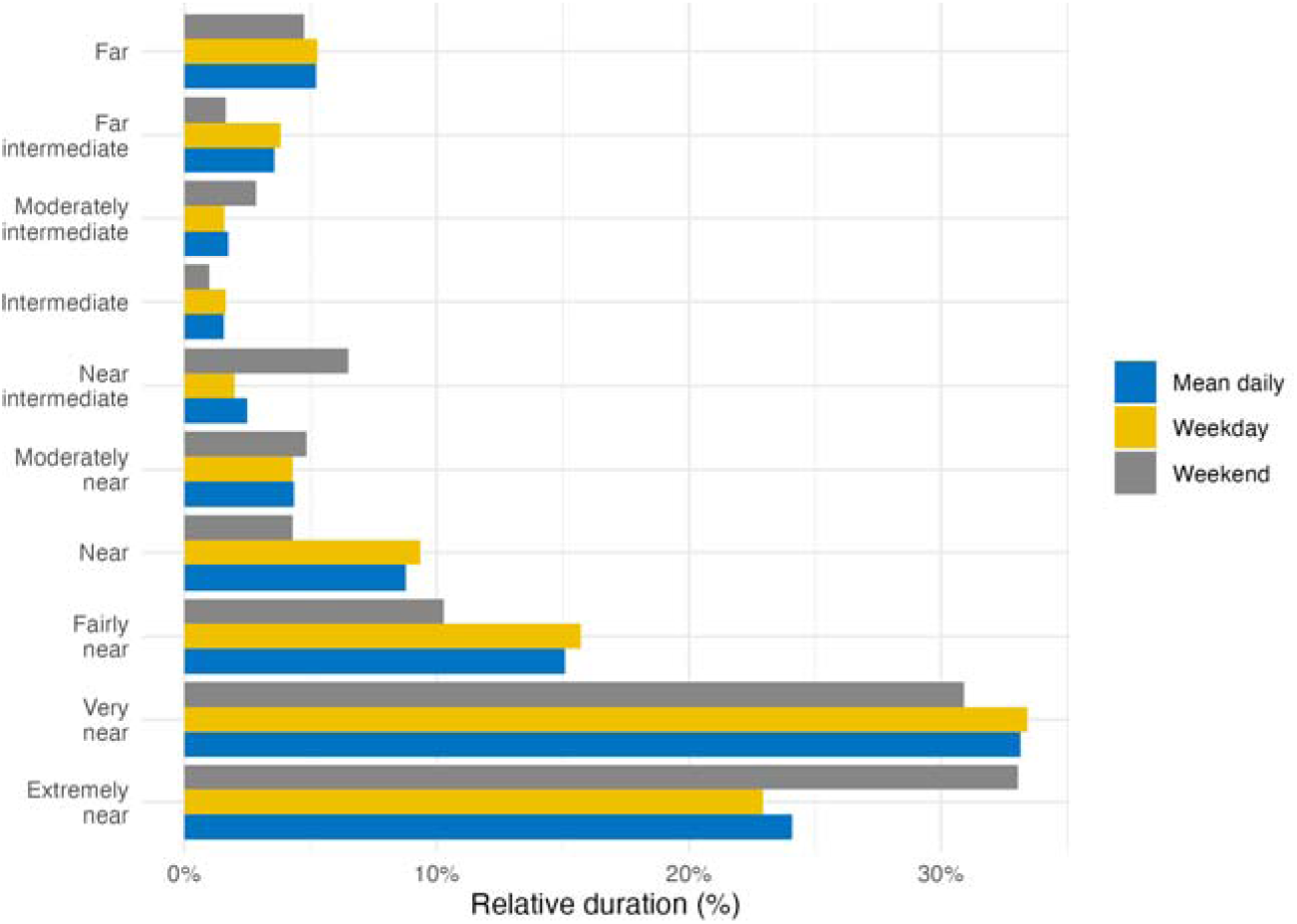
Percentage of total time spent in each viewing distance range for an average day (mean daily), average weekday, or weekend

### Frequency of continuous near work

Continuous near-work can be understood as sustained viewing within a near distance for some minimum duration, allowing only brief interruptions. We use LightLogR’s cluster function to identify episodes of continuous near work. Here, we define a near-work episode as viewing distance between 20 cm and 60 cm that lasts at least 30 minutes, with interruptions of up to 1 minute allowed (meaning short breaks ≤1 min do not end the episode). Using <u>extract_clusters()</u> with those parameters, we count how many such episodes occur per day.

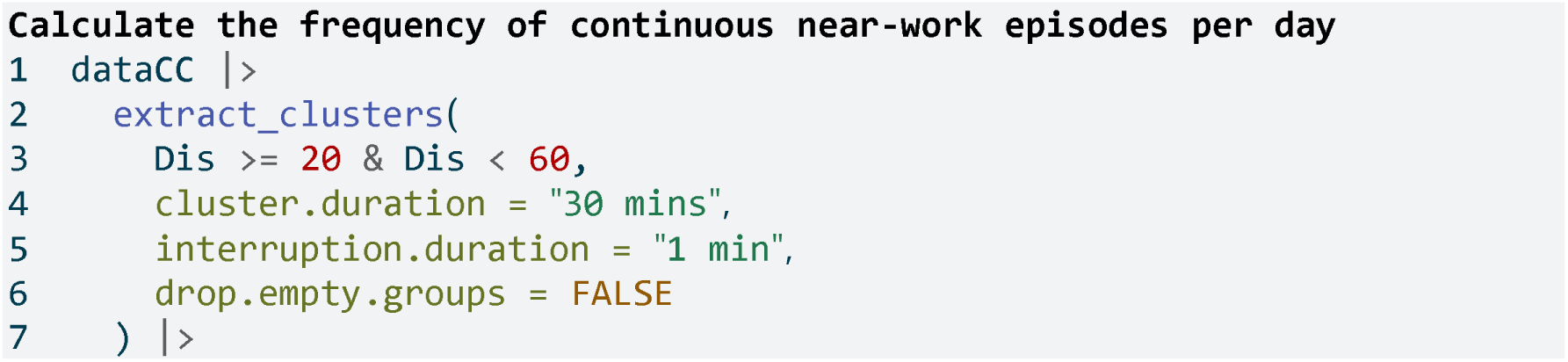
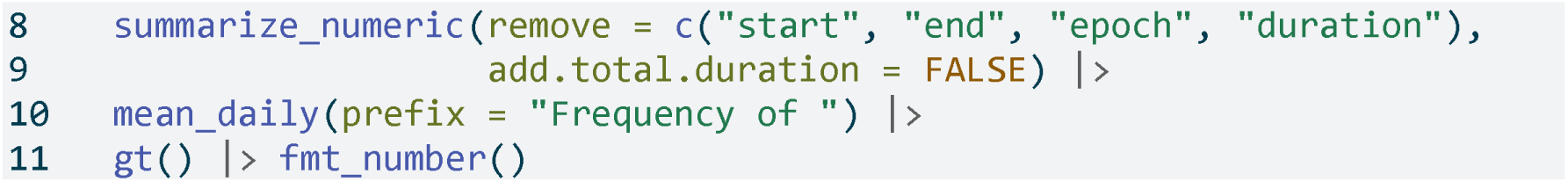

Table 6 summarizes the average frequency of continuous near-work episodes per day, and **Figure 4** provides an example visualization of these episodes on the distance time series.

**Figure 4:**
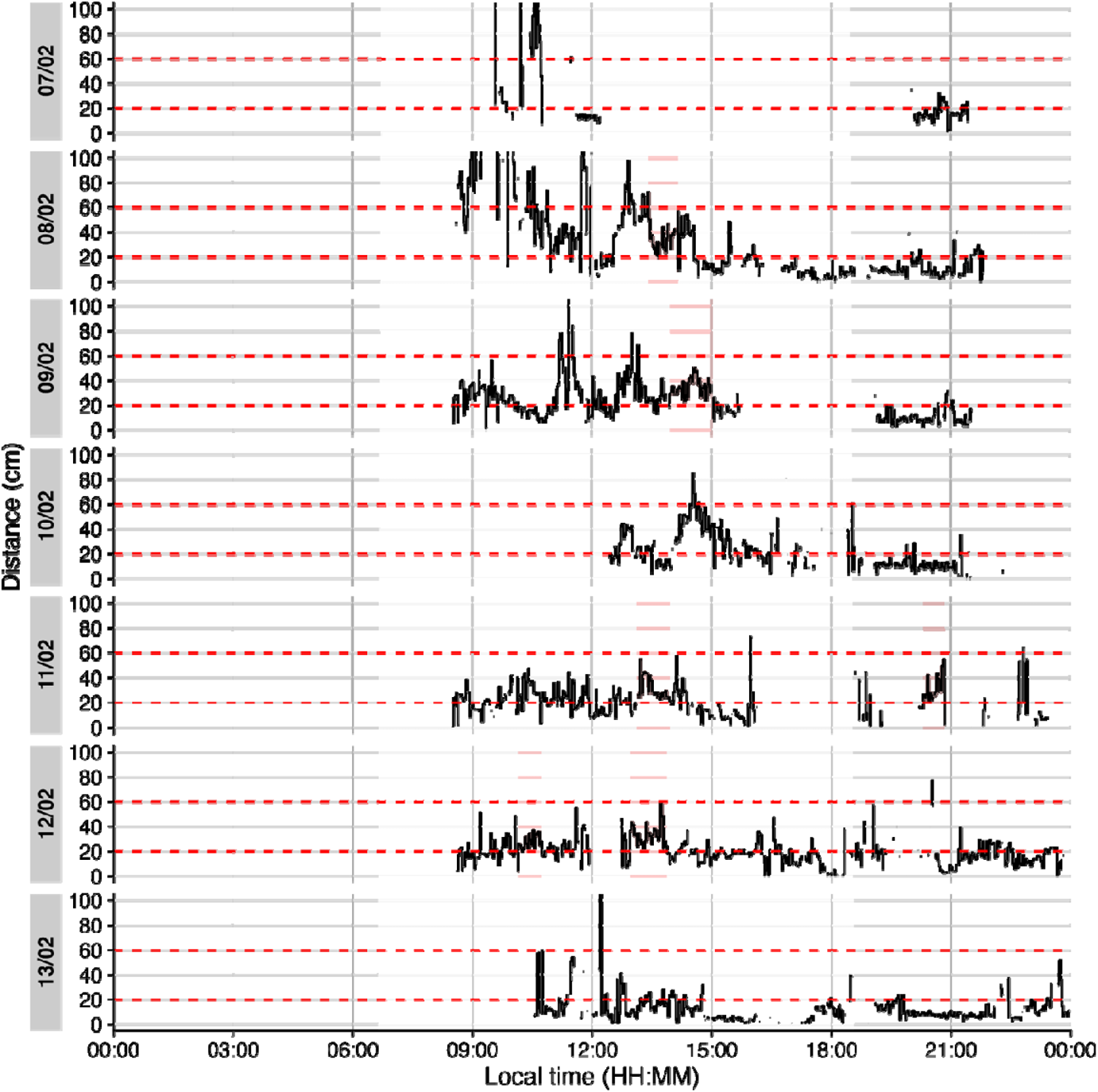
Example of continuous near-work episodes. Red shaded areas indicate periods of continuous near work (20–60 cm for ≥30 min, allowing ≤1 min interruptions). Black trace is viewing distance over time of day; red dashed lines mark the 20 cm and 60 cm boundaries. Grey shaded areas show nighttime between civil dusk and civil dawn

**Table 6:** Frequency of continuous near-work episodes per day

|  |  |
| --- | --- |
| Mean daily | 0.86 |
| Weekday | 1.20 |
| Weekend | 0 |

**Line 3** Condition: near-work distance

**Line 4** Minimum duration of a continuous episode

**Line 5** Maximum gap allowed within an episode

**Line 6** Keep days with zero episodes in output

**Line 9 C**ount number of episodes per day

**Line 10** Compute daily mean frequency

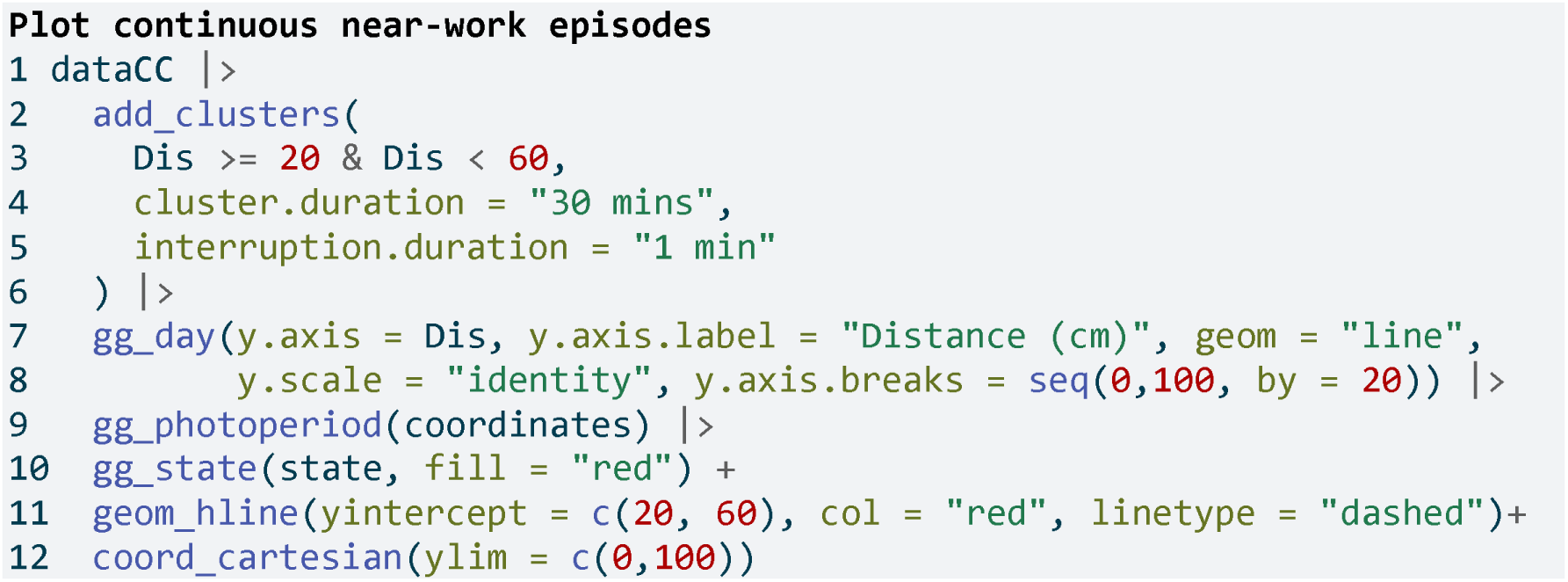

**Lines 1-6** As in code cell above

**Line 9** Add photoperiod information

**Line 10** Add state bands

### Near-work episodes

Beyond frequency, we can characterize near-work episodes by their duration and typical viewing distance. This section extracts all near-work episodes (using a 5-second minimum duration to capture more routine near-work bouts) and summarizes three aspects:

1. frequency (count of episodes per day),
2. average duration of episodes, and
3. average distance during those episodes. These results are combined in **Table 7**.

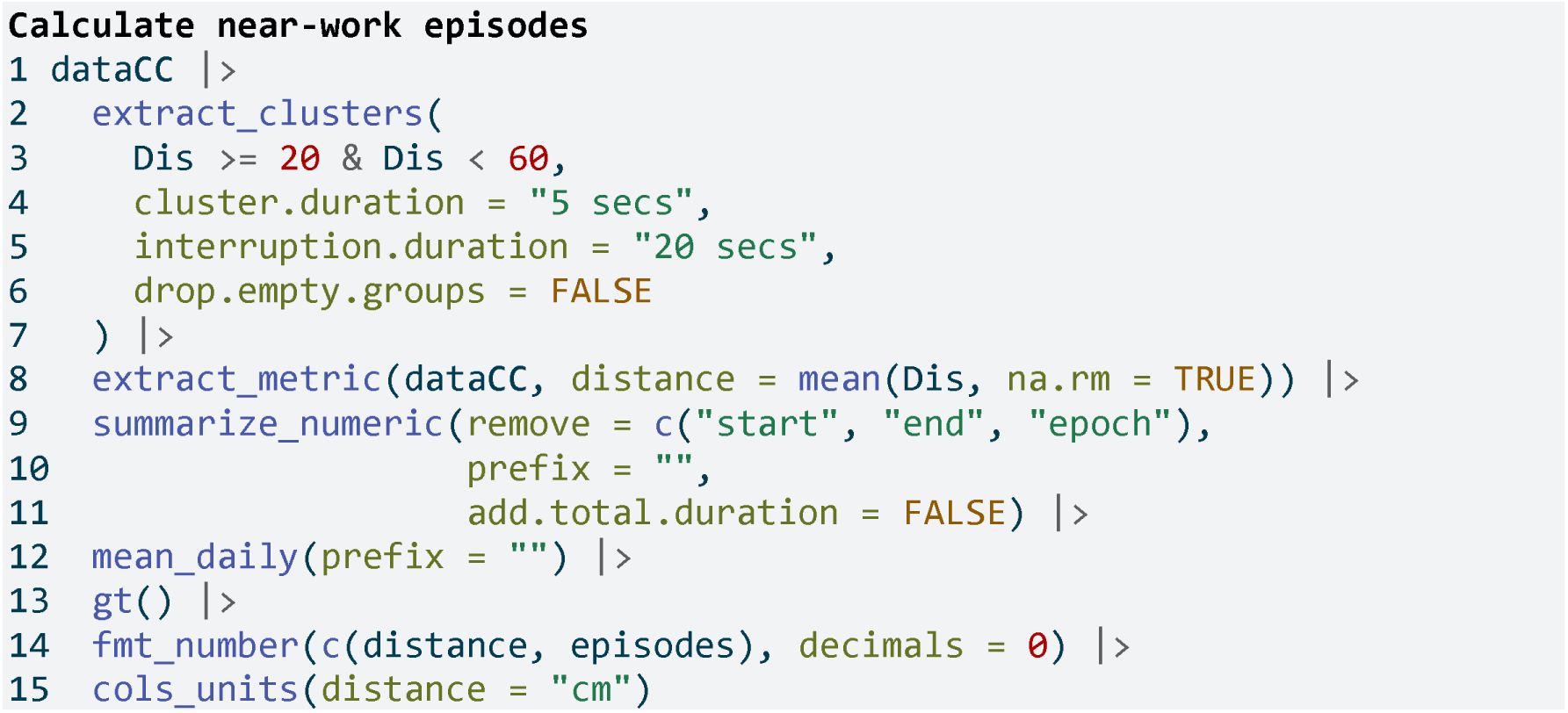

**Table 7:** Near-work episodes: frequency, mean duration, and mean viewing distance

|  | duration | distance, cm | episodes |
| --- | --- | --- | --- |
| Mean daily | 233s (~3.88 minutes) | 32 | 57 |
| Weekday | 284s (~4.73 minutes) | 32 | 64 |
| Weekend | 104s (~1.73 minutes) | 32 | 40 |

**Line 4** Minimal duration to count as an episode (set to interval level of dataCC)

**Line 8** Calculate mean distance during each episode

**Lines 9-11** Calculate averages for all numeric columns per group

**Line 12** Daily averages for each metric

**Lines 13-15** Table generation

In the code cell above, extract_metric(…, distance = mean(Dis, …)) computes the mean viewing distance during each episode, and the subsequent summarize_numeric and mean_daily steps derive daily averages of episode count, duration, and distance.

### Visual breaks

Visual breaks as defined in this article, require a minimum break-length, and the previous episode is important. This leads to a two step process, where we first extract instances of Distance above 100 cm for at least 20 seconds, before we filter for a previous duration of at maximum 20 minutes. **Table 8** provides the daily frequency of visual breaks and **Figure 5** shows when these occur.

**Figure 5:**
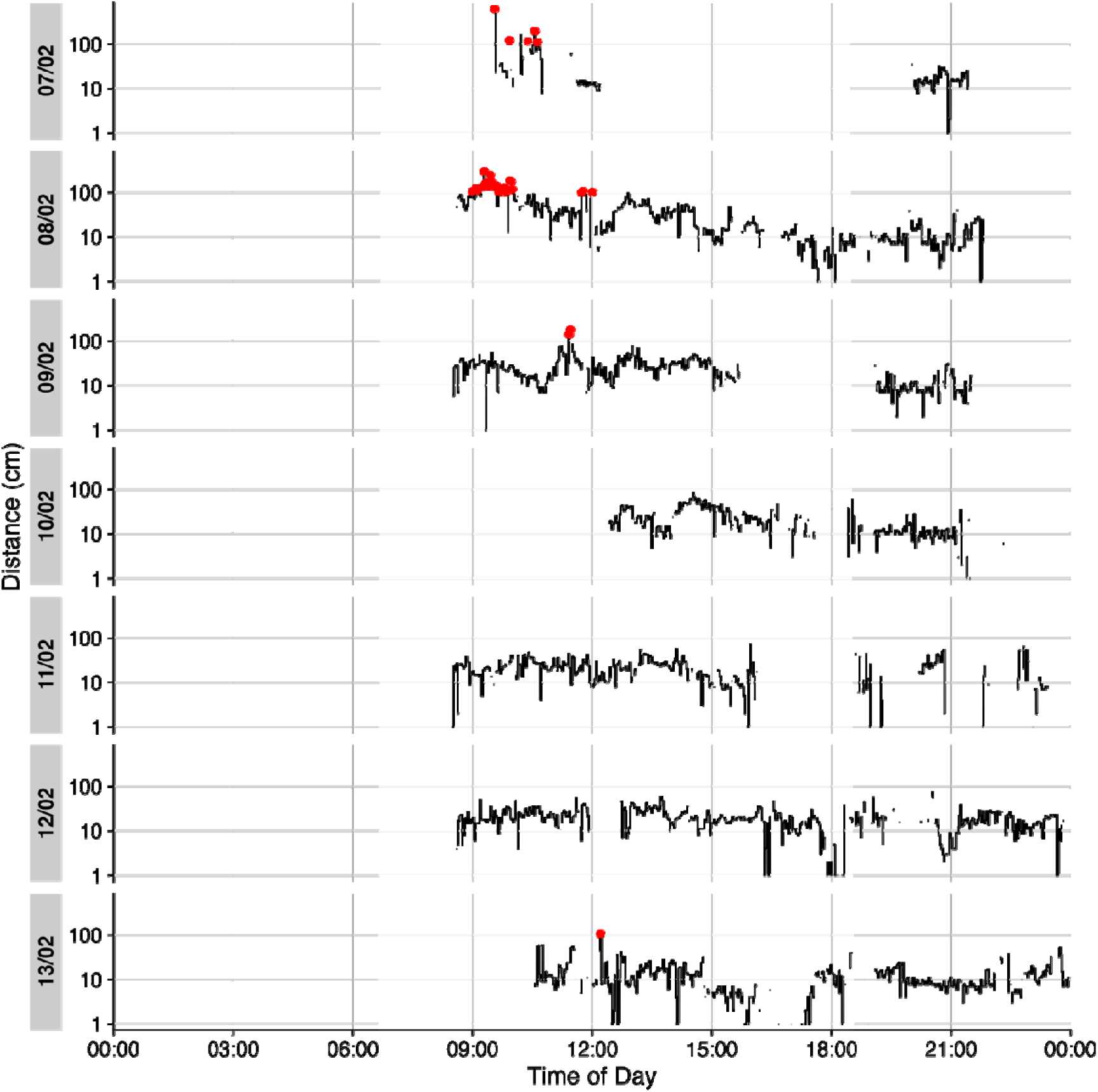
Plot of visual breaks (red dots). Black traces show distance measurement data. Grey shaded areas show nighttime between civil dusk and civil dawn (photoperiod)

**Table 8:**
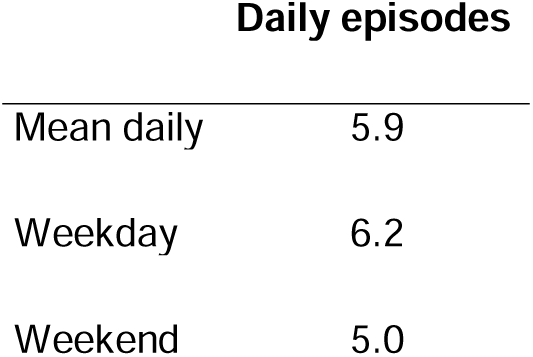
Frequency of visual breaks (n per day)

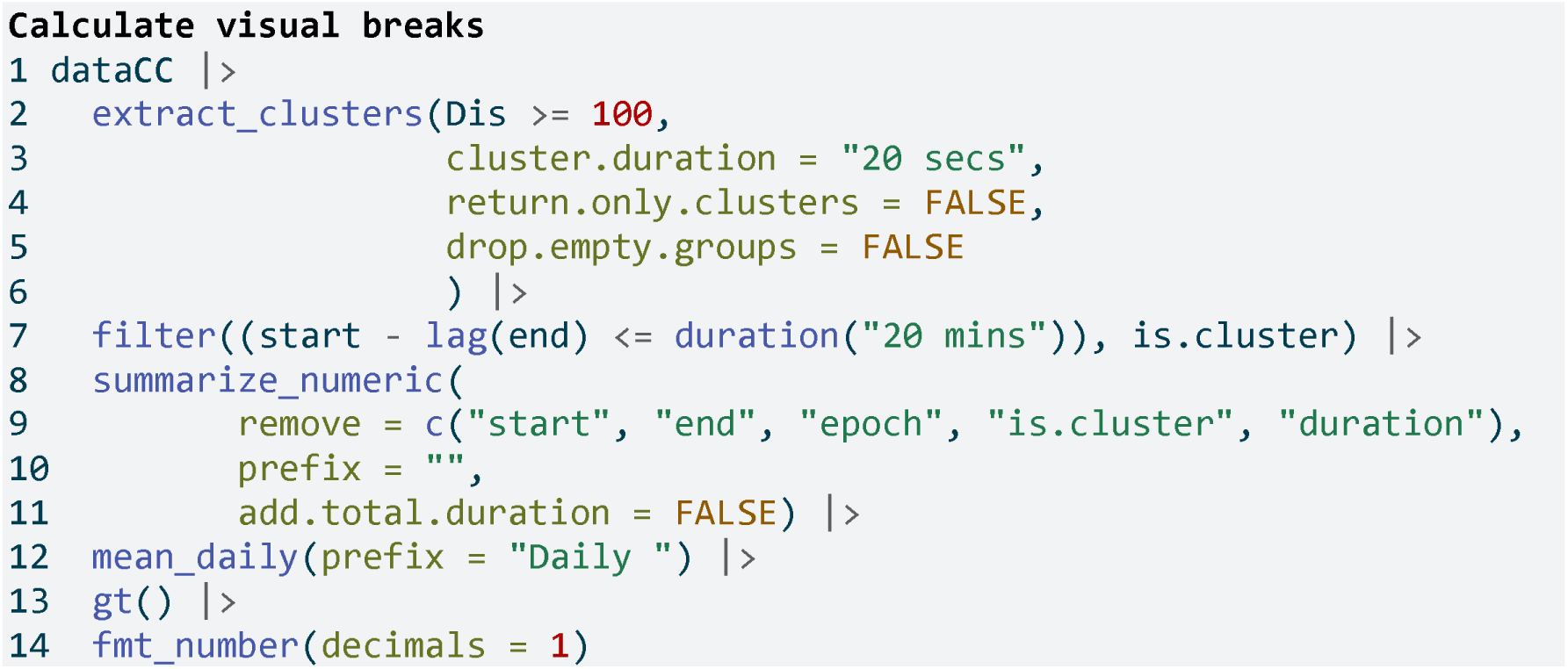

**Line 2** Define the condition, greater 100 cm away

**Line 3** Define the minimum duration

**Line 4** Return non-clusters as well

**Line 5** Keep all days, even without clusters

**Line 7** Return only clusters with previous episode lengths of maximum 20 minutes

**Lines 8-11** Count the number of episodes

**Line 12** Calculate daily means

**Lines 13-14** Table generation

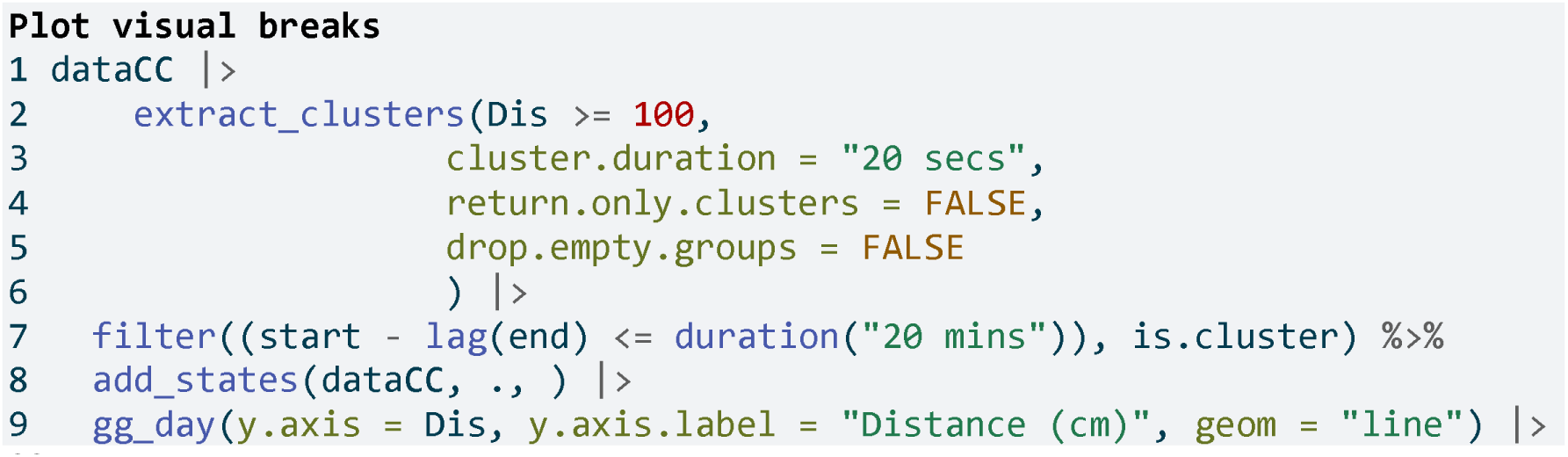
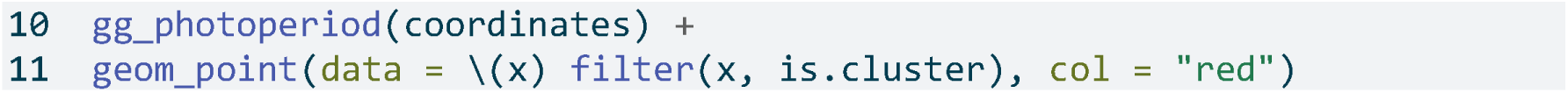

**Lines 1-7** As in the code cell above

**Line 8** Add the resulting states

**Line 10** Add photoperiod information to the plot

### Distance with spatial distribution

The Clouclip device outputs a singular measure for distance, while the visual environment in natural conditions contains many distances, depending on the solid angle and direction of the measurement. A device like the VEET increases the spatial resolution of these measurements, allowing for more in-depth analyses of the size and position of an object within the field of view. In the case of the VEET, data are collected from an 8x8 measurement grid, spanning 52° vertically and 41° horizontally. **Figure 6** shows sample observations from six different days at the same time.

**Figure 6:**
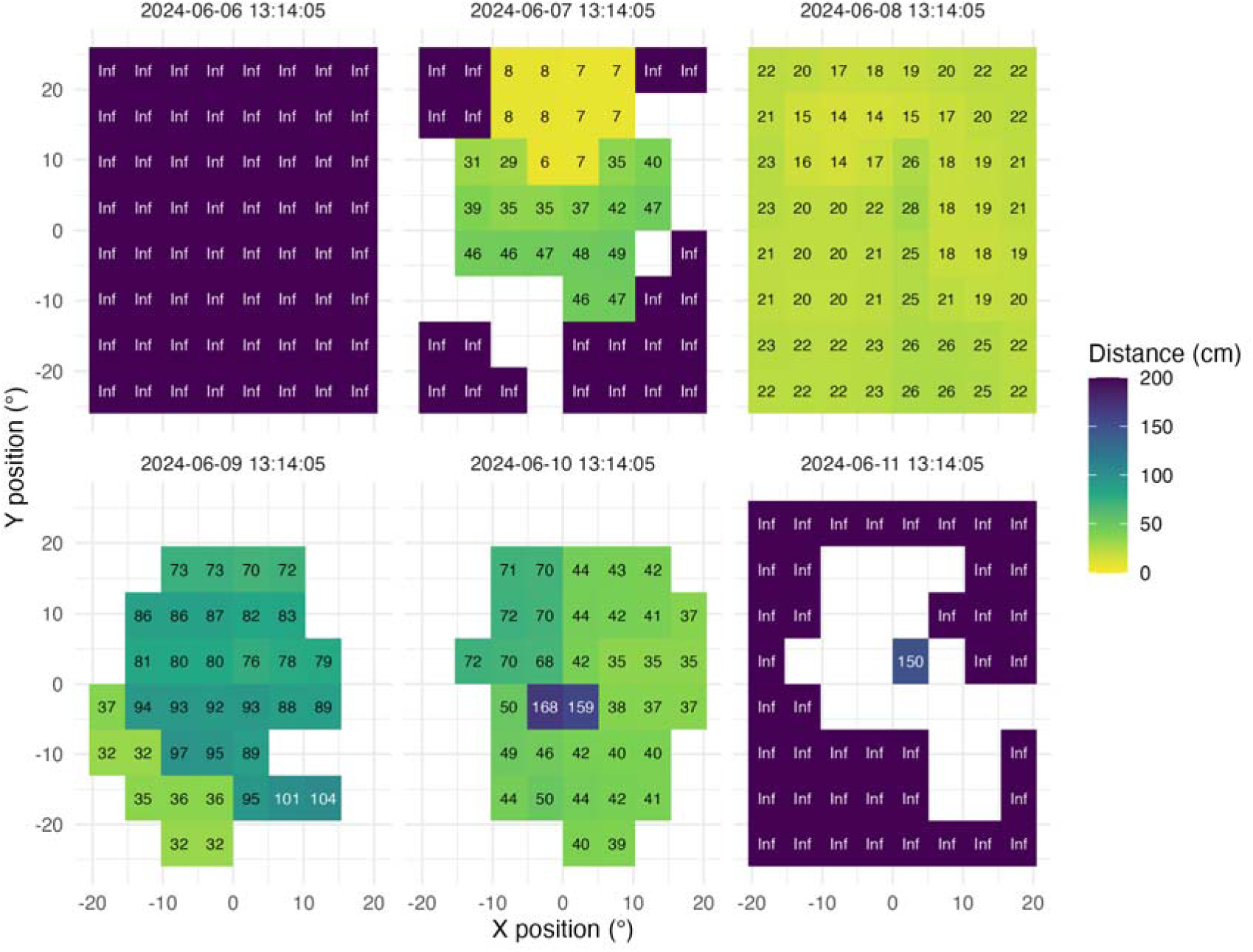
Example observations of the measurement grid at 1:14 p.m. for each measurement day. Text values show distance in cm. Empty grid points show values with low confidence. Zero-distance values were replaced with infinite distance and plotted despite low confidence.

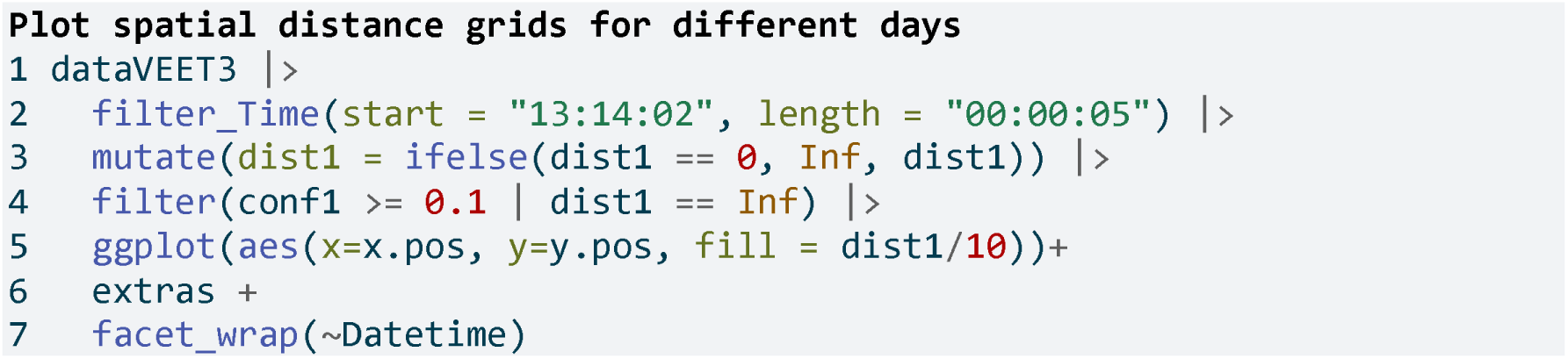

**Line 2** Choose a particular observation

**Line 3** Replace 0 distances with infinity

**Line 4** Remove data that has less than 10% confidence

**Line 5-6** Plot the data and add the plot partials from the code cell above

**Lines 7** Show one plot per day

To use these distance data in the framework shown above for the Clouclip device, a sensible method to condense the data has to be applied. There are many ways how a spatially resolved distance measure could be utilized for analysis:

- Where in the field of view are objects in close range?
- How large are near objects in the field of view?
- How varied are distances within the field of view?
- How close are objects / is viewing distance in a region of interest within the field of view?

Possible methods include:

- average across all (high confidence) distance values within the grid
- closest (high confidence) distance within the grid
- (high confidence) values at or around a given grid position, e.g., ±10 degrees around the central view (0°)

Many more options are available based on the spatial dataset, e.g., condensation rules based on the number of points in the grid with a given condition, or the variation within the grid.

We will demonstrate these three methods for a single day (2024-06-10), all leading to a data structure akin to the Clouclip, i.e., to be used for further calculation of visual experience metrics.

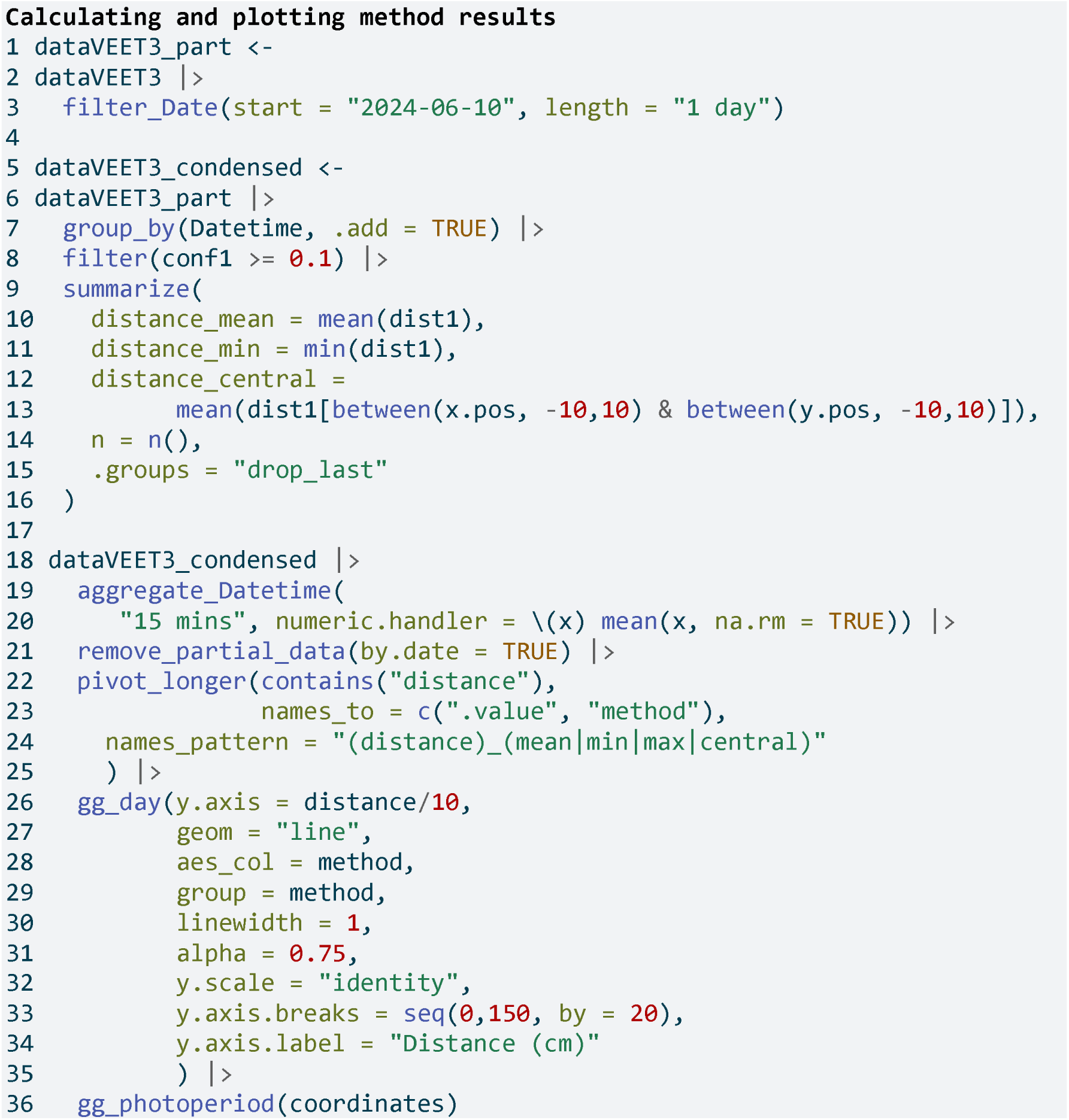

**Lines 1-3** Filter one day

**Line 7** Group additionally by every observation

**Line 8** Remove data with low confidence **Line 10** Average across all distance values **Line 11** Closest across all distance values **Line 12-13** Central distance

**Line 14** Number of (valid) grid points

**Line 19-20** Aggregate to 15 minute data

**Line 21** Remove data points that fall exactly on midnight of the following day

**Lines 22-25** Pivoting the method results from wide to long for plotting

**Lines 26-36** Setting up the plot for distance.

As can be seen in **Figure 7**, while the overall pattern is similar regardless of the used method, there are notable differences between the methods, which will consequently affect downstream analyses. Most importantly, the process of condensation has to be well documented and reproducible, as shown above. Any of these data could now be used to calculate the frequency of continuous near work, visual breaks, or near-work episodes as described above.

**Figure 7:**
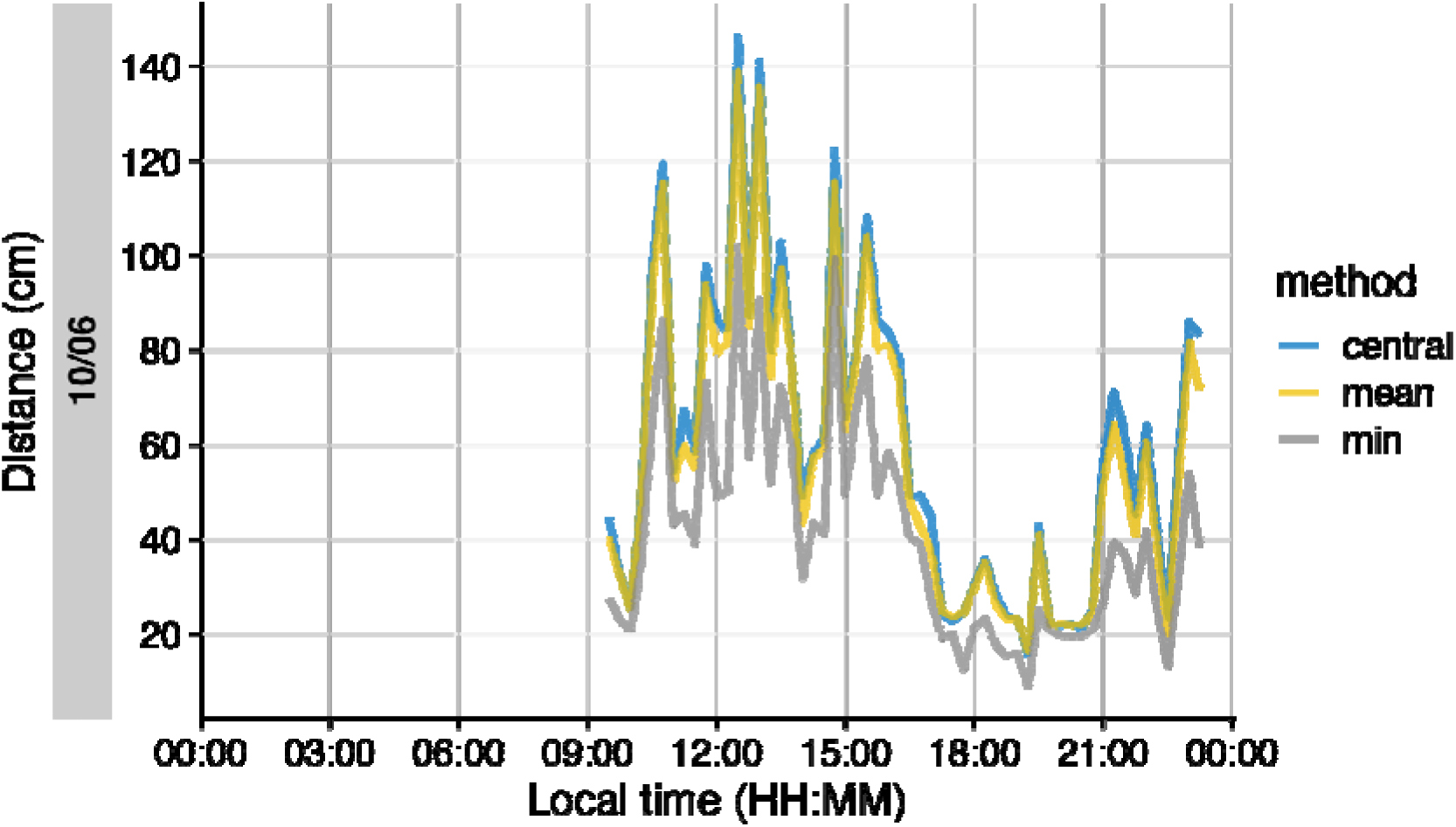
Comparison of condensation methods for spatial grid of distance measurements. The lines represent an average across all data points (yellow), the minimum distance (grey), or the central 10° (blue). Data points with confidence less than 10% were removed prior to calculation.

### Light

The Clouclip illuminance data in our example cover indoor environments and are thus comparatively low, which would make certain daylight exposure summaries trivial or not meaningful. To better illustrate light exposure metrics, we turn to a different dataset, this one taken from the VEET device’s illuminance data, which capture a broader range of lighting conditions (though both device types are able to capture broadly the same range of illuminance). We import the VEET ambient light data (already preprocessed to have regular 5-second intervals as described above) and briefly examine its distribution.

**Illuminance distribution**: The illuminance values from the Clouclip were comparatively low (**Figure 8)**, while the VEET (**Figure 9)** data include outdoor exposures up to several thousand lux. The contrast is evident from comparing histograms of the two datasets’ lux values (Clouclip vs. VEET), where the main peak is similarly positioned between 10 and 100 lx, but the tails differ. The VEET illuminance histogram (see **Figure 9**) shows a heavily skewed distribution with a considerable number of zero lx values (indicating intervals of complete darkness or the sensor being covered) and a long tail extending to very high lux values. Such zero-inflated and skewed data are common in wearable light measurements (Zauner, Guidolin, and Spitschan, 2025).

**Figure 8:**
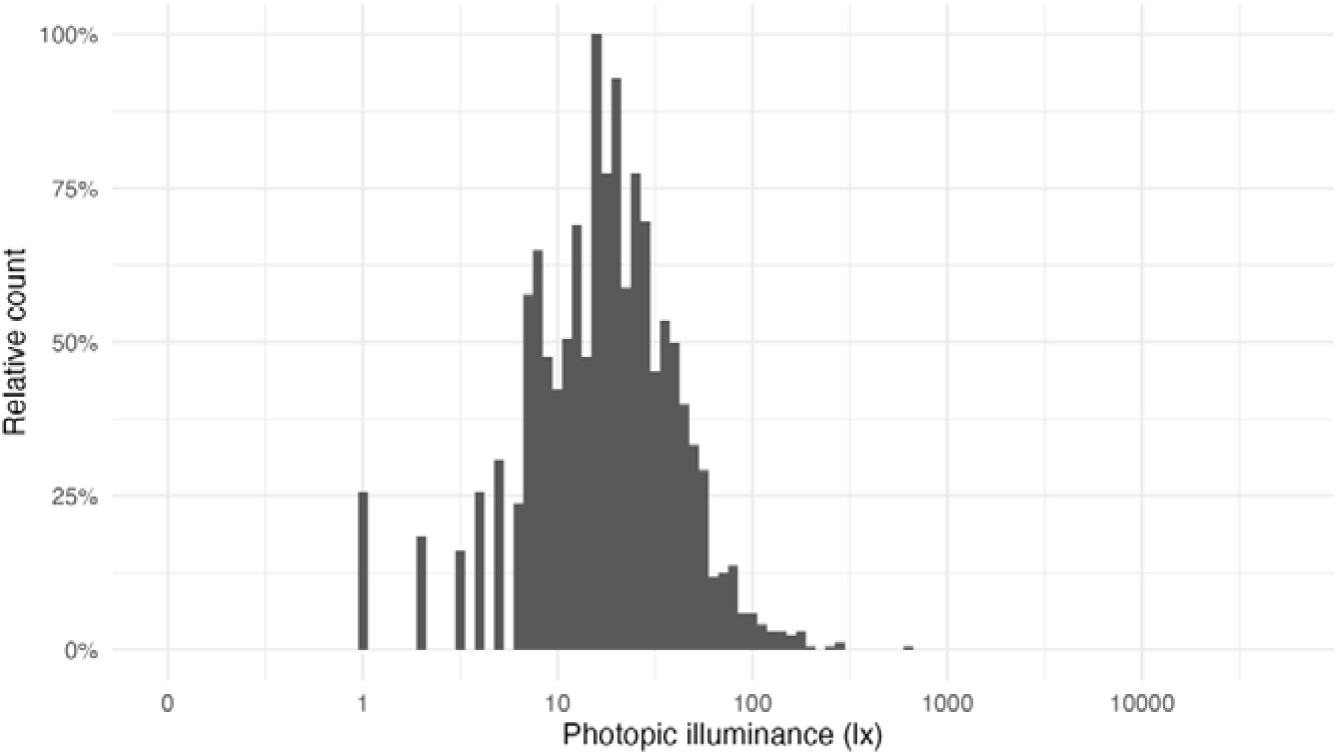
Histogram of illuminance values from the Clouclip dataset (5-second data). The values are typical of indoor conditions.

**Figure 9:**
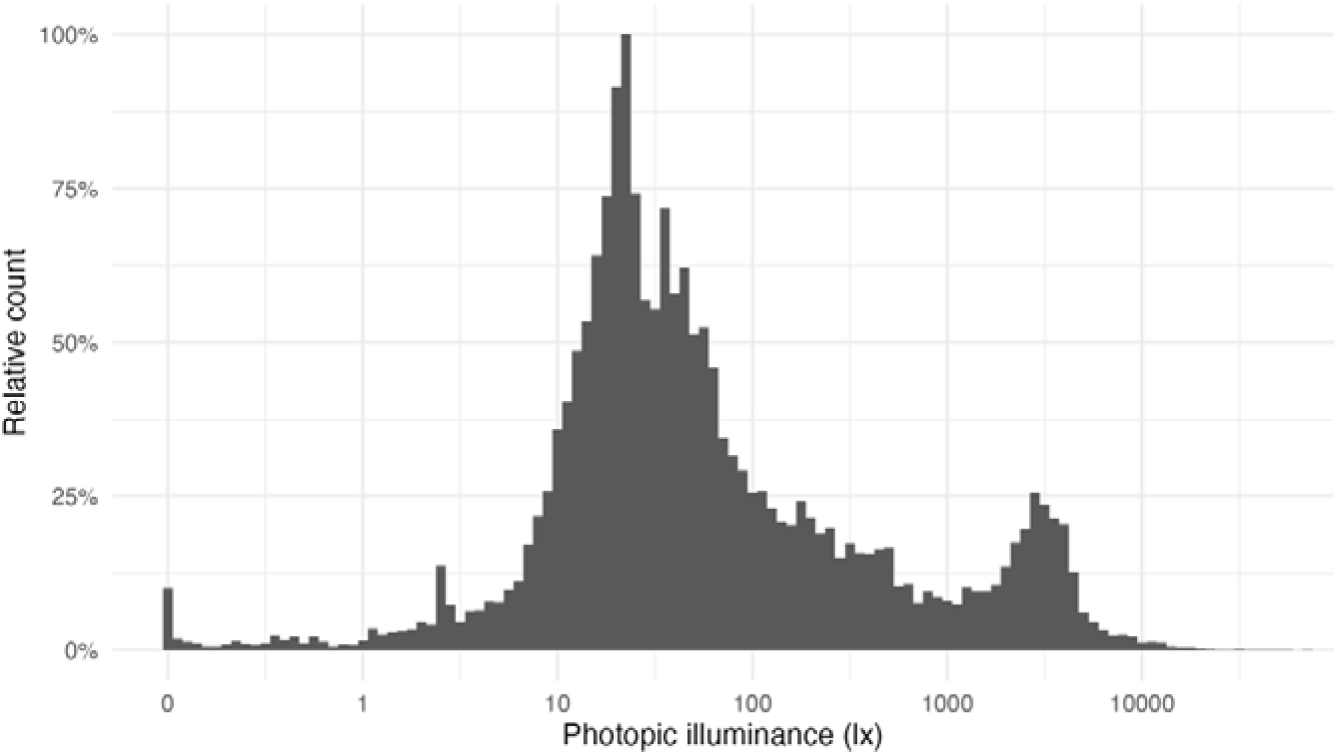
Histogram of illuminance values from the VEET dataset (aggregated to 5 s). Note the logarithmic x-axis: the distribution is highly skewed with many low values (including zeros) and a long tail of high lux readings. Outdoor light exposures in bright conditions are distinguishable around 10^3^ to 10^4^ lx.

After confirming that the VEET data cover a broad dynamic range of lighting, we proceed with calculating light exposure metrics. (The VEET data had been cleaned for gaps and irregularities as described earlier, and non-wear times were removed; see <u>Supplement 1</u> for the details.)

### Average light exposure

A basic metric is the average illuminance over the day. **Table 9** shows the mean illuminance (in lux) for weekdays, weekends, and the overall daily mean, calculated directly from the raw lux values.

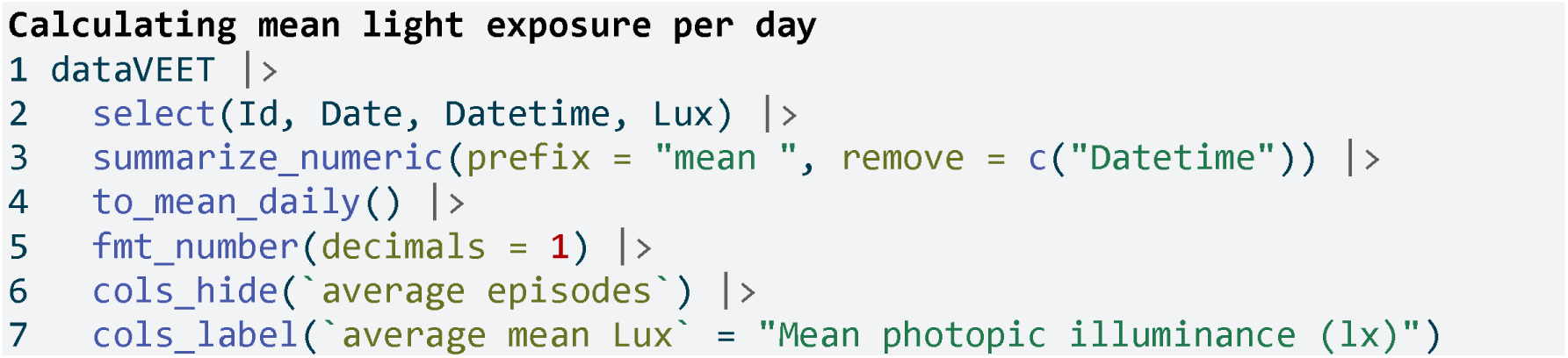

**Table 9:**
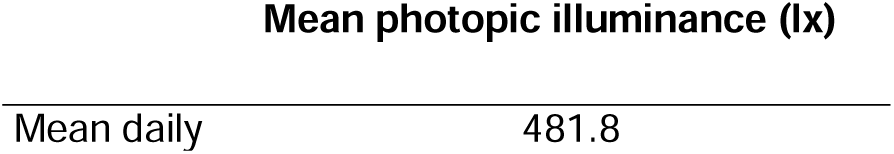
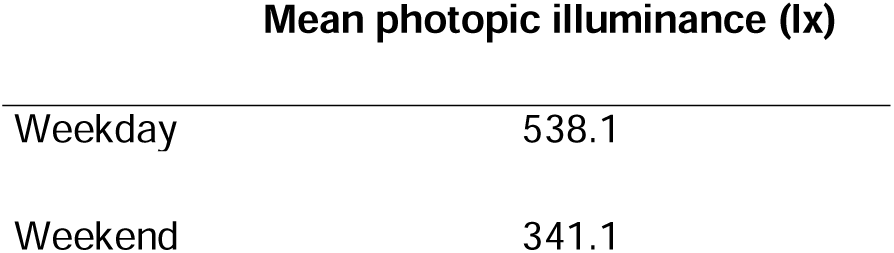
Mean light exposure (illuminance) per day

However, because illuminance data tend to be extremely skewed and contain many zero values (periods of darkness), the arithmetic mean can be <u>misleading</u>. A common approach is to apply a logarithmic transform to illuminance before averaging, which down-weights extreme values and accounts for the multiplicative nature of light intensity effects. LightLogR provides helper functions log_zero_inflated() and its inverse exp_zero_inflated() to handle log-transformation when zeros are present (by adding a small offset before log, and back-transforming after averaging). Using this approach, we recompute the daily mean illuminance. The results in **Table 10** show that the log-transformed mean (back-transformed to lux) is much lower, reflecting the fact that for much of the time illuminance was near zero.

**Table 10:** Mean light exposure per day (after logarithmic transformation to account for zero inflation and skewness)

|  |  |
| --- | --- |
| Mean daily | 57.0 |
| Weekday | 70.1 |
| Weekend | 33.9 |

This transformed mean is often more representative of typical exposure for skewed data.

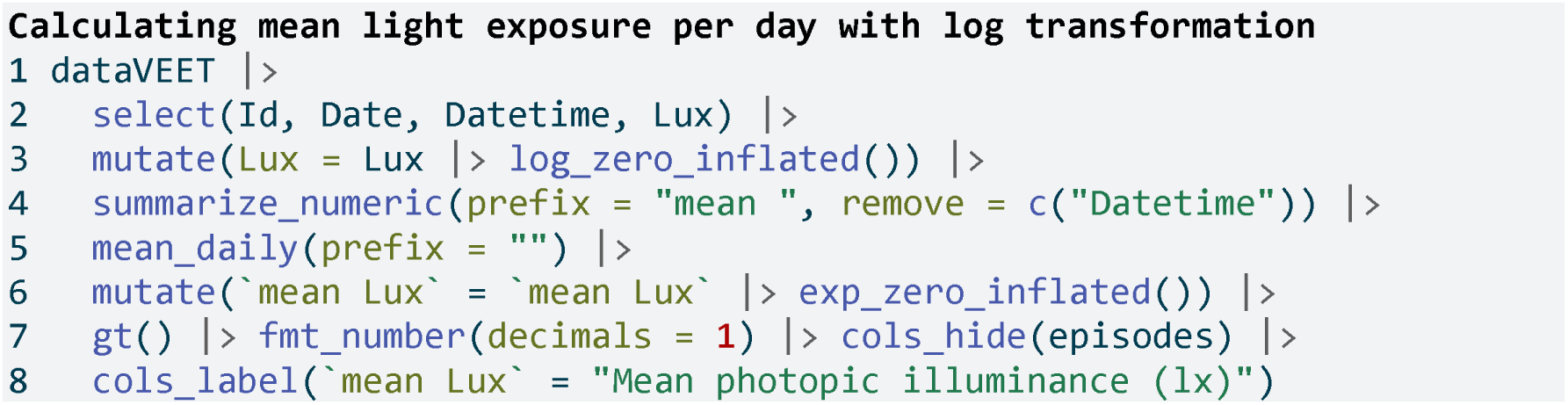

**Line 3** Log transform with zero handling (base 10)

**Line 5** Calculate daily mean of log-lux

**Line 6** Back-transform to lux

### Duration in high-light (outdoor) conditions

Another important metric is the amount of time spent under bright light, often used as a proxy for outdoor exposure. We define thresholds corresponding to outdoor light levels (e.g. 1000 lx and above). Here, we categorize each 5-second interval of illuminance into bands: Outdoor bright (≥1000 lx), Outdoor very bright (≥2000 lx), and Outdoor extremely bright (≥3000 lx). We then sum the duration in each category per day.

While daylight levels can far exceed the recorded light levels, those are usually recorded with direct sunlight and without obstruction. Under normal viewing conditions, at eye level, and avoiding glare, daylight levels of a few thousand lux are at the higher end of the distribution (Murukesu, Zauner, and Spitschan 2025).

Figure 9 shows a bimodal distribution, with the right mode representing outdoor lighting conditions. In a 2023 review of light dosimeters to investigate the light-myopia relationship (Hönekopp and Weigelt 2023), 1000 lx was the predominant cutoff value to distinguish indoor vs. outdoor environments. It is not, however, without critique, and both other thresholds (Patterson Gentile et al. 2025) and classification methods are proposed (Tabandeh and Spitschan 2025).

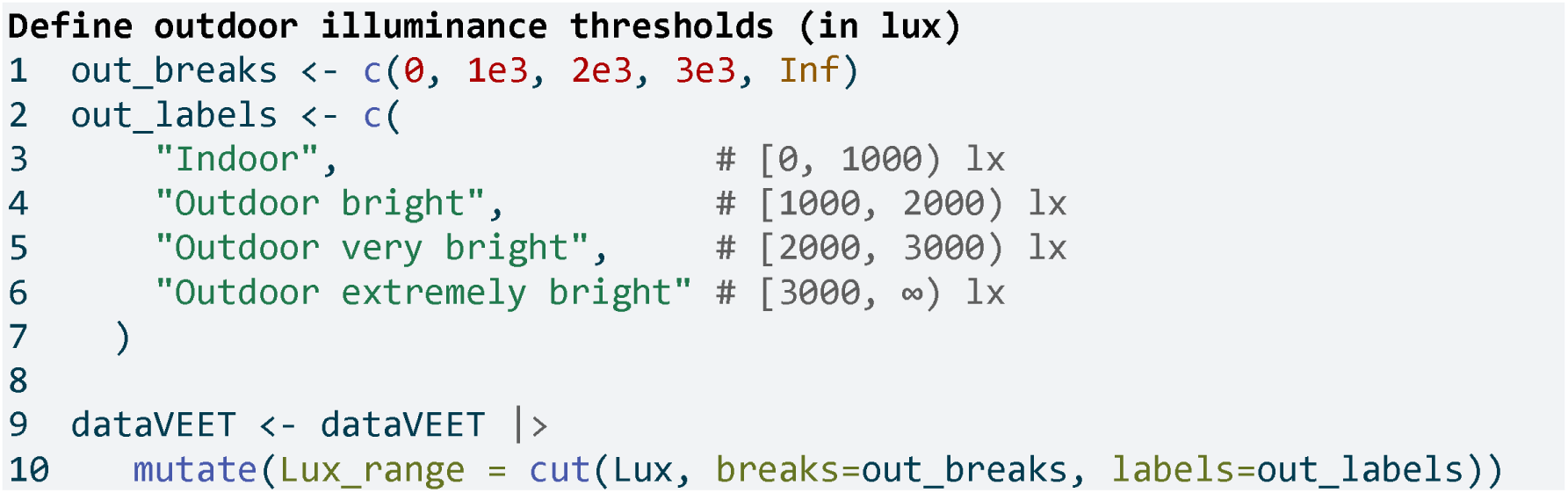

Now we compute the mean daily duration spent in each of these outdoor light ranges (**Table 11**):

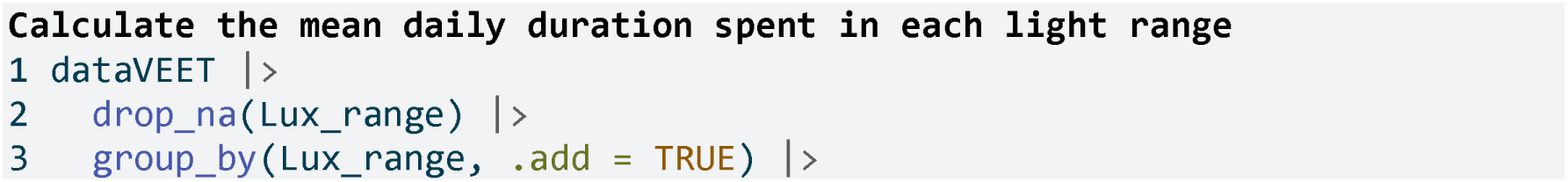
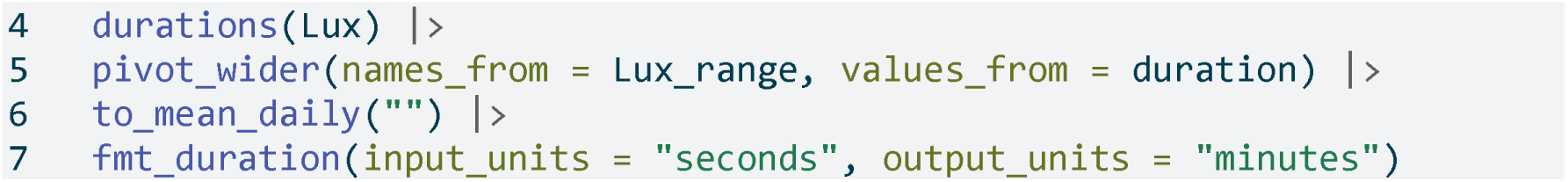

**Table 11:**
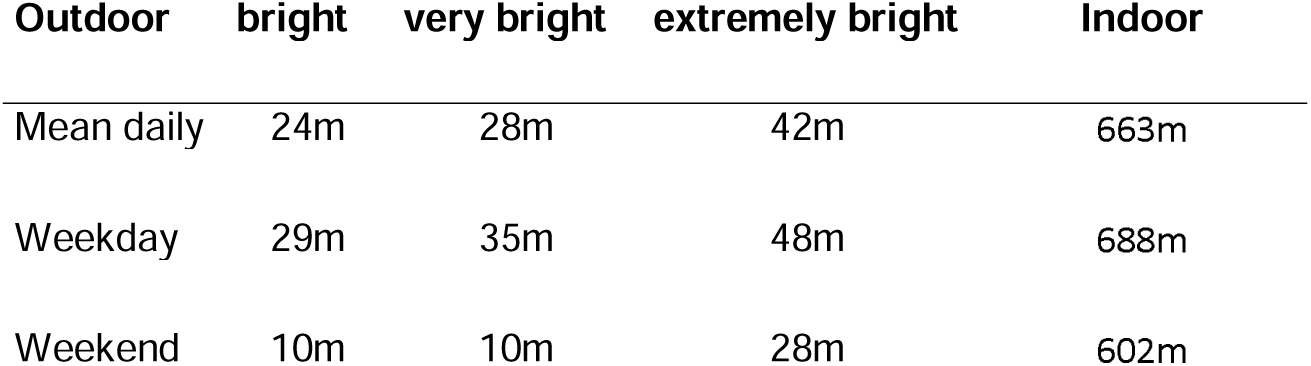
Average daily duration in outdoor-equivalent light conditions (column order is different from code output)

It is also informative to visualize when these high-light conditions occurred.

Figure 10 shows a timeline plot with periods of outdoor-level illuminance highlighted in color. In this example, violet denotes ≥1000 lx, green ≥2000 lx, and yellow ≥3000 lx. Grey shading indicates nighttime (from civil dusk to dawn) for context.

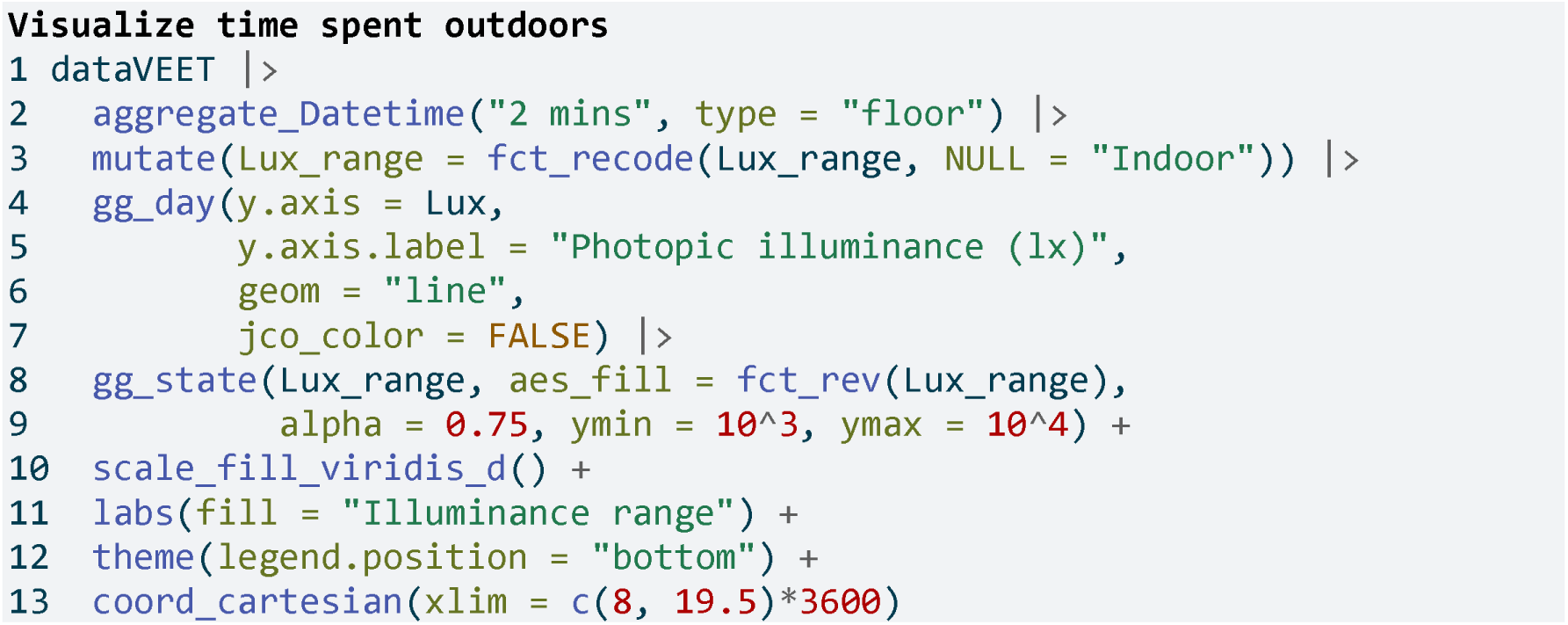

**Figure 10:**
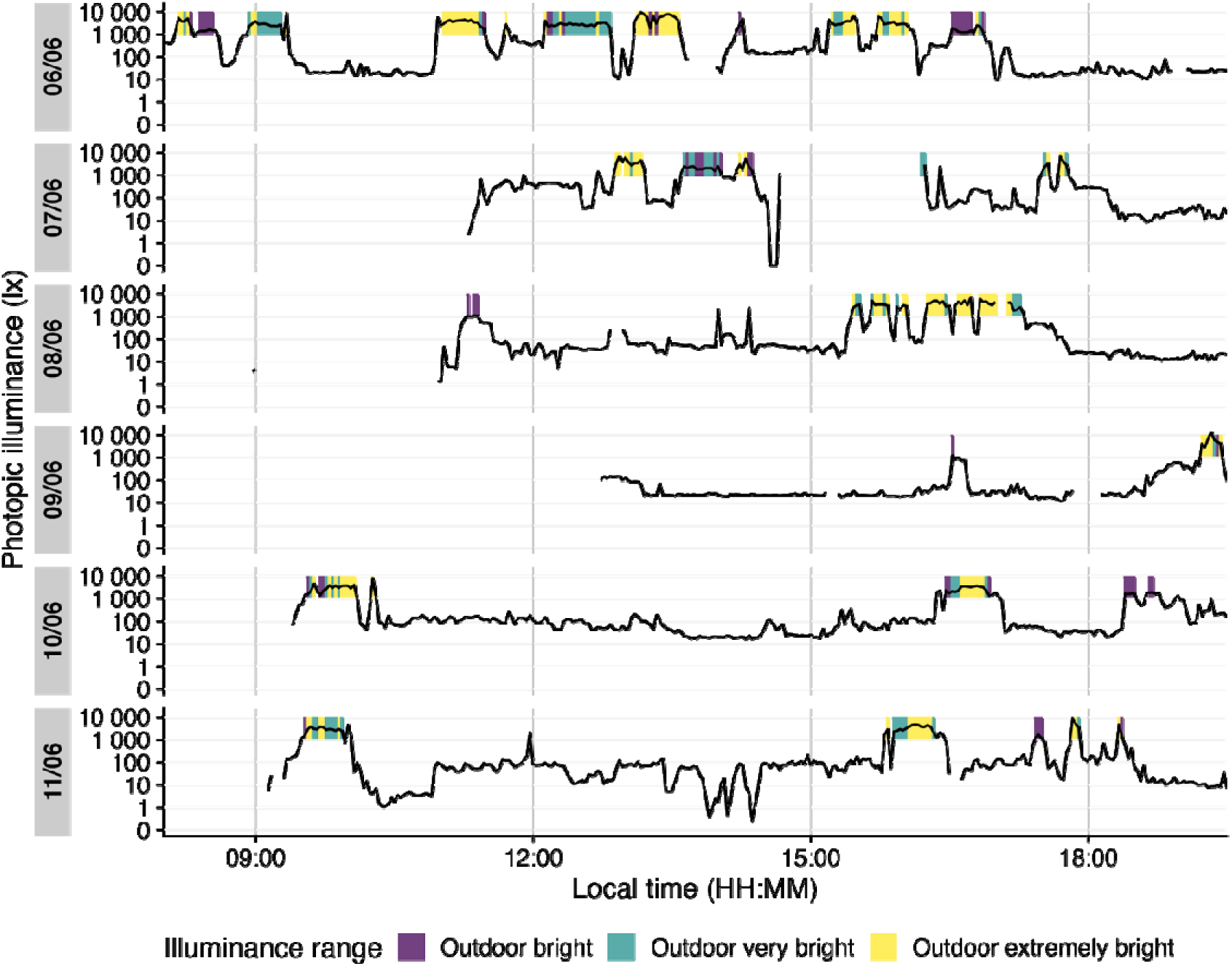
Outdoor light exposure over time with 2-minute interval. Colored bands indicate periods when illuminance exceeded outdoor thresholds for at least half of each interval: violet for ≥1000 lx, green for ≥2000 lx, and yellow for ≥3000 lx. Grey shaded regions denote night (from civil dusk to dawn).

**Line 2** Aggregating data to 5-minute bins

**Line 3** Removing the indoor condition

**Lines 4-7** Setting up the basic plot

**Lines 8-9** Adding state information on the illuminance ranges

**Line 13** Setting the x-axis limits to cover daytime hours

### Frequency of transitions from indoor to outdoor light

We next consider how often the subject moved from an indoor light environment to an outdoor-equivalent environment. We operationally define an “outdoor transition” as a change from <1000 lx to ≥1000 lx. Using the cleaned VEET data, we extract all instances where illuminance crosses that threshold from below to above.

**Table 12** shows the average number of such transitions per day. Note that if data are recorded at a fine temporal resolution (5 s here), very brief excursions above 1000 lx could count as transitions and inflate this number. Indeed, the initial count is fairly high, reflecting fleeting spikes above 1000 lx that might not represent meaningful outdoor exposures.

**Table 12:** Average daily count of transitions from indoor (<1000 lx) to outdoor (≥1000 lx) lighting when looking at 5-second epochs

|  | mean epoch | episodes |
| --- | --- | --- |
| Mean daily | 5s | 64 |
| Weekday | 5s | 72 |
| Weekend | 5s | 46 |

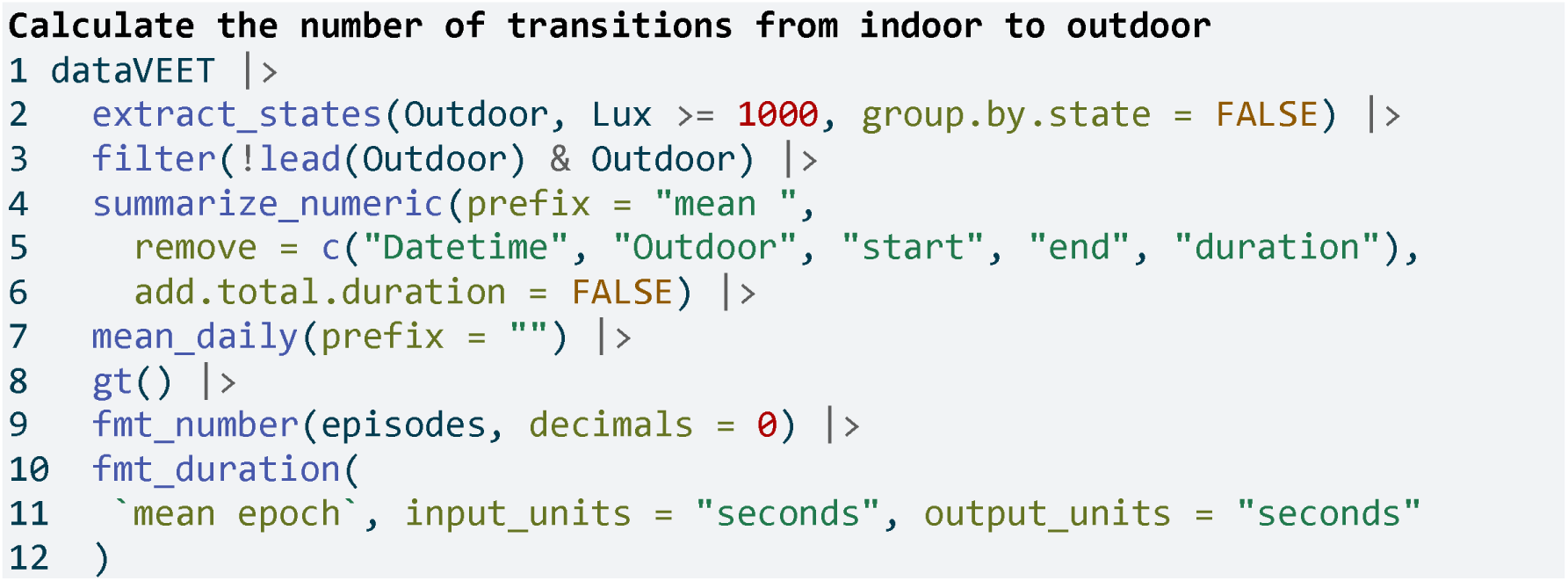

**Line 2** Label each interval as Outdoor (Lux≥1000) or not

**Line 3** Find instances where the previous interval was “indoor” and current is “outdoor”

To obtain a more meaningful measure, we can require that the outdoor state persists for some minimum duration to count as a true transition (filtering out momentary fluctuations around the 1000 lx mark). For example, we can require that once ≥1000 lx is reached, it continues for at least 5 minutes (allowing short interruptions up to 20 s). **Table 13** applies this criterion, resulting in a lower, more plausible transition count.

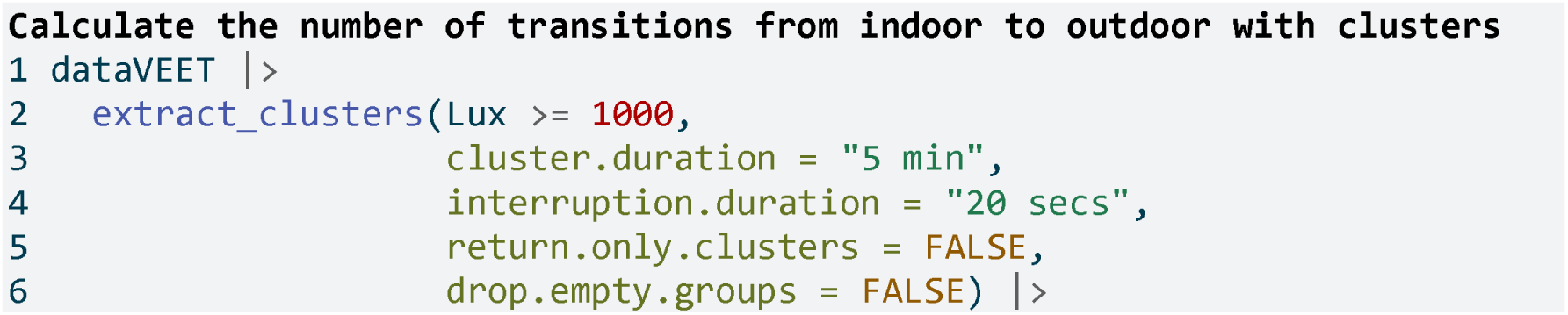

**Table 13:**
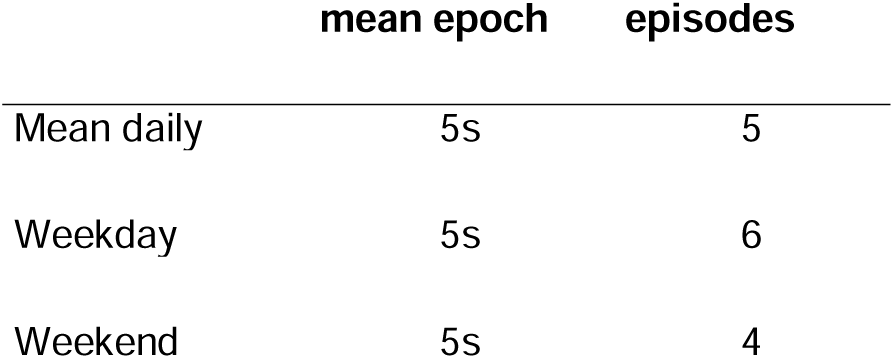
Daily indoor-to-outdoor transition count (requiring ≥5 min duration of ≥1000 lx to count)

|  | mean epoch | epsisodes |
| --- | --- | --- |
| Mean daily | 5s | 5 |
| Weekday | 5s | 6 |
| Weekend | 5s | 4 |

### Longest sustained bright-light period

The final light exposure metric we illustrate is the longest continuous period above a certain illuminance threshold (often termed longest period above threshold, e.g. PAT1000 for 1000 lx). This gives us a sense of the longest outdoor exposure in a day. Along with it, one might report the total duration above that threshold in the day (TAT1000). While we could derive these from the earlier analyses, LightLogR provides dedicated <u>metric</u> functions for such calculations, which can compute multiple related metrics at once.

Using the function <u>period_above_threshold()</u> for PAT and <u>duration_above_threshold()</u> for TAT, we calculate both metrics for the 1000 lx threshold. **Table 14** shows the mean of these metrics across days (i.e., average longest bright period and average total bright time per day).

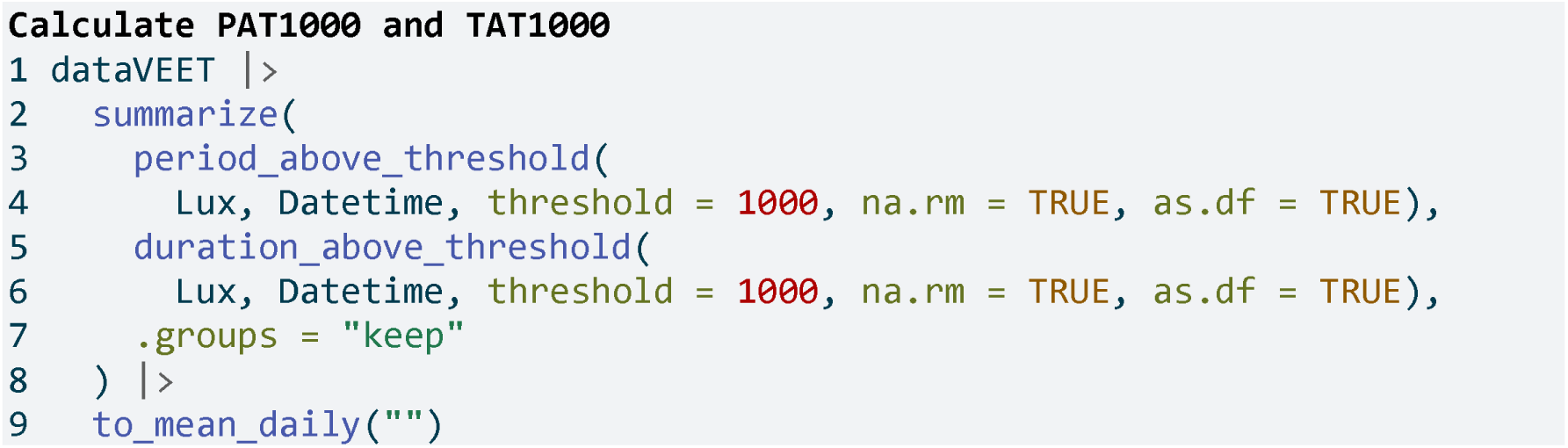
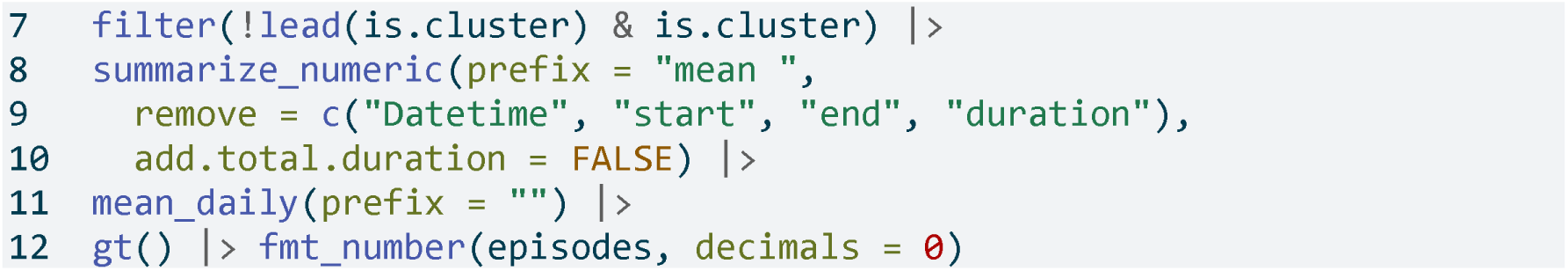

**Table 14:** Longest period and total duration above 1000 lx (PAT1000 and TAT1000)

|  | period above 1000 | duration above 1000 |
| --- | --- | --- |
| Mean daily | 1208s (~20.13 minutes) | 5703s (~1.58 hours) |
| Weekday | 1469s (~24.48 minutes) | 6815s (~1.89 hours) |
| Weekend | 555s (~9.25 minutes) | 2922s (~48.7 minutes) |

**Table 15:**
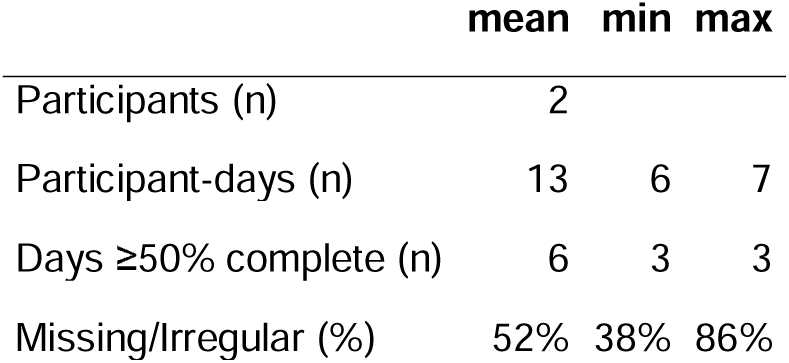
Overview of the merged dataset

|  | mean | min | max |
| --- | --- | --- | --- |
| Participants (n) | 2 |  |  |
| Participant-days (n) | 13 | 6 | 7 |
| Days ≥50% complete (n) | 6 | 3 | 3 |
| Missing/Irregular (%) | 52% | 38% | 86% |

### Merging data streams

Note that while imports from different devices can be <u>merged</u>, devices differ in their sensors, electronics, housing or diffuser form factors, and on-device data-processing pipelines. All of these factors affect the comparability of measurements, even when devices output the same variable (e.g., illuminance or distance). If data from different devices with the same measurement variable are to be merged, the corresponding variable names should be standardized beforehand - for example, renaming Lux and LIGHT to illuminance. If we wanted to analyse the VEET data together with the Clouclip data, for example, we would not have to rename anything, as both carry their illuminance measurements in the variable Lux. The following example shows how the combination of datasets would lead to a combined dataset, and how that would affect analysis outcomes. It is the responsibility of the researcher to perform device calibration and/or checks for a similar measurement fidelity.

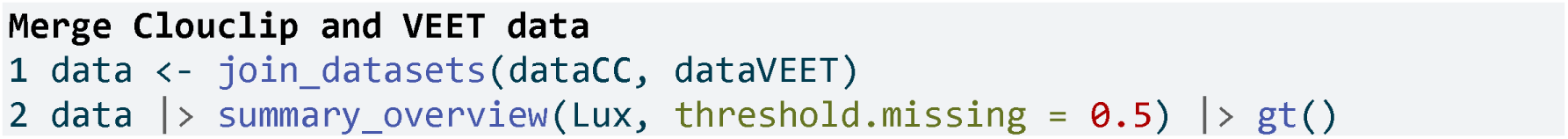

We will reuse the example from Section 3.2.1, but instead of one participant, we now have data from two devices and participants

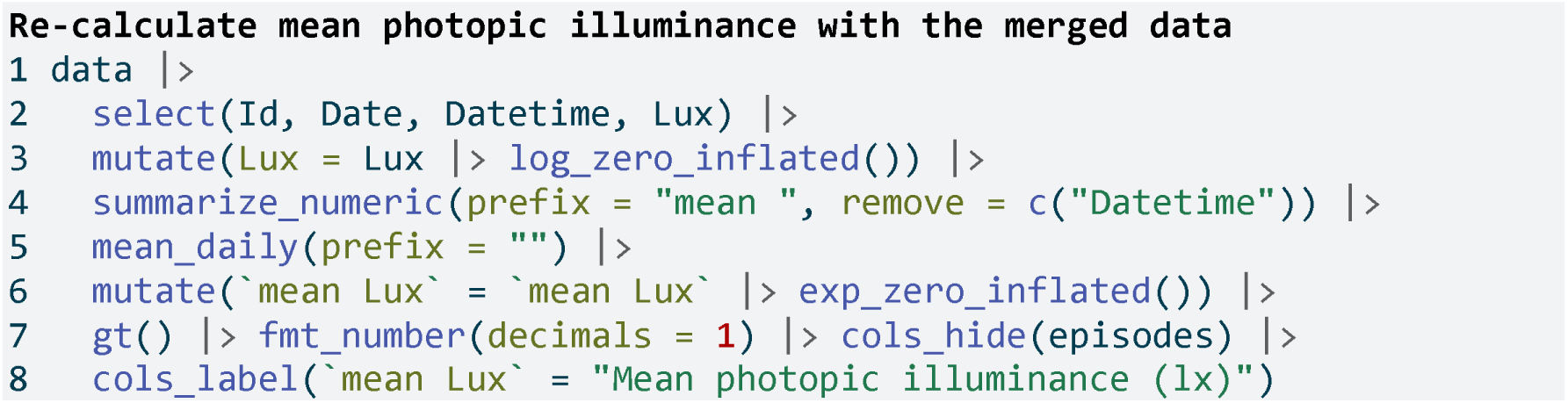

**Line 1** Instead of dataVEET we now supply the merged data object

**Lines 2-8** Verbatim from Section 3.2.1

### Spectrum

The VEET device’s spectral sensor provides multimodal data beyond simple lux values, but it requires reconstruction of the actual light spectrum from raw sensor counts. We processed the spectral sensor data in order to compute two example spectrum-based metrics. Detailed data import, normalization, and spectral reconstruction steps are given in <u>Supplement 1</u>; here we present the resulting metrics. Briefly, the VEET’s spectral sensor recorded counts in nine wavelength bands (roughly 415 nm to 910 nm), plus a Dark, a Clear, and a flicker detection channel^1^. After normalizing by sensor gain and applying the calibration matrix, we obtained an estimated spectral irradiance distribution for each 5-minute interval in the recording. With these reconstructed spectra, we can derive novel metrics that consider spectral content of the light.

Spectrum-based metrics in wearable data are relatively new and less established compared to distance or broadband light metrics. The following examples illustrate potential uses of spectral data in a theoretical sense, which can be adapted as needed for specific research questions.

### Ratio of short- vs. long-wavelength light

Our first spectral metric is the ratio of short-wavelength light to long-wavelength light, which is relevant, for example, in assessing the blue-light content of exposure. We define “short” wavelengths as 400–500 nm and “long” as 600–700 nm (which are not standardized thresholds and can be freely adjusted). Using the list-column of spectra in our dataset, we integrate each spectrum over these ranges (using spectral_integration()), and then compute the ratio short/long for each time interval. We then summarize these ratios per day.

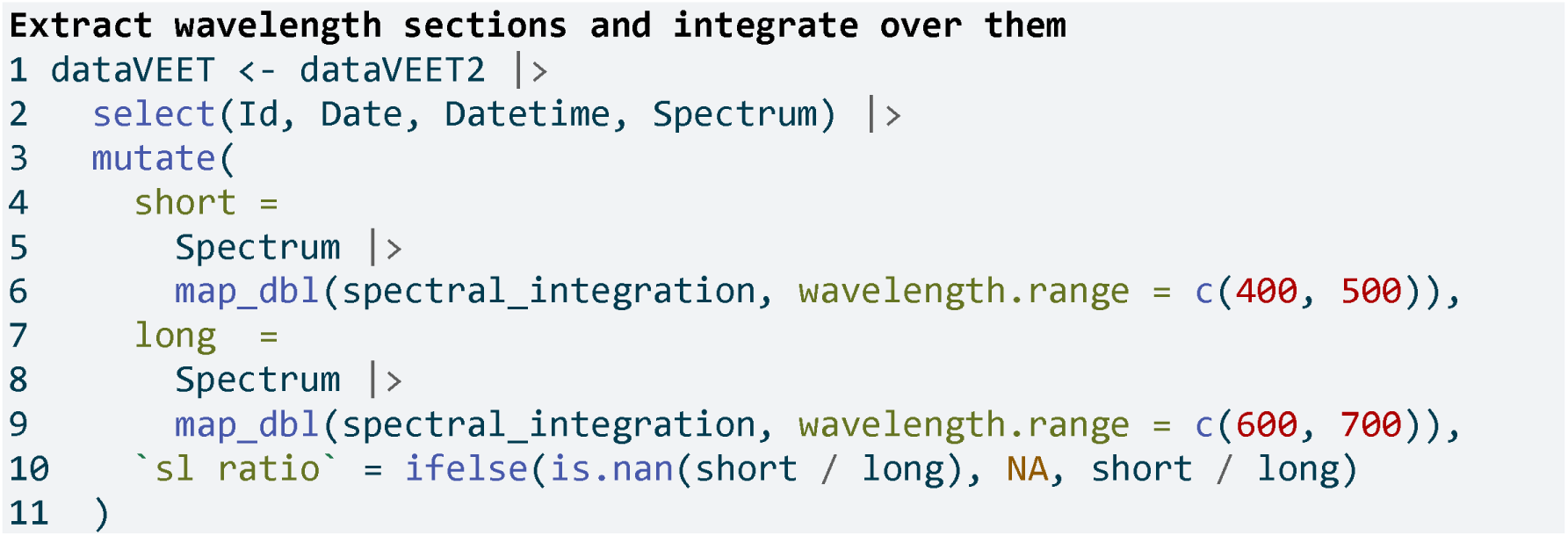

**Line 2** Focus on ID, date, time, and spectrum

**Line 10** Compute short-to-long wavelength ratio

**Table 17** shows the average short/long wavelength ratio, averaged over each day (and then as weekday/weekend means if applicable). In this dataset, the values give an indication of the spectral balance of the light the individual was exposed to (higher values mean relatively more short-wavelength content).

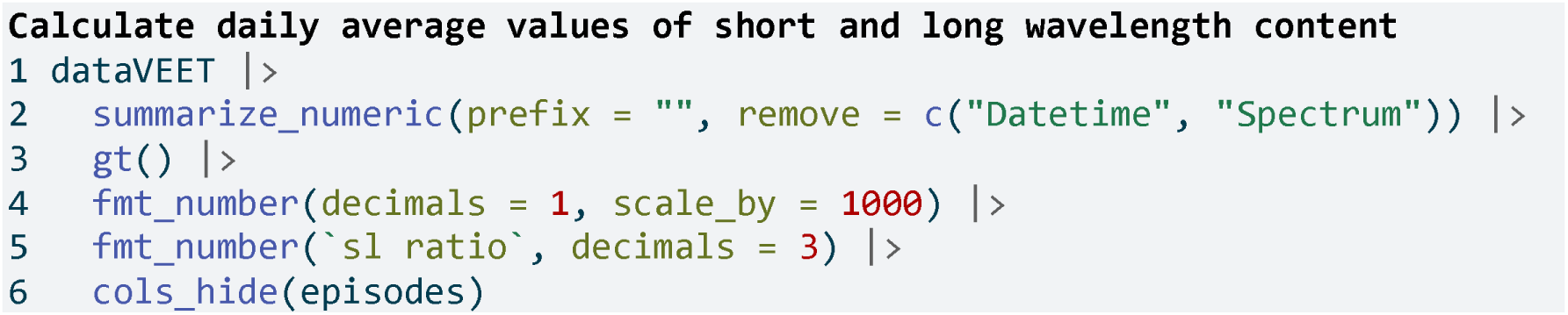

**Table 16:**
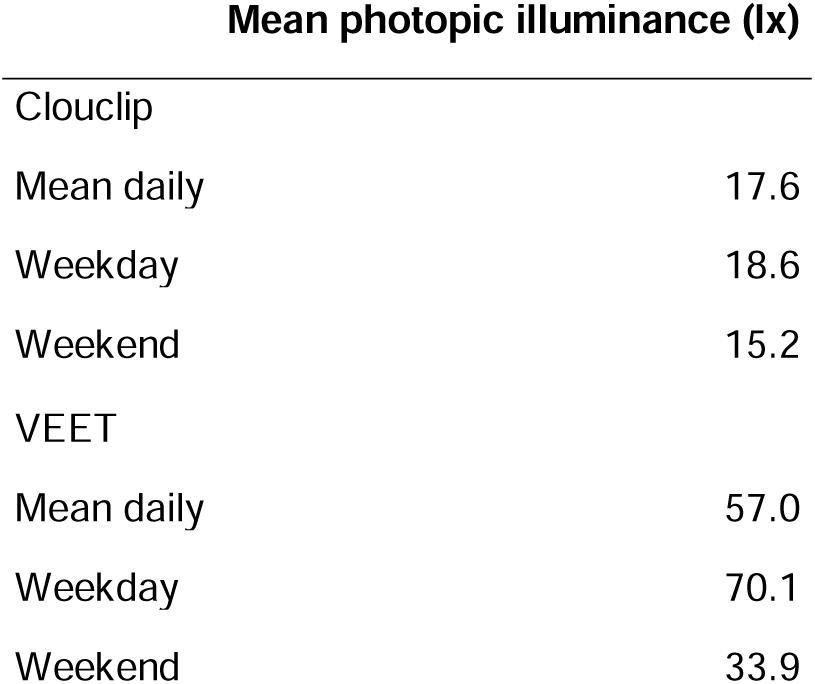
Recalculation of the mean light exposure per day (after logarithmic transformation to account for zero inflation and skewness) with the merged dataset

**Table 17:**
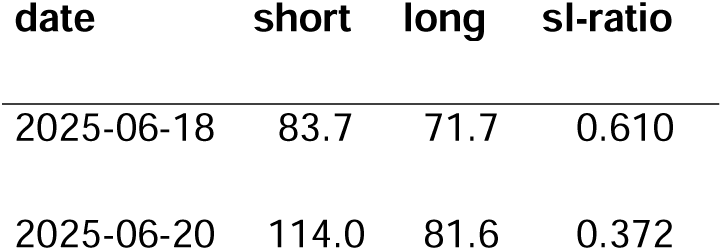
Average (mW/m²) and ratio of short-wavelength (400–500 nm) to long-wavelength (600–700 nm) light

### Melanopic daylight efficacy ratio (MDER)

The same idea is behind calculating the melanopic daylight efficacy ratio (or MDER), which is defined by the CIE (“CIE System for Metrology of Optical Radiation for ipRGC-Influenced Responses to Light” 2018) as the melanopic EDI divided by the photopic illuminance (Hartmeyer and Andersen 2023). Results are shown in **Table 18**. In this case, instead of a simple integration over a wavelength band, we apply an action spectrum to the spectral power distribution (SPD), integrate over the weighted SPD, and apply a correction factor. All alphaopic action spectra are implemented in the spectral_integration() function. These will result in photopic illuminance and melanopic equivalent daylight illuminance (melEDI)

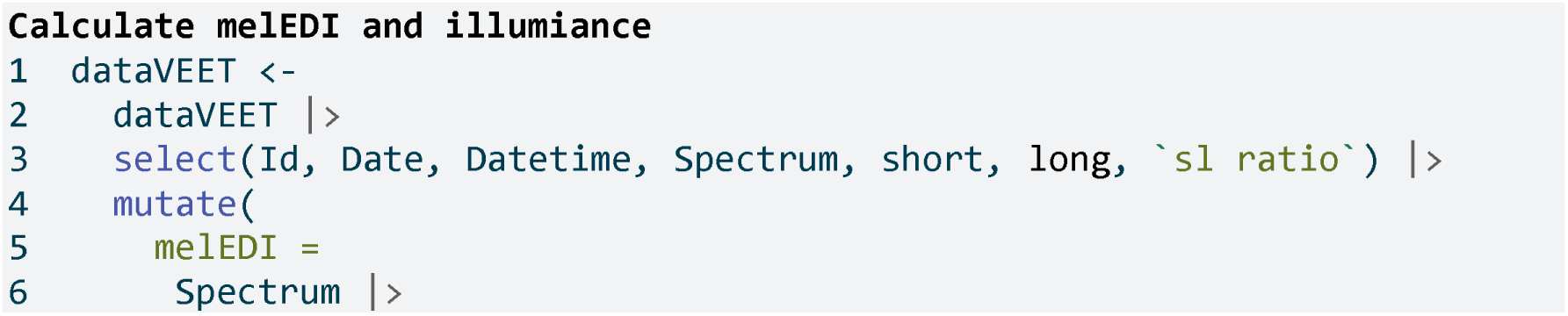
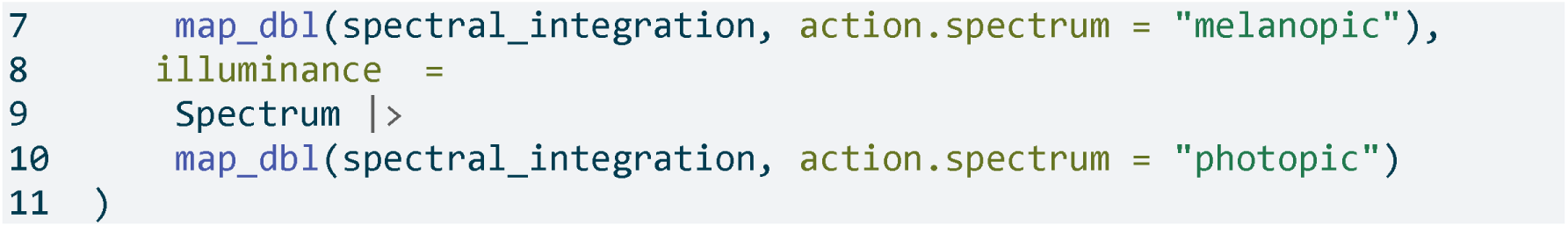

**Table 18:**
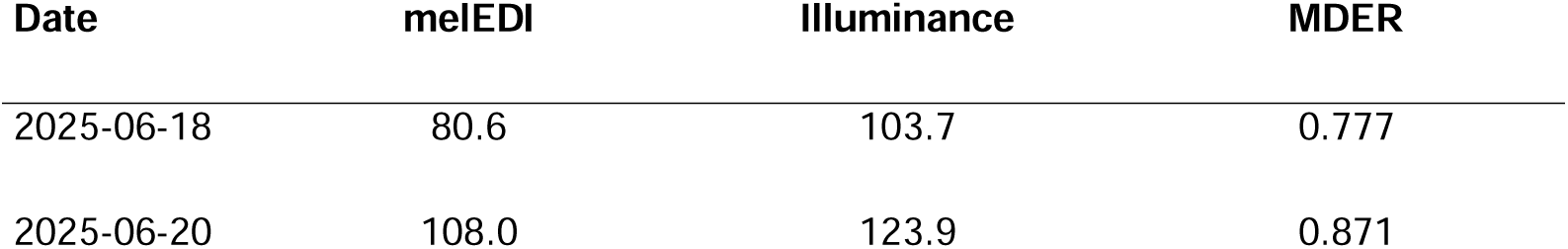
Average melanopic daylight efficacy ratio (MDER)

**Lines 5-7** Calculate melanopic EDI by applying the S_mel(l)_ action spectrum, integrating, and weighing

**Lines 8-10** Calculate photopic illuminance by applying the V_(l)_ action spectrum, integrating, and weighing

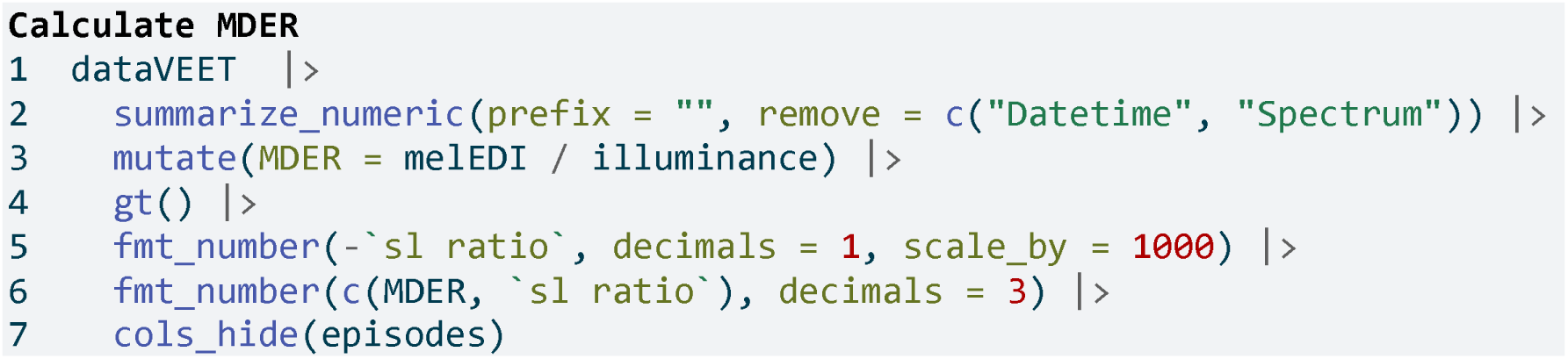

### Short-wavelength light at specific times of day

The third spectral example examines short-wavelength light exposure as a function of time of day. Certain studies might be interested in, for instance, blue-light exposure during midday versus morning or night. We demonstrate three approaches: (a) filtering the data to a specific local time window, and (b) aggregating by hour of day to see a daily profile of short-wavelength exposure. Additionally, we (c) look at differences between day and night periods.

### Local morning exposure

**Table 19** isolates the time window between 7:00 and 11:00 each day and computes the average short-wavelength irradiance in that interval. This represents a straightforward query: “How much blue light does the subject get in the morning on average?”

**Table 19:** Average short-wavelength light (400–500 nm) exposure between 7:00 and 11:00 each day

| Date | Short-wavelength<br>irradiance (mW/m²) |
| --- | --- |
| 2025-06-18 | 5.44 |
| 2025-06-20 | 0.95 |

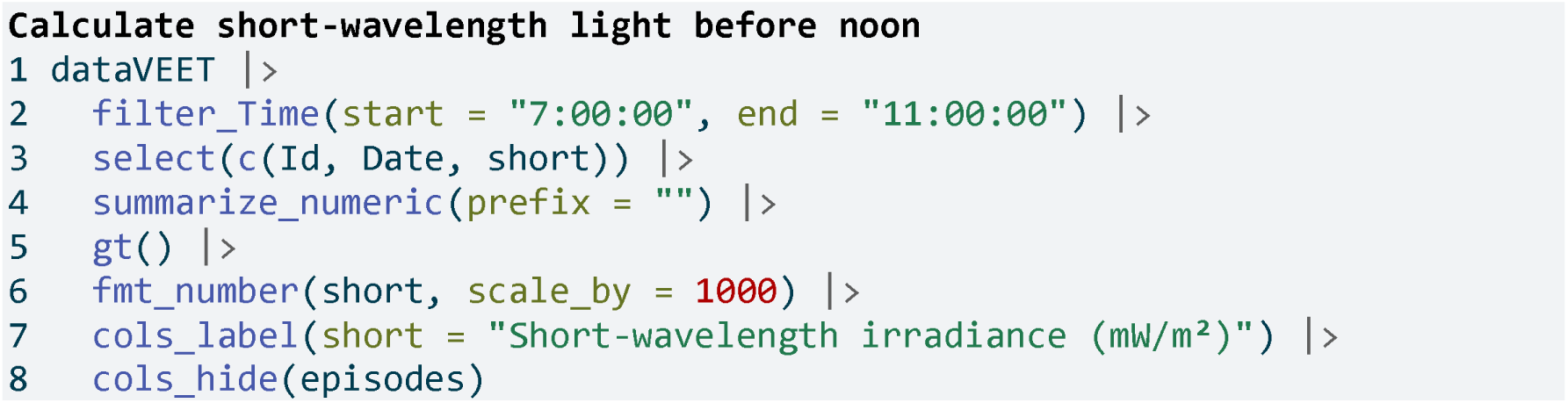

**Line 2** Filter data to local 7am–11am

### Hourly profile across the day

To visualize short-wavelength exposure over the course of a day, we aggregate the data into hourly bins. We cut the timeline into 1-hour segments (using local time), compute the mean short-wavelength irradiance in each hour for each day. Figure 11 shows the resulting diurnal profile, with short-wavelength exposure expressed as a fraction of the daily maximum for easier comparison.

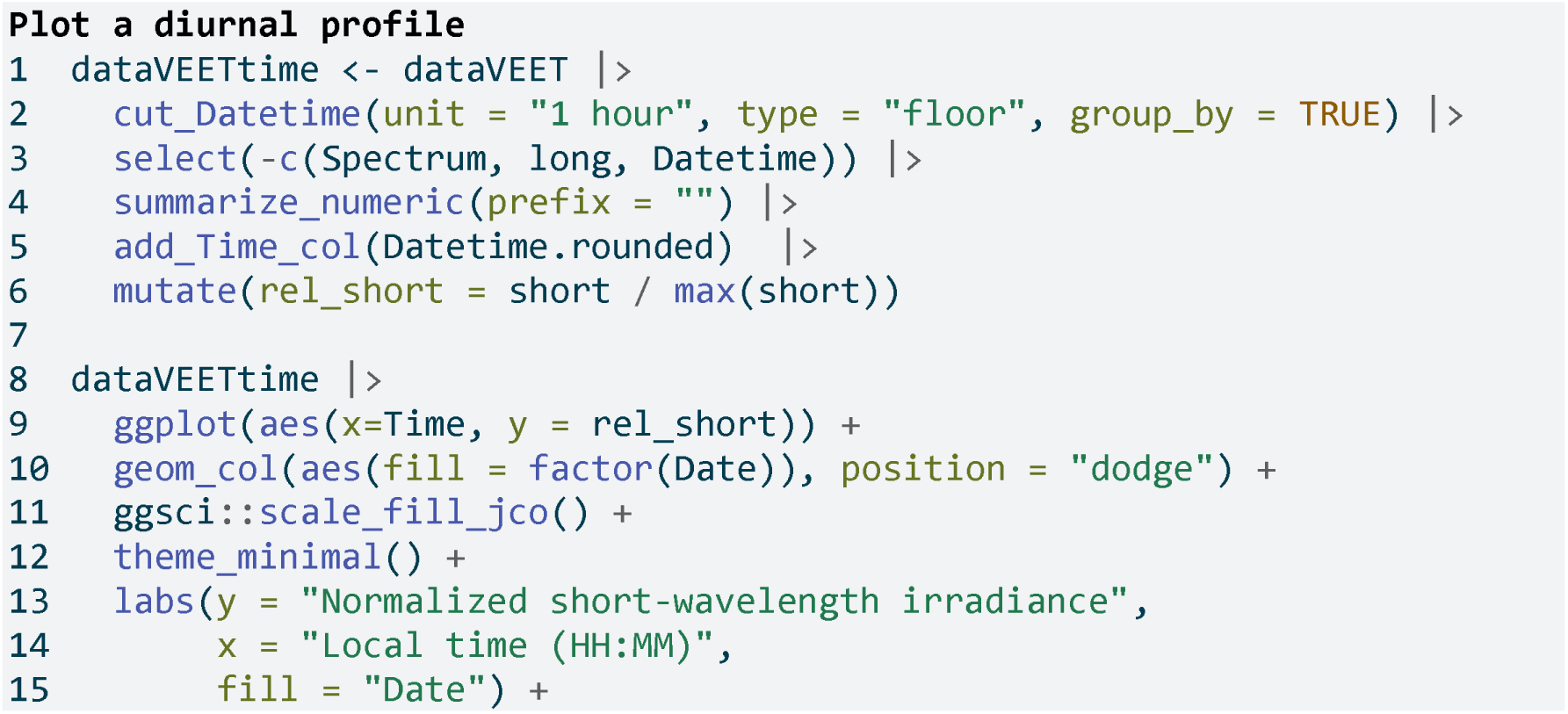
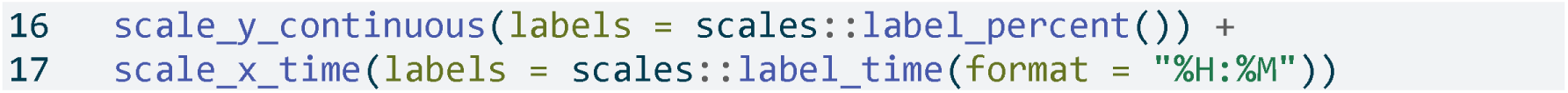

**Figure 11:**
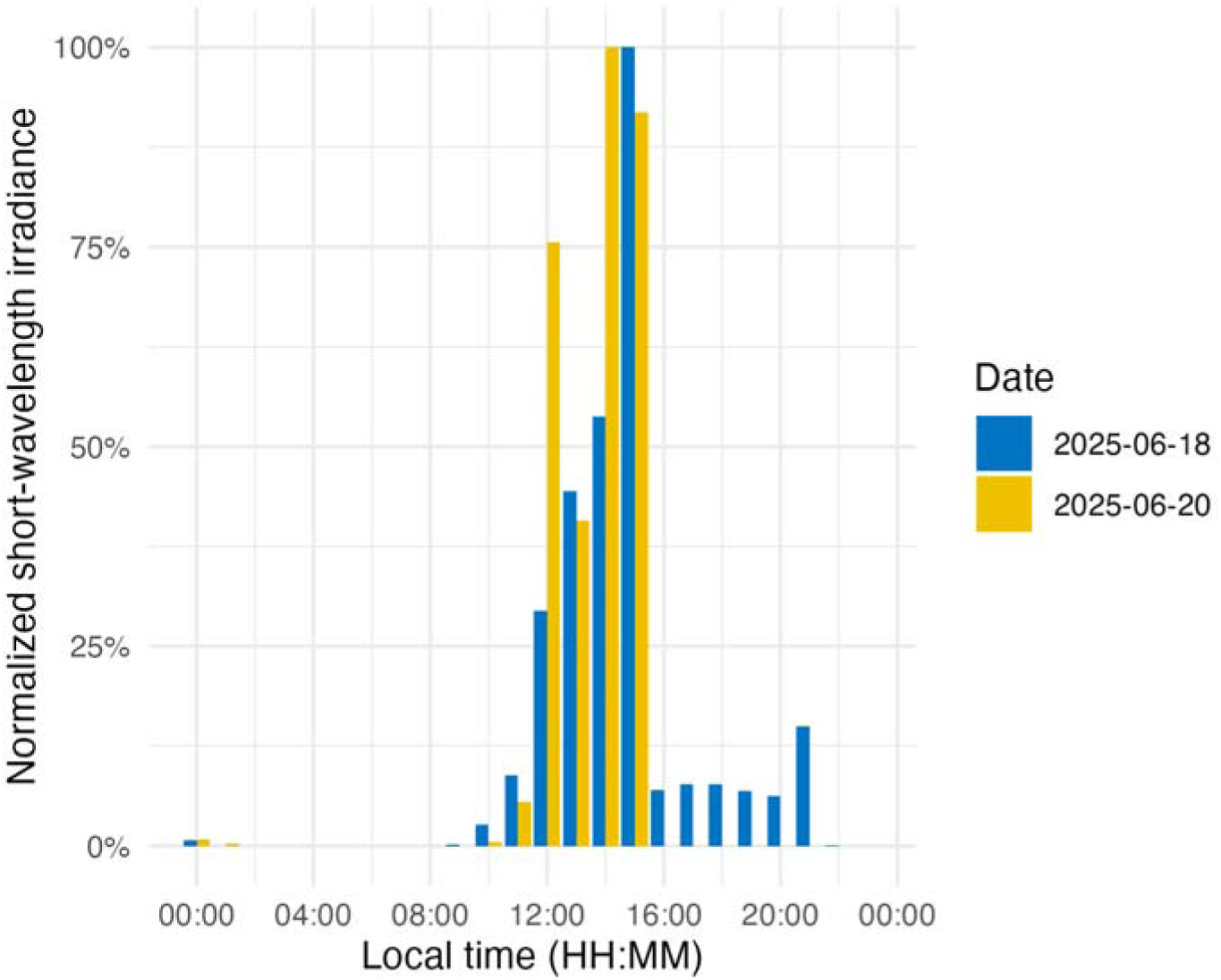
Diurnal profile of short-wavelength light exposure. Each bar represents the average short-wavelength irradiance at that hour of the day (0–23 h), normalized to the daily maximum.

**Line 1** Prepare hourly binned data

**Line 2** Bin timestamps by hour

**Line 5** Add a Time column (hour of day)

**Line 8-17** Creating the plot

### Day vs. night (photoperiod)

Finally, we compare short-wavelength exposure during daytime vs. nighttime. Using civil dawn and dusk information (based on geographic coordinates, here set for Houston, TX, USA), we label each measurement as day or night and then compute the total short-wavelength exposure in each period. **Table 20** summarizes the daily short-wavelength dose received during the day vs. during the night.

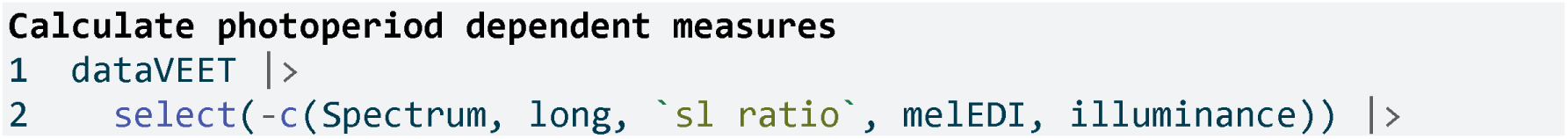
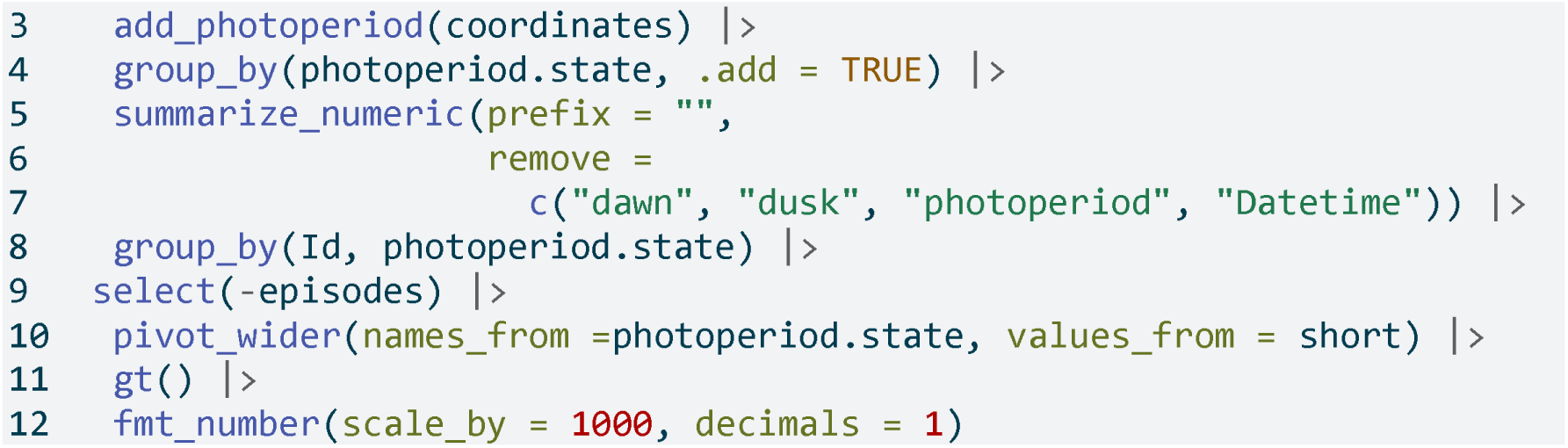

**Table 20:** Short wavelength light exposure (mW/m²) during the day and at night

| date | day | night |
| --- | --- | --- |
| 2025-06-18 | 126.5 | 12.3 |
| 2025-06-20 | 181.8 | 1.0 |

In the code cell above, add_photoperiod(coordinates) is used as a convenient way to add columns to the data frame, indicating for each timestamp whether it was day or night, given the latitude/longitude.

## Discussion and conclusion

This tutorial demonstrates a standardized, step-by-step pipeline to calculate a variety of visual experience metrics. We illustrated how a combination of LightLogR functions and tidyverse workflows can yield clear and reproducible analyses for wearable device data. While the full pipeline is detailed, each metric is computed through a dedicated sequence of well-documented steps, yet remains configurable to realize different metric definitions or thresholds.

By leveraging LightLogR’s framework alongside common data analysis approaches, the process remains transparent and relatively easy to follow. The overall goal is to make analysis transparent (with open-source functions), accessible (through thorough documentation, tutorials, and human-readable function naming, all under an MIT license), robust (the package includes >900 unit tests and continuous integration with bug tracking on GitHub), and community-driven (open feature requests and contributions via GitHub).

Even with standardized pipelines, researchers must still make and document many decisions during data cleaning, time-series handling, and metric calculations — especially for complex metrics that involve grouping data in multiple ways (for example, grouping by distance range as well as by duration for cluster metrics). We have highlighted these decision points in the tutorial (such as how to handle irregular intervals, choosing thresholds for “near” distances or “outdoor” light, and deciding on minimum durations for sustained events). Explicitly considering and reporting these choices is important for reproducibility and for comparing results across studies.

The broad set of features in LightLogR — ranging from data import and cleaning tools (for handling time gaps and irregularities) to visualization functions and metric calculators — make it a powerful toolkit for visual experience research. Our examples spanned circadian-light metrics and myopia-related metrics, demonstrating the versatility of a unified analysis approach. By using community-supported tools and workflows, researchers in vision science, chronobiology, myopia, and related fields can reduce time spent on low-level data wrangling and focus more on interpreting results and advancing scientific understanding.

## Statements

### Acknowledgement

We thank George Hatoun and David Sullivan (Reality Labs Research) for reviewing a draft of the tutorial manuscript and providing sample data for the VEET device for spectral analyses.

### Data availability statement

All data and code in this tutorial and Supplement 1 are available from the GitHub repository: https://github.com/tscnlab/ZaunerEtAl_JVis_2026/, archived on Zenodo: https://doi.org/10.5281/zenodo.16566014 under a MIT license (data under CC-BY license).

### Statement of generative AI and AI-assisted technologies in the writing process

The authors used ChatGPT during the preparation of this work. After using this tool, the authors reviewed and edited the content as needed and take full responsibility for the content of the publication.

Use of AI in contributor roles ^2^: Conceptualization: no Data curation: no Formal analysis: bug fixing Methodology: no Software: bug fixing Validation: no Visualization: tweaking of options Writing – original draft: abstract refinement Writing – review & editing: improve readability and language

## Footnotes

1 Note that older firmware versions of the VEET prior to 2.1.7 contained two Clear channels and the highest spectral channel was indicated as 940 nm. Data collected with this early firmware version are not suitable for spectral reconstruction in the context of research projects.

2 Based on the CRediT taxonomy. Funding acquisition, investigation, project administration, resources, and supervision were deemed irrelevant in this context and thus removed.

