## Supplement 1 for "Analysis of human visual experience data": Supplement.html


#### Table of contents

- Abstract
- 1 Introduction
- 2 `Clouclip` Data: Raw Structure and Import
- 3 `VEET` Data: Ambient Light (Illuminance) Processing
  - 3.1 Identifying non-wear times
- 4 `VEET` Data: Spectral Data Processing
  - 4.1 Import
  - 4.2 Spectral calibration
  - 4.3 Spectral reconstruction
- 5 `VEET` Data: Time of flight (distance)
  - 5.1 Removing non-wear times
- 6 Saving the Cleaned Data
- 7 References

### Supplement 1

 Code

- Show All Code
- Hide All Code
- ---
- View Source

Analysis of human visual experience data

Authors

Affiliations

Johannes Zauner

Technical University of Munich, Germany

Max Planck Institute for Biological Cybernetics, Tübingen, Germany

Aaron Nicholls

Reality Labs Research, USA

Lisa A. Ostrin

University of Houston College of Optometry, USA

Manuel Spitschan

Technical University of Munich, Germany

Max Planck Institute for Biological Cybernetics, Tübingen, Germany

Technical University of Munich Institute of advanced study (TUM-IAS), Munich, Germany

Doi

10.5281/zenodo.16566014

#### Abstract

This supplementary document provides a detailed, step-by-step tutorial on importing and preprocessing raw data from two wearable devices: `Clouclip` and `VEET`. We describe the structure of the raw datasets recorded by each device and explain how to parse these data, specify time zones, handle special sentinel values, clean missing observations, regularize timestamps to fixed intervals, and aggregate data as needed. All original R code from the main tutorial is shown here for transparency, with additional guidance intended for a broad research audience. We demonstrate how to detect gaps and irregular sampling, convert implicit missing periods into explicit missing values, and address device-specific quirks such as the `Clouclip`’s use of sentinel codes for “sleep mode” and “out of range” readings. Special procedures for processing the `VEET`’s rich spectral data (e.g. normalizing sensor counts and reconstructing full spectra from multiple sensor channels) are also outlined. Finally, we show how to save the cleaned datasets from both devices into a single R data file for downstream analysis. This comprehensive walkthrough is designed to ensure reproducibility and to assist researchers in understanding and adapting the data pipeline for their own visual experience datasets.

#### 1 Introduction

Wearable sensors like the `Clouclip` and the Visual Environment Evaluation Tool (`VEET`) produce high-dimensional time-series data on viewing distance and light exposure. Proper handling of these raw data is essential before any analysis of visual experience metrics. In the main tutorial, we introduce an analysis pipeline using the open-source R package LightLogR (Zauner, Hartmeyer, and Spitschan 2025) to calculate various distance and light exposure metrics. Here, we present a full account of the data import and preparation steps as a supplement to the methods, with an emphasis on clarity for researchers who may be less familiar with data processing in R. Figure 1 shows the main pre-processing steps and how they relate.

Code

```
flowchart TD

classDef input fill:#f7f7f7,stroke:#2f3d4a,color:#0f1a22,stroke-width:1px;
classDef process fill:#ffffff,stroke:#2f3d4a,color:#0f1a22,stroke-width:1px;
classDef output fill:#eef2f7,stroke:#2f3d4a,color:#0f1a22,stroke-width:1px;

%% ===== Raw inputs =====
RAW_CC[(Raw Clouclip file)]:::input
RAW_VEET[(Raw VEET file)]:::input

%% ===== Clouclip preprocessing =====
CC_IMP[Import<br/>Clouclip data]:::process
CC_QC[Inspect data,<br/>check for gaps and<br/>irregular data]:::process
CC_REG[Regularize timestamps<br/>and handle gaps]:::process
CC_VIS[Visualize cleaned<br/>Clouclip data]:::process

RAW_CC --> Clouclip
subgraph Clouclip[**Light and distance**]
CC_IMP --> CC_QC
CC_QC --> CC_REG
CC_REG --> CC_VIS
end

%% ===== VEET ALS light preprocessing =====
ALS_IMP[Import VEET<br/>*ALS*<br/>light modality]:::process
ALS_QC[Inspect, adjust interval,<br/>handle gaps, remove high<br/>missing days]:::process
IMU_IMP[Import VEET<br/>*IMU*<br/>actigraphy modality]:::process
IMU_NONWEAR[Detect **non-wear**<br/>using activity rules]:::process
ALS_NONWEAR[Add, label and visualize<br/>**non-wear**]:::process
ALS_CLEAN[Remove **non-wear**<br/>observations]:::process

RAW_VEET --> ALS_IMP
RAW_VEET --> IMU_IMP
subgraph Light[**Ambient light**]
ALS_IMP --> ALS_QC
ALS_QC --> ALS_NONWEAR
IMU_IMP --> IMU_NONWEAR
IMU_NONWEAR --> ALS_NONWEAR
ALS_NONWEAR --> ALS_CLEAN
end

%% ===== VEET spectral preprocessing =====
SPEC_IMP[Import VEET<br/>*PHO*<br/>spectral channels]:::process
SPEC_NORM[Normalize spectral<br/>channels by gain and<br/>integration time]:::process
SPEC_CLEAN[Inspect, adjust interval,<br/>handle gaps, remove high<br/>missing days]:::process
SPEC_CALfile[(Calibration matrix)]:::input
SPEC_CAL[Import calibration matrix]:::process
SPEC_RECON[Reconstruct and visualize<br/>spectra,<br/>calculate illuminance]:::process

RAW_VEET --> SPEC_IMP
subgraph Spectrum[**Spectral data**]
SPEC_IMP --> SPEC_NORM
SPEC_NORM --> SPEC_CLEAN
SPEC_CLEAN --> SPEC_RECON
SPEC_CALfile --> SPEC_CAL
SPEC_CAL --> SPEC_RECON
end

%% ===== VEET distance preprocessing =====
TOF_IMP[Import VEET distance<br/>*TOF*<br/>modality]:::process
TOF_QC[Inspect, adjust interval,<br/>handle gaps, remove high<br/>missing days]:::process
TOF_PIV[Pivot from *wide* to *long*<br/>format and label spatial<br/>position]:::process
TOF_NONWEAR[Add, label and remove<br/>**non-wear**]:::process

IMU_NONWEAR -.-> TOF_NONWEAR
RAW_VEET --> Distance
subgraph Distance[**Spatial distance**]
TOF_IMP --> TOF_QC
TOF_QC --> TOF_PIV
TOF_PIV --> TOF_NONWEAR
end
%% ===== Save outputs =====
SAVE[(Pre-processed datasets)]:::input

CC_VIS -- Cleaned Clouclip dataset **dataCC** --> SAVE
ALS_CLEAN -- Cleaned VEET ALS dataset **dataVEET** --> SAVE
SPEC_RECON -- Cleaned VEET spectral dataset **dataVEET2** --> SAVE
TOF_NONWEAR -- Cleaned VEET distance dataset **dataVEET3** --> SAVE

class Clouclip,Distance,Spectrum,Light output;
```

```
flowchart TD

classDef input fill:#f7f7f7,stroke:#2f3d4a,color:#0f1a22,stroke-width:1px;
classDef process fill:#ffffff,stroke:#2f3d4a,color:#0f1a22,stroke-width:1px;
classDef output fill:#eef2f7,stroke:#2f3d4a,color:#0f1a22,stroke-width:1px;

%% ===== Raw inputs =====
RAW_CC[(Raw Clouclip file)]:::input
RAW_VEET[(Raw VEET file)]:::input

%% ===== Clouclip preprocessing =====
CC_IMP[Import<br/>Clouclip data]:::process
CC_QC[Inspect data,<br/>check for gaps and<br/>irregular data]:::process
CC_REG[Regularize timestamps<br/>and handle gaps]:::process
CC_VIS[Visualize cleaned<br/>Clouclip data]:::process

RAW_CC --> Clouclip
subgraph Clouclip[**Light and distance**]
CC_IMP --> CC_QC
CC_QC --> CC_REG
CC_REG --> CC_VIS
end

%% ===== VEET ALS light preprocessing =====
ALS_IMP[Import VEET<br/>*ALS*<br/>light modality]:::process
ALS_QC[Inspect, adjust interval,<br/>handle gaps, remove high<br/>missing days]:::process
IMU_IMP[Import VEET<br/>*IMU*<br/>actigraphy modality]:::process
IMU_NONWEAR[Detect **non-wear**<br/>using activity rules]:::process
ALS_NONWEAR[Add, label and visualize<br/>**non-wear**]:::process
ALS_CLEAN[Remove **non-wear**<br/>observations]:::process

RAW_VEET --> ALS_IMP
RAW_VEET --> IMU_IMP
subgraph Light[**Ambient light**]
ALS_IMP --> ALS_QC
ALS_QC --> ALS_NONWEAR
IMU_IMP --> IMU_NONWEAR
IMU_NONWEAR --> ALS_NONWEAR
ALS_NONWEAR --> ALS_CLEAN
end

%% ===== VEET spectral preprocessing =====
SPEC_IMP[Import VEET<br/>*PHO*<br/>spectral channels]:::process
SPEC_NORM[Normalize spectral<br/>channels by gain and<br/>integration time]:::process
SPEC_CLEAN[Inspect, adjust interval,<br/>handle gaps, remove high<br/>missing days]:::process
SPEC_CALfile[(Calibration matrix)]:::input
SPEC_CAL[Import calibration matrix]:::process
SPEC_RECON[Reconstruct and visualize<br/>spectra,<br/>calculate illuminance]:::process

RAW_VEET --> SPEC_IMP
subgraph Spectrum[**Spectral data**]
SPEC_IMP --> SPEC_NORM
SPEC_NORM --> SPEC_CLEAN
SPEC_CLEAN --> SPEC_RECON
SPEC_CALfile --> SPEC_CAL
SPEC_CAL --> SPEC_RECON
end

%% ===== VEET distance preprocessing =====
TOF_IMP[Import VEET distance<br/>*TOF*<br/>modality]:::process
TOF_QC[Inspect, adjust interval,<br/>handle gaps, remove high<br/>missing days]:::process
TOF_PIV[Pivot from *wide* to *long*<br/>format and label spatial<br/>position]:::process
TOF_NONWEAR[Add, label and remove<br/>**non-wear**]:::process

IMU_NONWEAR -.-> TOF_NONWEAR
RAW_VEET --> Distance
subgraph Distance[**Spatial distance**]
TOF_IMP --> TOF_QC
TOF_QC --> TOF_PIV
TOF_PIV --> TOF_NONWEAR
end
%% ===== Save outputs =====
SAVE[(Pre-processed datasets)]:::input

CC_VIS -- Cleaned Clouclip dataset **dataCC** --> SAVE
ALS_CLEAN -- Cleaned VEET ALS dataset **dataVEET** --> SAVE
SPEC_RECON -- Cleaned VEET spectral dataset **dataVEET2** --> SAVE
TOF_NONWEAR -- Cleaned VEET distance dataset **dataVEET3** --> SAVE

class Clouclip,Distance,Spectrum,Light output;
```

Figure 1: Pre-processing steps in the supplement document

We use example datasets from a `Clouclip` device (Wen et al. 2021, 2020) and from a `VEET` device (Sah, Narra, and Ostrin 2025) (both provided in the accompanying repository). The `Clouclip` is a glasses-mounted sensor that records working distance (distance from eyes to object, in cm) and ambient illuminance (in lux) at 5-second intervals. The `VEET` is a head-mounted multi-modal sensor that logs ambient light and spectral information (along with other data like motion and distance via a depth sensor) - in this exemplary case at 2-second intervals. A single week of continuous data illustrates the contrast in complexity: approximately 1.6 MB for the `Clouclip`’s simple two-column output versus up to 270 MB for the `VEET`’s multi-channel output (due to its higher sampling rate and richer sensor modalities).

In the following sections, we detail how these raw data are structured and how to import and preprocess them using LightLogR. We cover device-specific considerations such as file format quirks and sensor range limitations, as well as general best practices like handling missing timestamps and normalizing sensor readings. All code blocks can be executed in R (with the required packages loaded) to reproduce the cleaning steps. The end result will be clean, regularized datasets (`dataCC` for Clouclip, `dataVEET` for VEET light data, `dataVEET2` for VEET spectral data, and `dataVEET3` for VEET distance) ready for calculating visual experience metrics. We conclude by saving these cleaned datasets into a single file for convenient reuse.

#### 2 `Clouclip` Data: Raw Structure and Import

The `Clouclip` device exports its data as a text file (not a true Excel file despite sometimes using an .xls extension), which is actually tab-separated values. See Figure 2 for the raw file.

Figure 2: Screenshot of the first rows of the `Clouclip` export file format as seen in a text editor

Each record corresponds to one timestamped observation (nominally every 5 seconds) and includes two measured variables: **distance** and **illuminance**. In the sample dataset (`Sample_Clouclip.csv` provided), the columns are:

- `Date` – the date and time of the observation (in the device’s local time, here one week in 2021).
- `Dis` – the viewing distance in centimeters.
- `Lux` – the ambient illuminance in lux.

For example, a raw data line might look like:

```
2021-07-01 08:00:00 45  320
```

indicating that at *2021-07-01 08:00:00* local time, the device recorded a working distance of 45 cm and illuminance of 320 lx. The Clouclip uses special **sentinel values**^1^ in these measurement columns to denote certain device states. Specifically, a distance (`Dis`) value of **204** is a code meaning the object is *out of the sensor’s range*, and a value of **-1** in either `Dis` or `Lux` indicates the device was in *sleep mode* (not actively recording). During normal operation, distance measurements are limited by the device’s range, and illuminance readings are positive lux values. Any sentinel codes in the raw file need to be handled appropriately, as described below.

We will use `LightLogR`’s built-in import function for `Clouclip`, which automatically reads the file, parses the timestamps, and converts sentinel codes into a separate status annotation. To begin, we load the necessary libraries and import the raw `Clouclip` dataset:

```
Load required packages
```

```
1library(tidyverse)
2library(LightLogR)
3library(gt)
4library(ggridges)
5library(downlit)
library(magick)
```

1
:   For tidy data science

2
:   Wearable analysis package

3
:   For great tables

4
:   For ridgeline plots

5
:   These packages are not used, but are needed for dependencies

```
Import of Clouclip data
```

```
1path <- "data/Sample_Clouclip.csv"
2tz   <- "US/Central"
3dataCC <- import$Clouclip(path, tz = tz, manual.id = "Clouclip")
```

1
:   Define file path

2
:   Time zone in which device was recording (e.g., US Central Time)

3
:   Import Clouclip data

```
Successfully read in 58'081 observations across 1 Ids from 1 Clouclip-file(s).
Timezone set is US/Central.
The system timezone is Europe/Berlin. Please correct if necessary!

First Observation: 2021-02-06 17:12:47
Last Observation: 2021-02-14 17:12:36
Timespan: 8 days

Observation intervals: 
  Id       interval.time            n pct     
1 Clouclip 5s                   54572 93.9601%
2 Clouclip 17s                     12 0.0207% 
3 Clouclip 18s                     14 0.0241% 
4 Clouclip 120s (~2 minutes)     3479 5.9900% 
5 Clouclip 128s (~2.13 minutes)     1 0.0017% 
6 Clouclip 132s (~2.2 minutes)      1 0.0017% 
7 Clouclip 133s (~2.22 minutes)     1 0.0017%
```

Figure 3: Overview plot of imported Clouclip data

In this code, `import$Clouclip()` reads the tab-delimited file and returns a **tibble**^2^ (saved in the variable `dataCC`) containing the data. We specify `tz = "US/Central"` because the device’s clock was set to U.S. Central time; this ensures that the `Datetime`values are properly interpreted with the correct time zone. The argument `manual.id = "Clouclip"` simply tags the dataset with an identifier (useful if combining data from multiple devices).

During import, LightLogR automatically handles the `Clouclip`’s sentinel codes. The `Date` column from the raw file is parsed into a `POSIXct` date-time (`Datetime`) with the specified time zone. The `Lux` and `Dis` columns are read as numeric, but any occurrences of **-1** or **204** are treated specially: these are replaced with `NA` (missing values) in the numeric columns, and corresponding status columns `Lux_status` and `Dis_status` are created to indicate the reason for those `NA` values. For example, if a `Dis` value of *204* was encountered, that row’s `Dis` will be set to NA and `Dis_status` will contain `"out_of_range"`; if `Lux` or `Dis` was -1, the status is `"sleep_mode"`. We will set all other readings to  `"operational"` (meaning the device was functioning normally at that time) for visualisation purposes.

```
Print the first 6 rows of the Clouclip dataset
```

```
dataCC |> head()
```

```
# A tibble: 6 × 7
# Groups:   Id [1]
  Id       file.name       Datetime              Dis   Lux Lux_status Dis_status
  <fct>    <chr>           <dttm>              <dbl> <dbl> <chr>      <chr>     
1 Clouclip Sample_Clouclip 2021-02-06 17:12:47    92    92 <NA>       <NA>      
2 Clouclip Sample_Clouclip 2021-02-06 17:12:52    92    92 <NA>       <NA>      
3 Clouclip Sample_Clouclip 2021-02-06 17:12:57    92    92 <NA>       <NA>      
4 Clouclip Sample_Clouclip 2021-02-06 17:13:02    92    92 <NA>       <NA>      
5 Clouclip Sample_Clouclip 2021-02-06 17:13:07    92    92 <NA>       <NA>      
6 Clouclip Sample_Clouclip 2021-02-06 17:13:12    92    92 <NA>       <NA>
```

After import, it is good practice to get an overview of the data. The import function by default prints a brief summary (and generates an overview plot of the timeline) showing the number of observations, the time span, and any irregularities or large gaps. In our case, the `Clouclip` summary indicates the data spans one week and reveals that there are **irregular intervals** in the timestamps. This means some observations do not occur exactly every 5 seconds as expected. We can programmatically check for irregular timing:

```
Check if data are on a perfectly regular 5-second schedule
```

```
dataCC |> has_irregulars()
```

```
[1] TRUE
```

If the result is `TRUE`, it confirms that the time sequence has deviations from the regular interval. Indeed, our example dataset has many small timing irregularities and gaps (periods with no data). Understanding the pattern of these missing or irregular readings is important. We can visualize them using a gap plot:

```
Plot gaps and irregular timestamps for Clouclip data
```

```
y.label <- "Distance (cm)"
1dataCC |> gg_gaps(Dis,
2                  include.implicit.gaps = FALSE,
3                  show.irregulars = TRUE,
                  y.axis.label = y.label,
                  group.by.days = TRUE, col.irregular = alpha("red", 0.03)
                  ) + labs(title = NULL)
```

1
:   Basing the gap-figure on the `Dis` distance variable

2
:   Only show missing observations (`NA` values)

3
:   Highlight irregular timing

Figure 4: Visualization of gaps and irregular data. Black traces show available data. Red shaded areas show times of missing data. Red dots show instances where observations occur off the regular interval from start to finish, i.e., irregular data.

In Figure 4, time periods where data are missing appear as red-shaded areas, and any off-schedule observation times are marked with red dots. The `Clouclip` example shows extensive gaps (red blocks) on certain days and irregular timing on all days except the first and last. These irregular timestamps likely arise from the device’s logging process (e.g. slight clock drift or buffering when the device was turned on/off). Such issues must be addressed before further analysis.

When faced with irregular or gapped data, we recommend a few strategies:

- *Remove leading/trailing segments that cause irregularity.* For example, if only the first day is regular and subsequent days drift, one might exclude the problematic portion using date filters (see filter\_Date() / filter\_Datetime() in LightLogR).
- *Round timestamps to the nearest regular interval.* This relabels slightly off-schedule times back to the 5-second grid (using cut\_Datetime() with a 5-second interval), provided the deviations are small and this rounding won’t create duplicate timestamps.
- *Aggregate to a coarser time interval.* For instance, grouping and averaging data into 1-minute bins with aggregate\_Datetime() can mask irregularities at finer scales, at the cost of some temporal resolution.

In this case, the deviations from the 5-second schedule are relatively minor. We choose to **round the timestamps to the nearest 5 seconds** to enforce a uniform sampling grid, which simplifies downstream gap handling. We further add a separate date column for convenience:

```
Regularize timestamps by rounding to nearest 5-second interval
```

```
dataCC <- dataCC |>
1  cut_Datetime("5 secs", New.colname = Datetime) |>
2  add_Date_col(group.by = TRUE)
```

1
:   Round times to 5-second bins

2
:   Add a Date column for grouping by day

After this operation, all `Datetime` entries in `dataCC` align perfectly on 5-second boundaries (e.g. **08:00:00**, **08:00:05**, **08:00:10**, etc.). We can verify that no irregular intervals remain by re-running `has_irregulars()`:

```
Re-check if data are on a perfectly regular 5-second schedule
```

```
dataCC |> has_irregulars()
```

```
[1] FALSE
```

Next, we want to quantify the missing data. LightLogR distinguishes between **explicit missing values** (actual `NA`s in the data, possibly from sentinel replacements or gaps we have filled in) and **implicit missing intervals** (time points where the device *should* have a reading but none was recorded, and we have not yet filled them in). See Figure 5 for a visual aid to these terms. Initially, many gaps are still implicit (between the first and last timestamp of each day).

Terminology of gaps and irregular data in LightLogR

Handling of gaps with LightLogR’s `gap_handler()` function

Figure 5: Gaps and irregular data

We can generate a gap summary table:

```
Summarize observed vs missing data by day for distance
```

```
dataCC |> gap_table(Dis, Variable.label = "Distance (cm)") |>
1  cols_hide(contains("_n"))
```

1
:   Hide absolute counts for brevity in output

Table 1: Summary of missing and observed data for the Clouclip device

|  |  |  |  |  |  |  |  |  |  |  |  |  |  |
| --- | --- | --- | --- | --- | --- | --- | --- | --- | --- | --- | --- | --- | --- |
| Summary of available and missing data | | | | | | | | | | | | | |
| Variable: Distance (cm) | | | | | | | | | | | | | |
|  | Data | | | | | Missing | | | | | | | |
|  | Regular | | Irregular | Range | Interval | Gaps | | | | Implicit | | Explicit | |
|  | Time | % | n^1,2^ | Time | Time | N | ø | Time | % | Time | % | Time | % |
| **Overall** | 2d 14h 43m 10s | 29.0%^3^ | 0 | 1w 2d | 5 | 2,690 | 1h 35m 57s | 6d 9h 16m 50s | 71.0%^3^ | 5d 15h 19m 55s | 62.7%^3^ | 17h 56m 55s | 8.3%^3^ |
| Clouclip - 2021-02-06 | | | | | | | | | | | | | |
|  | 43m 30s | 3.0% | 0 | 1d | 5s | 26 | 53m 43s | 23h 16m 30s | 97.0% | 22h 53m 20s | 95.4% | 23m 10s | 1.6% |
| Clouclip - 2021-02-07 | | | | | | | | | | | | | |
|  | 2h 45m | 11.5% | 0 | 1d | 5s | 139 | 9m 10s | 21h 15m | 88.5% | 19h 42m 35s | 82.1% | 1h 32m 25s | 6.4% |
| Clouclip - 2021-02-08 | | | | | | | | | | | | | |
|  | 11h 13m 55s | 46.8% | 0 | 1d | 5s | 443 | 1m 44s | 12h 46m 5s | 53.2% | 10h 47m 50s | 45.0% | 1h 58m 15s | 8.2% |
| Clouclip - 2021-02-09 | | | | | | | | | | | | | |
|  | 8h 46m 25s | 36.6% | 0 | 1d | 5s | 278 | 3m 17s | 15h 13m 35s | 63.4% | 13h 50m | 57.6% | 1h 23m 35s | 5.8% |
| Clouclip - 2021-02-10 | | | | | | | | | | | | | |
|  | 7h 1m 30s | 29.3% | 0 | 1d | 5s | 367 | 2m 47s | 16h 58m 30s | 70.7% | 14h 18m 40s | 59.6% | 2h 39m 50s | 11.1% |
| Clouclip - 2021-02-11 | | | | | | | | | | | | | |
|  | 8h 31m 55s | 35.5% | 0 | 1d | 5s | 423 | 2m 12s | 15h 28m 5s | 64.5% | 11h 43m 30s | 48.9% | 3h 44m 35s | 15.6% |
| Clouclip - 2021-02-12 | | | | | | | | | | | | | |
|  | 12h 17m 55s | 51.2% | 0 | 1d | 5s | 417 | 1m 41s | 11h 42m 5s | 48.8% | 9h 17m 45s | 38.7% | 2h 24m 20s | 10.0% |
| Clouclip - 2021-02-13 | | | | | | | | | | | | | |
|  | 10h 32m 15s | 43.9% | 0 | 1d | 5s | 527 | 1m 32s | 13h 27m 45s | 56.1% | 10h 42m 10s | 44.6% | 2h 45m 35s | 11.5% |
| Clouclip - 2021-02-14 | | | | | | | | | | | | | |
|  | 50m 45s | 3.5% | 0 | 1d | 5s | 70 | 19m 51s | 23h 9m 15s | 96.5% | 22h 4m 5s | 92.0% | 1h 5m 10s | 4.5% |
|  |  |  |  |  |  |  |  |  |  |  |  |  |  |
| --- | --- | --- | --- | --- | --- | --- | --- | --- | --- | --- | --- | --- | --- |
| ^1^ If n > 0: it is possible that the other summary statistics are affected, as they are calculated based on the most prominent interval. | | | | | | | | | | | | | |
| ^2^ Number of (missing or actual) observations | | | | | | | | | | | | | |
| ^3^ Based on times, not necessarily number of observations | | | | | | | | | | | | | |

This summary (Table 1) breaks down, for each day, how much data is present vs. missing. It reports the total duration of recorded data and the duration of gaps. After rounding the times, there are no irregular timestamp issues, but we see substantial **implicit gaps** — periods where the device was not recording (e.g., overnight when the device was likely not worn or was in sleep mode). Notably, the first and last days of the week have very little data (less than 1 hour each), since they probably represent partial recording days (the trial started and ended on those days).

To prepare the dataset for analysis, we will convert all those implicit gaps into explicit missing entries, and remove days that are mostly incomplete. Converting implicit gaps means inserting rows with `NA` for each missing 5-second slot, so that the time series becomes continuous and explicit about missingness (see Figure 5). We use `gap_handler()` for this, and then drop the nearly-empty days:

```
Convert implicit gaps to explicit NA gaps, and drop days with <1 hour of data
```

```
dataCC <- dataCC |> 
1  mutate(across(c(Lux_status, Dis_status), ~ replace_na(.x, "operational"))) |>
2  gap_handler(full.days = TRUE) |>
3  remove_partial_data(Dis, threshold.missing = "-1 hour")
```

1
:   First ensure that status columns have an “operational” tag for non-missing periods

2
:   Fill in all implicit gaps with explicit NA rows (for full days range)

3
:   Remove any day that has less than 1 hour of recorded data

After these steps, `dataCC` contains continuous 5-second timestamps for each day that remains. We chose a threshold of “-1 hours” to remove days with less than one hour of data, which in this dataset removes the first and last (partial) days. The cleaned `Clouclip` data now covers six full days with bouts of continuous wear.

It is often helpful to double-check how the sentinel values and missing data are distributed over time. We can visualize the distance time-series with status annotations and day/night periods:

```
Visualize observations and sentinel states
```

```
1coordinates <- c(29.75, -95.36)
dataCC |> 
2  fill(c(Lux_status, Dis_status), .direction = "downup") |>
3  gg_day(y.axis = Dis, geom = "line", y.axis.label = y.label) |>
4  gg_state(Dis_status, aes_fill = Dis_status, ymin = 0, ymax = 0.5, alpha = 1) |>
5  gg_photoperiod(coordinates, alpha = 0.1) +
  theme(legend.position = "bottom")
```

1
:   Setting coordinates for Houston, Texas (recording location)

2
:   Retain status markers until a new marker arrives

3
:   Create the basic plot

4
:   Add the status times

5
:   Add the photoperiod (day/night)

Figure 6: Distance measurements across days. Blue, grey and yellow-colored bars at the bottom of each day show sentinel states of the device. Blue indicates an operational status, grey sleep mode (not recording), and yellow an out of range measurement. Shaded areas in the main plot show nighttime from civil dusk until dawn, which are calculated based on the recording date and geographic coordinates

In this plot, blue segments indicate times when the Clouclip was *operational* (actively measuring), grey segments indicate the device in *sleep mode* (no recording, typically at night), and yellow segments indicate *out-of-range* distance readings. The red outlined regions show nighttime (from civil dusk to dawn) based on the given latitude/longitude and dates. As expected, most of the grey “sleep” periods align with night hours, and we see a few yellow spans when the user’s viewing distance exceeded the device’s range (e.g. presumably when no object was within 100 cm, such as when the user looked into the distance). At this stage, the Clouclip dataset `dataCC` is fully preprocessed: all timestamps are regular 5-second intervals, missing data are explicitly marked, extraneous partial days are removed, and sentinel codes are handled via the status columns. The data are ready for calculating daily distance and light exposure metrics (as done in the main tutorial’s Results).

#### 3 `VEET` Data: Ambient Light (Illuminance) Processing

The **VEET** device Sullivan et al. (2024) is a more complex logger that records multiple data modalities in one combined file. Its raw data file contains interleaved records for different sensor types, distinguished by a “modality” field. We focus first on the ambient light sensor modality (abbreviated **ALS**), which provides broad-spectrum illuminance readings (lux) and related information like sensor gains and flicker, recorded every 2 seconds. Later we will import the spectral sensor modality (**PHO**) for spectral irradiance data, and the time-of-flight modality (**TOF**) for distance data.

In the `VEET`’s export file, each line includes a timestamp and a modality code, followed by fields specific to that modality. Importantly, this means that the `VEET` export is not rectangular, i.e., tabular (see Figure 7). This makes it challenging for many import functions that expect the equal number of columns in every row, which is not the case in this instance. For the **ALS modality**, the relevant columns include a high-resolution timestamp (in Unix epoch format), integration time, UV/VIS/IR sensor gain settings, raw UV/VIS/IR sensor counts, a flicker measurement, and the computed illuminance in lux. For example, the ALS data columns are named: `time_stamp`, `integration_time`, `uvGain`, `visGain`, `irGain`, `uvValue`, `visValue`, `irValue`, `Flicker`, and `Lux`.

Figure 7: Screenshot of the first rows of the `VEET` export file format as seen in a text editor

For the **PHO (spectral) modality**, the columns include a timestamp, integration time, a general `Gain` factor, and nine sensor channel readings covering different wavelengths (with names like `s415, s445, ..., s680, s940`) as well as a `Dark` channel, a broadband channel `Clear`, and another broadband channel for flicker detection (`FD`). In essence, the `VEET`’s spectral sensor captures light in several wavelength bands (from ~415 nm up to 940 nm) rather than outputting a single lux value like the ambient light sensor does (*PHO*).

To import the `VEET` ambient light data, we again use the LightLogR import function, specifying the `ALS` modality. The raw VEET data in our example is provided as a zip file (`01_VEET_L.csv.zip`) containing the logged data for one week. We do the following:

```
Import VEET Ambient Light Sensor (ALS) data
```

```
path <- "data/01_VEET_L.csv.zip"
tz   <- "US/Central"
1dataVEET <- import$VEET(path, tz = tz, modality = "ALS", manual.id = "VEET")
```

1
:   In difference to the `Clouclip` file, we simply respecify the device type with `import$VEET(...)`, but must also provide a `modality` argument.

```
Successfully read in 304'193 observations across 1 Ids from 1 VEET-file(s).
Timezone set is US/Central.
The system timezone is Europe/Berlin. Please correct if necessary!
1 observations were dropped due to a missing or non-parseable Datetime value (e.g., non-valid timestamps during DST jumps). 

First Observation: 2024-06-04 15:00:37
Last Observation: 2024-06-12 08:29:43
Timespan: 7.7 days

Observation intervals: 
  Id    interval.time              n pct      
1 VEET  0s                         1 0.00033% 
2 VEET  1s                      1957 0.64334% 
3 VEET  2s                    300147 98.67025%
4 VEET  3s                      2074 0.68181% 
5 VEET  4s                         3 0.00099% 
6 VEET  9s                         5 0.00164% 
7 VEET  10s                        3 0.00099% 
8 VEET  109s (~1.82 minutes)       1 0.00033% 
9 VEET  59077s (~16.41 hours)      1 0.00033%
```

Figure 8: Overview plot of imported VEET data

Note

We get one warning as a single time stamp could not be parsed into a datetime - with ~300k observations, this is not an issue.

This call reads in only the lines corresponding to the `ALS` modality from the `VEET` file. The result `dataVEET` is a tibble^3^ with columns such as `Datetime` (parsed from the `time_stamp`  to POSIXct in US/Central time), `Lux` (illuminance in lux), `Flicker`, and the various sensor gains/values. Columns we don’t need for our analysis, like the modality code or file name, are also included but can be ignored or removed. From the import summary, we learn that the `VEET` light data, like the `Clouclip`, also exhibits irregularities and gaps. (The device nominally records every 2 seconds, but timing may drift or pause when not worn.)

```
Print the first 6 rows of the VEETs light dataset
```

```
dataVEET |> head()
```

```
# A tibble: 6 × 14
# Groups:   Id [1]
  Id    Datetime            file.name     time_stamp modality integration_time
  <fct> <dttm>              <chr>              <dbl> <chr>               <dbl>
1 VEET  2024-06-04 15:00:37 01_VEET_L.csv 1717531237 ALS                   100
2 VEET  2024-06-04 15:00:39 01_VEET_L.csv 1717531239 ALS                   100
3 VEET  2024-06-04 15:00:41 01_VEET_L.csv 1717531241 ALS                   100
4 VEET  2024-06-04 15:00:43 01_VEET_L.csv 1717531243 ALS                   100
5 VEET  2024-06-04 15:00:45 01_VEET_L.csv 1717531245 ALS                   100
6 VEET  2024-06-04 15:00:47 01_VEET_L.csv 1717531247 ALS                   100
# ℹ 8 more variables: uvGain <dbl>, visGain <dbl>, irGain <dbl>, uvValue <dbl>,
#   visValue <dbl>, irValue <dbl>, Flicker <dbl>, Lux <dbl>
```

To make the VEET light data comparable to the Clouclip’s and to simplify analysis, we choose to **aggregate the VEET illuminance data to 5-second intervals**. This slight downsampling will both reduce data volume and remove irregularities.

```
Aggregate VEET light data to 5-second intervals and mark gaps
```

```
dataVEET <- dataVEET |>
1  aggregate_Datetime(unit = "5 seconds") |>
2  gap_handler(full.days = TRUE) |>
3  add_Date_col(group.by = TRUE) |>
4  remove_partial_data(Lux, threshold.missing = "1 hour")
```

1
:   Resample to 5-sec bins (e.g. average Lux over 2-sec readings)

2
:   Fill in implicit gaps with NA rows

3
:   Add Date column for daily grouping

4
:   Remove participant days with more than one hour of missing data

First, `aggregate_Datetime(unit = "5 seconds")` combines the high-frequency 2-second observations into 5-second slots. By default, this function will average numeric columns like `Lux` over each 5-second period (and have sensible defaults for strings or categorical data). All of these *data type handlers* can be changed with the function call. The result is that `dataVEET` now has a reading every 5 seconds (or an NA if none were present in that window). Next, `gap_handler(full.days = TRUE)` inserts explicit NA entries for any 5-second timestamp that had no data within the continuous span of the recording. Then we add a `Date` column for grouping, and finally we remove days with more than 1 hour of missing data (using a more strict criterion as we did for `Clouclip`). According to the gap summary (Table 2), this leaves six full days of `VEET` light data with good coverage, after dropping the very incomplete start/end days.

We can inspect the missing-data summary for the `VEET` illuminance data:

```
dataVEET |> gap_table(Lux, "Illuminance (lx)") |> cols_hide(contains("_n"))
```

Table 2: Summary of missing and observed data for the VEET device, light modality

|  |  |  |  |  |  |  |  |  |  |  |  |  |  |
| --- | --- | --- | --- | --- | --- | --- | --- | --- | --- | --- | --- | --- | --- |
| Summary of available and missing data | | | | | | | | | | | | | |
| Variable: Illuminance (lx) | | | | | | | | | | | | | |
|  | Data | | | | | Missing | | | | | | | |
|  | Regular | | Irregular | Range | Interval | Gaps | | | | Implicit | | Explicit | |
|  | Time | % | n^1,2^ | Time | Time | N | ø | Time | % | Time | % | Time | % |
| **Overall** | 5d 23h 57m 40s | 100.0%^3^ | 0 | 6d | 5 | 8 | 58s | 2m 20s | 0.0%^3^ | 0s | 0.0%^3^ | 2m 20s | 0.0%^3^ |
| VEET - 2024-06-06 | | | | | | | | | | | | | |
|  | 23h 58m 5s | 99.9% | 0 | 1d | 5s | 3 | 38s | 1m 55s | 0.1% | 0s | 0.0% | 1m 55s | 0.1% |
| VEET - 2024-06-07 | | | | | | | | | | | | | |
|  | 1d | 100.0% | 0 | 1d | 5s | 0 | 0s | 0s | 0.0% | 0s | 0.0% | 0s | 0.0% |
| VEET - 2024-06-08 | | | | | | | | | | | | | |
|  | 23h 59m 55s | 100.0% | 0 | 1d | 5s | 1 | 5s | 5s | 0.0% | 0s | 0.0% | 5s | 0.0% |
| VEET - 2024-06-09 | | | | | | | | | | | | | |
|  | 23h 59m 50s | 100.0% | 0 | 1d | 5s | 2 | 5s | 10s | 0.0% | 0s | 0.0% | 10s | 0.0% |
| VEET - 2024-06-10 | | | | | | | | | | | | | |
|  | 23h 59m 55s | 100.0% | 0 | 1d | 5s | 1 | 5s | 5s | 0.0% | 0s | 0.0% | 5s | 0.0% |
| VEET - 2024-06-11 | | | | | | | | | | | | | |
|  | 23h 59m 55s | 100.0% | 0 | 1d | 5s | 1 | 5s | 5s | 0.0% | 0s | 0.0% | 5s | 0.0% |
|  |  |  |  |  |  |  |  |  |  |  |  |  |  |
| --- | --- | --- | --- | --- | --- | --- | --- | --- | --- | --- | --- | --- | --- |
| ^1^ If n > 0: it is possible that the other summary statistics are affected, as they are calculated based on the most prominent interval. | | | | | | | | | | | | | |
| ^2^ Number of (missing or actual) observations | | | | | | | | | | | | | |
| ^3^ Based on times, not necessarily number of observations | | | | | | | | | | | | | |

Table 2 shows, for each retained day, the total recorded duration and the duration of gaps. The `VEET` device, like the `Clouclip`, was not worn continuously 24 hours per day, so there are nightly gaps of wear time (when the device was likely off the participant), but not of recordings. After our preprocessing, any implicit gaps are represented as explicit missing intervals. The `VEET`’s time sampling was originally more frequent, but by aggregating to 5 s we have ensured a uniform timeline akin to the `Clouclip`’s.

At this point, the `dataVEET` object (for illuminance) is cleaned and ready for computing light exposure metrics. For example, one could calculate daily mean illuminance or the duration spent above certain light thresholds (e.g. “outdoor light exposure” defined as >1000 lx) using this dataset. Indeed, basic summary tables in the main tutorial illustrate the highly skewed nature of light exposure data and the calculation of outdoor light metrics. We will not repeat those metric calculations here in the supplement, as our focus is on data preprocessing; however, having a cleaned, gap-marked time series is crucial for those metrics to be accurate.

##### 3.1 Identifying non-wear times

Not all recorded time points of the `VEET` can be considered as `wear time`. `Non-wear` times should generally be labelled in the wearable time series, if possible. There are several ways how `Non-wear` can be classified (Zauner et al. 2025):

1. A device detects `non-wear` and does not record any measurements during non-wear times, or has a variable in the exported data to denote non-wear times. The `Clouclip` is an example here with the `sleep mode` sentinel state.
2. Record non-wear separately, e.g. with a wear-log or diary. This is easy to implement, but puts a higher burden on participants and research staff, and can be error prone: forgotten, or incorrect data entries can mis-classify wear and non-wear times. For an implementation on how to remove non-wear times based on such a log, see, e.g., this tutorial. These information can also be derived from contextual information. For example, the `VEET` records charging times in a separate file `log.csv`, which can be considered as definitive non-wear times.
3. Instruct participants with a certain behavior when removing the device to ease automated detection of non-wear times in the data. These include putting a device in a black opaque bag, so that sensor readings for light are zero, or pressing an event button (if one exists). Up- and downsides are similar to the previous option, but the measures are usually harder to decipher in the data: when a buttonpress was forgotten, does that mess up the automated detection moving forward, which expects two presses per non-wear period (at beginning and at start). When there is zero lux during the day - was it in a black bag, or was this person just in a dark environment?
4. Automated detection by an algorithm. There are several possibilities to try to classify non-wear times based on the available data from a wearable. E.g., if there is an activity tracker, that can be used to determine periods of inactivity, which could indicate non-wear. If only light data is available, the standard deviation (or coefficient of variance) of light in a moving window of a few minutes can be used to determine low variance, which also indicates inactivity. Usually a histogram of values shows a spike in those periods of inactivity.

NoteHow to handle non-wear intervals

Whether classified non-wear intervals should be removed, i.e., observations set to `NA`, or not should be a deliberate decision by the researcher. If it can be assumed that non-wear times misrepresent the measurement environment of the participant, then these observations are best set to `NA`. If, on the other hand, they might be somewhat representative, then it might be enough to label these times but still include them in analyses. Usually, this requires special instructions to the participants for non-wear. E.g., instructions for bed-time could include setting the device on the night stand facing into the room (i.e., sensors not obstructed). While this would not represent distance well, it can be considered as sufficient for ambient light measurements.

If datasets are large enough, i.e., collect data from a long enought period, `non-wear` has a diminishing effect, at least as daily metrics of light are concerned. E.g., in a monthlong study of 39 participants from Switzerland and Malaysia, (Biller et al. 2025) found that based on a sensitivity anaylsis up to 6 hours of data could be missing **every day**, and still the daily metrics across that month would not significantly change. This assumption only holds when non-wear times are not systematic, i.e., at the same time each day.

We will perform a simple variant of the third option, and look at the `VEET`’s `IMU` modality.

```
Import VEET activity (IMU) data
```

```
path <- "data/01_VEET_L.csv.zip"
tz   <- "US/Central"
dataIMU <- 
  import$VEET(path, tz = tz, modality = "IMU", manual.id = "VEET",
1              remove_duplicates = TRUE,
2              silent = TRUE)
```

1
:   Some rows in the data are duplicated. These will be removed during import

2
:   In this instance, we skip the import summary

```
15 duplicate rows were removed during import.
```

To get a quick feeling for the data, we visualize the activity variable for the x axis in Figure 9

```
Plotting the IMU timeline
```

```
dataIMU |> 
  gg_days(ax, 
          aes_col = abs(ax) < 1, 
          group = consecutive_id(abs(ax) < 1),
          y.axis.label = "ax"
          )
```

Figure 9: Timeline of the activity sensor (x direction)

`ax` seems to vary between ±10. In general, the periods of inactivity seem to lie within a band of ±1. However, simply choosing this band does not eliminate false detections during times of high movement. Some data transformations can make these more clear:

```
Plotting a transformed IMU timeline
```

```
dataIMU |> 
  aggregate_Datetime(
    "5 mins",
    numeric.handler = sd
  ) |> 
  pivot_longer(cols = c(ax, ay, az)) |> 
  group_by(name) |> 
  gg_days(value, 
          aes_col = value < 0.05,
          group = consecutive_id(value < 0.05),
          y.axis.label = "activity axes"
          )
```

Figure 10: Timeline of the activity sensor (x direction) - transformed to distinguish times of low activity

What Figure 10 shows is that the 5-minute standard deviation of the activity channels (x, y, and z direction) allows a pretty stable distinction between periods of high and low activity, with a threshold of **0.05**. We can us this to create a wear column. As the choice of channel does not seem to matter much, we will use `ax`.

```
Calculating times of wear and non-wear
```

```
wear_data <- 
dataIMU |> 
1  aggregate_Datetime("5 mins",numeric.handler = sd) |>
  mutate(wear = ax > 0.05) |>
  select(Id, Datetime, wear) |>
2  extract_states(wear)
3wear_data |>
  ungroup(Id) |>
  summarize_numeric(remove = c("start", "end", "epoch")) |>
  gt()
wear_data <- wear_data |> select(Id, wear, start, end)
```

1
:   Calculation of the wear-variable

2
:   Extracting the start and end times of wear and non-wear

3
:   Summarizing wear and non-wear times in a table

| wear | mean\_duration | total\_duration | episodes |
| --- | --- | --- | --- |
| FALSE | 11447s (~3.18 hours) | 366300s (~4.24 days) | 32 |
| TRUE | 9431s (~2.62 hours) | 301800s (~3.49 days) | 32 |

We see that most of the recorded timespan is actually non-wear (including sleep), with an average wear time of 2.6 hours.

Let’s add this data to the `ALS` dataset and visualize it in Figure 11

```
Plotting non-wear for light
```

```
dataVEET <- 
dataVEET |> 
  group_by(Id) |> 
  add_states(wear_data)

dataVEET |> 
  gg_days(Lux) |> 
  gg_states(wear, ymax = 0, alpha = 1, fill = "red")
```

Figure 11: Wear times for light data, shown as red bars

This looks sensible, e.g., when we look at noon of 7 June 2024. We can remove these observations based on the `wear` column to make our summaries more valid.

```
Removing non-wear observations from light modality
```

```
dataVEET <- dataVEET |> mutate(Lux = ifelse(wear, Lux, NA)) |> group_by(Id, Date)
```

#### 4 `VEET` Data: Spectral Data Processing

##### 4.1 Import

In addition to broad-band illuminance and distance, the `VEET` provides **spectral sensor data** through its `PHO` modality. Unlike illuminance, the spectral data are not given as directly interpretable radiometric metrics but rather as raw sensor counts across multiple wavelength channels, which require conversion to reconstruct a spectral power distribution. In our analysis, spectral data allow us to compute metrics like the relative contribution of short-wavelength (blue) light versus long-wavelength light in the participant’s environment. Processing this spectral data involves several necessary steps.

First, we import the spectral modality from a second `VEET` file. This time we need to extract the lines marked as `PHO`. We will store the spectral dataset in a separate object `dataVEET2` so as not to overwrite the `dataVEET` illuminance data in our R session:

```
Import VEET Spectral Sensor (PHO) data
```

```
path <- "data/02_VEET_L.csv.zip"
dataVEET2 <- import$VEET(path, tz = tz, modality = "PHO", manual.id = "VEET")
```

```
Successfully read in 173'013 observations across 1 Ids from 1 VEET-file(s).
Timezone set is US/Central.
The system timezone is Europe/Berlin. Please correct if necessary!

First Observation: 2025-06-17 12:25:13
Last Observation: 2025-06-21 22:47:01
Timespan: 4.4 days

Observation intervals: 
   Id    interval.time      n pct      
 1 VEET  1s               417 0.24102% 
 2 VEET  2s            171837 99.32086%
 3 VEET  3s               738 0.42656% 
 4 VEET  4s                 7 0.00405% 
 5 VEET  9s                 1 0.00058% 
 6 VEET  11s                1 0.00058% 
 7 VEET  12s                2 0.00116% 
 8 VEET  13s                4 0.00231% 
 9 VEET  15s                1 0.00058% 
10 VEET  16s                1 0.00058% 
# ℹ 3 more rows
```

Figure 12: Overview plot of imported VEET data

```
Print the first 6 rows of the VEETs spectral dataset
```

```
dataVEET2 |> head()
```

```
# A tibble: 6 × 18
# Groups:   Id [1]
  Id    Datetime            file.name time_stamp modality integration_time  Gain
  <fct> <dttm>              <chr>          <dbl> <chr>               <dbl> <dbl>
1 VEET  2025-06-17 12:25:13 02_VEET_… 1750181113 PHO                   100   512
2 VEET  2025-06-17 12:25:17 02_VEET_… 1750181117 PHO                   100   512
3 VEET  2025-06-17 12:25:19 02_VEET_… 1750181119 PHO                   100   512
4 VEET  2025-06-17 12:25:21 02_VEET_… 1750181121 PHO                   100   512
5 VEET  2025-06-17 12:25:23 02_VEET_… 1750181123 PHO                   100   512
6 VEET  2025-06-17 12:25:25 02_VEET_… 1750181125 PHO                   100   512
# ℹ 11 more variables: s415 <dbl>, s445 <dbl>, s480 <dbl>, s515 <dbl>,
#   s555 <dbl>, s590 <dbl>, s630 <dbl>, s680 <dbl>, s910 <dbl>, Dark <dbl>,
#   Clear <dbl>
```

After import, `data` contains columns for the timestamp (`Datetime`), `Gain` (the sensor gain setting), and the nine spectral sensor channels plus a clear channel. These appear as numeric columns named `s415, s445, ..., s940, Dark, Clear`. Other columns are also present but not needed for now. The spectral sensor was logging at a 2-second rate. It is informative to look at a snippet of the imported spectral data before further processing. Table 3 shows three rows of `data` after import (before calibration), with some technical columns omitted for brevity:

```
Table overview of spectral sensor data
```

```
dataVEET2 |> 
  slice(6000:6003) |> 
  select(-c(modality, file.name, time_stamp)) |> 
  gt() |> 
  fmt_number(s415:Clear)
```

Table 3: Overview of the spectral sensor import from the VEET device (3 observations). Each row corresponds to a 2-second timestamp (Datetime) and shows the raw sensor readings for the spectral channels (s415–s940, Dark, Clear). All values are in arbitrary sensor units (counts). Gain values and integration\_time are also relevant for each interval, depending on the downstream computation.

| Datetime | integration\_time | Gain | s415 | s445 | s480 | s515 | s555 | s590 | s630 | s680 | s910 | Dark | Clear |
| --- | --- | --- | --- | --- | --- | --- | --- | --- | --- | --- | --- | --- | --- |
| VEET | | | | | | | | | | | | | |
| --- | --- | --- | --- | --- | --- | --- | --- | --- | --- | --- | --- | --- | --- |
| 2025-06-18 00:11:22 | 100 | 512 | 20.00 | 27.00 | 31.00 | 42.00 | 47.00 | 63.00 | 90.00 | 139.00 | 756.00 | 0.00 | 534.00 |
| 2025-06-18 00:11:24 | 100 | 512 | 21.00 | 28.00 | 31.00 | 41.00 | 47.00 | 63.00 | 90.00 | 140.00 | 756.00 | 0.00 | 534.00 |
| 2025-06-18 00:11:26 | 100 | 512 | 21.00 | 27.00 | 31.00 | 41.00 | 47.00 | 63.00 | 90.00 | 139.00 | 756.00 | 0.00 | 534.00 |
| 2025-06-18 00:11:28 | 100 | 512 | 21.00 | 27.00 | 31.00 | 42.00 | 47.00 | 63.00 | 90.00 | 140.00 | 755.00 | 0.00 | 534.00 |

##### 4.2 Spectral calibration

Now we proceed with **spectral calibration**. The `VEET`’s spectral sensor counts need to be converted to physical units (spectral irradiance) via a calibration matrix provided by the manufacturer. For this example, we assume we have a calibration matrix that maps all the channel readings to an estimated spectral power distribution (SPD). The LightLogR package provides a function `spectral_reconstruction()` to perform this conversion. However, before applying it, we must ensure the sensor counts are in a normalized form. This procedure is laid out by the manufacturer. In our version, we refer to the *VEET SPD Reconstruction Guide.pdf*, version *06/05/2025*. Note that each manufacturer has to specify the method of count normalization (if any) and spectral reconstruction. In our raw data, each observation comes with a `Gain` setting that indicates how the sensor’s sensitivity was adjusted; we need to divide the raw counts by the gain to get normalized counts. LightLogR offers `normalize_counts()` for this purpose. We further need to scale by integration time (in milliseconds) and adjust depending on counts in the `Dark` sensor channel.

```
Normalize spectral sensor counts
```

```
1count.columns <- c("s415", "s445", "s480", "s515", "s555", "s590", "s630",
                      "s680", "s910", "Dark", "Clear")

2gain.ratios <-
  tibble(
    gain = c(0.5, 1, 2, 4, 8, 16, 32, 64, 128, 256, 512),
    gain.ratio =
      c(0.008, 0.016, 0.032, 0.065, 0.125, 0.25, 0.5, 1, 2, 3.95, 7.75)
  )

#normalize data:
dataVEET2 <-
  dataVEET2 |> 
3  mutate(across(c(s415:Clear), \(x) (x - Dark)/integration_time)) |>
4  normalize_counts(
5    gain.columns = rep("Gain", 11),
6    count.columns = count.columns,
7    gain.ratios
  ) |> 
8  select(-c(s415:Clear)) |>
  rename_with(~ str_remove(.x, ".normalized"))
```

1
:   Column names of variables that need to be normalized

2
:   Gain ratios as specified by the manufacturer’s reconstruction guide

3
:   Remove dark counts & scale by integration time

4
:   Function to normalize counts

5
:   All sensor channels share the gain value

6
:   Sensor channels to normalize (see 1.)

7
:   Gain ratios (see 2.)

8
:   Drop original raw count columns

In this call, we specified `gain.columns = rep("Gain", 11)` because we have 11 sensor columns that all use the same gain factor column (`Gain`). This step will add new columns (with a suffix, e.g. `.normalized`) for each spectral channel representing the count normalized by the gain. We then dropped the `raw` count columns and renamed the normalized ones by dropping `.normalized` from the names. After this, `dataVEET2` contains the normalized sensor readings for `s415, s445, ..., s940, Dark, Clear` for each time point time point.

Because we do not need this high a resolution, we will **aggregate it to a 5-minute interval** for computational efficiency. The assumption is that spectral composition does not need to be examined at every 2-second instant for our purposes, and 5-minute averages can capture the general trends while drastically reducing data size and downstream computational costs.

```
Aggregate spectral data to 5-minute intervals and mark gaps
```

```
dataVEET2 <- dataVEET2 |>
1  aggregate_Datetime(unit = "5 mins") |>
2  gap_handler(full.days = TRUE) |>
3  add_Date_col(group.by = TRUE) |>
4  remove_partial_data(Clear, threshold.missing = "1 hour")
```

1
:   Aggregate to 5-minute bins

2
:   Explicit NA for any gaps in between

3
:   Add a date identifier for grouping

4
:   remove days with more than one hour of data missing

We aggregate over 5-minute windows; within each 5-minute bin, multiple spectral readings (if present) are combined (averaged). We use one of the channels (here `Clear`) as the reference variable for `remove_partial_data` to drop incomplete days (the choice of channel is arbitrary as all channels share the same level of completeness).

Warning

Please note that `normalize_counts()` requires the `Gain` values according to the gain table. If we had aggregated the data before normalizing it, `Gain` values would have been averaged within each bin (5 minutes in this case). If the `Gain` did not change in that time, it is not an issue. Any mix of `Gain` values will lead to a `Gain` value that is not represented in the gain table. While outputs for `normalize_counts()` are not wrong in these cases, they will output `NA` if a `Gain` value is not found in the table. Thus we recommend to always normalize counts based on the raw dataset.

##### 4.3 Spectral reconstruction

For spectral reconstruction, we require a calibration matrix that corresponds to the `VEET`’s sensor channels. This matrix would typically be obtained from the device manufacturer or a calibration procedure. It defines how each channel’s normalized count relates to intensity at various wavelengths. For demonstration, the calibration matrix was provided by the manufacturer and is specific to the make and model (see Figure 13). It should not be used for research purposes without confirming its accuracy with the manufacturer.

Figure 13: Calibration matrix

```
Calibration matrica and reconstruction of spectral power distribution
```

```
1calib_mtx <-
  read_csv("data/VEET_calibration_matrix.csv",
           show_col_types = FALSE) |>
  column_to_rownames("wavelength")

dataVEET2 <-
  dataVEET2 |> 
  mutate(
  Spectrum = 
2    spectral_reconstruction(
3      pick(s415:s910),
4      calibration_matrix = calib_mtx,
5      format = "long"
  )
)
```

1
:   Import the calibration matrix and make certain `wavelength` is set a rownames

2
:   The function `spectral_reconstruction()` does not work on the level of the dataset, but has to be called within `mutate()`(or provided the data directly)

3
:   Pick the normalized sensor columns

4
:   Provide the calibration matrix

5
:   Return a long-form list column (wavelength, intensity)

Here, we use `format = "long"` so that the result for each observation is a **list-column** `Spectrum`, where each entry is a tibble^4^ containing two columns: `wavelength` and `irradiance` (one row per wavelength in the calibration matrix). In other words, each row of `dataVEET2` now holds a full reconstructed spectrum in the `Spectrum` column. The long format is convenient for further calculations and plotting. (An alternative `format = "wide"` would add each wavelength as a separate column, but that is less practical when there are many wavelengths.)

To visualize the data we will calculate the photopic illuminance based on the spectra and plot each spectrum color-scaled by their illuminance. For clarity, we reduce the data to observations within the day that has the most observations (non-`NA`).

```
Calculate photopic illuminance
```

```
data_spectra <- 
dataVEET2 |> 
1  sample_groups(order.by = sum(!is.implicit)) |>
  mutate( 
2    Illuminance = Spectrum |>
3      map_dbl(spectral_integration,
4              action.spectrum = "photopic",
5              general.weight = "auto")
  ) |> 
6  unnest(Spectrum)
data_spectra |> select(Id, Date, Datetime, Illuminance) |> distinct()
```

1
:   Keep only observations for one day (with the lowest missing intervals)

2
:   Use the spectrum,…

3
:   … call the function `spectral_integration()` for each,…

4
:   … use the brightness sensitivity function,…

5
:   … and apply the appropriate efficacy weight.

6
:   Create a long format of the data where the spectrum is unnested

```
# A tibble: 288 × 4
# Groups:   Id, Date [1]
   Id    Date       Datetime            Illuminance
   <fct> <date>     <dttm>                    <dbl>
 1 VEET  2025-06-18 2025-06-18 00:00:00        3.70
 2 VEET  2025-06-18 2025-06-18 00:05:00        3.79
 3 VEET  2025-06-18 2025-06-18 00:10:00        3.72
 4 VEET  2025-06-18 2025-06-18 00:15:00        6.90
 5 VEET  2025-06-18 2025-06-18 00:20:00        3.53
 6 VEET  2025-06-18 2025-06-18 00:25:00        3.26
 7 VEET  2025-06-18 2025-06-18 00:30:00        3.59
 8 VEET  2025-06-18 2025-06-18 00:35:00        3.49
 9 VEET  2025-06-18 2025-06-18 00:40:00        3.54
10 VEET  2025-06-18 2025-06-18 00:45:00        3.62
# ℹ 278 more rows
```

The following plot visualizes the spectra:

```
Plot spectra
```

```
data_spectra |> 
  ggplot(aes(x = wavelength,group = Datetime)) +
  geom_line(aes(y = irradiance*1000, col = Illuminance)) +
  labs(y = "Irradiance (mW/m²/nm)", 
       x = "Wavelength (nm)", 
       col = "Photopic illuminance (lx)") +
  scale_color_viridis_b(breaks = c(0, 10^(0:3))) +
  scale_y_continuous(trans = "symlog", breaks = c(0, 1, 10, 50)) +
  coord_cartesian(ylim = c(0,NA), expand = FALSE) +
  theme_minimal()
```

Figure 14: Overview of the reconstructed spectra, color-scaled by photopic illuminance (lx)

The following ridgeline plot can be used to assess **when** in the day certain spectral wavelenghts are dominant:

```
Plot spectra across the time of day
```

```
data_spectra |> 
  ggplot(aes(x = wavelength, group = Datetime)) +
  geom_ridgeline(aes(height = irradiance*1000, 
                     y = Datetime, 
                     fill = Illuminance), 
                 scale = 400, lwd = 0.1, alpha = 0.7) +
  labs(y = "Local time & Irradiance (mW/m²/nm)", 
       x = "Wavelength (nm)", 
       fill = "Photopic illuminance (lx)")+
  scale_fill_viridis_b(breaks = c(0, 10^(0:3))) +
  theme_minimal()
```

Figure 15: Overview of the reconstructed spectra by time of day, color-scaled by photopic illuminance (lx)

At this stage, the `dataVEET2` dataset has been processed to yield time-series of spectral power distributions. We can use these to compute biologically relevant light metrics. For instance, one possible metric is the proportion of power in short wavelengths versus long wavelengths.

In the main analysis, we defined **short-wavelength (blue light) content** as the integrated intensity in the 400–500 nm range, and **long-wavelength content** as the integrated intensity in a longer range (e.g. 600–700 nm), then computed the *short-to-long ratio* (“sl ratio”). Calculating these metrics is the first step of spectrum analysis in the main tutorial.

```
dataVEET2 |> 
  select(Id, Date, Datetime, Spectrum) |>    # focus on ID, date, time, and spectrum
  mutate(
    short = Spectrum |> map_dbl(spectral_integration, wavelength.range = c(400, 500)),
    long  = Spectrum |> map_dbl(spectral_integration, wavelength.range = c(600, 700)),
    `sl ratio` = short / long   # compute short-to-long ratio
  )
```

*(The cutoff of 500 nm here is hypothetical for demonstration; actual definitions might vary.)* We would then have columns `short`, `long`, and `sl_ratio` for each observation, which could be averaged per day or analyzed further. The cleaned spectral data in `dataVEET2` makes it straightforward to calculate such metrics or apply spectral weighting functions (for melatonin suppression, circadian stimulus, etc., if one has the spectral sensitivity curves).

With the `VEET` spectral preprocessing complete, we emphasize that these steps – normalizing by gain, applying calibration, and perhaps simplifying channels – are **device-specific** requirements. They ensure that the raw sensor counts are translated into meaningful physical measures (like spectral irradiance). Researchers using other spectral devices would follow a similar procedure, adjusting for their device’s particulars (some may output spectra directly, whereas others, like `VEET`, require reconstruction.

Note

Some devices may output normalized counts instead of raw counts. For example, the `ActLumus` device outputs normalized counts, while the `VEET` device records raw counts and the gain. Manufacturers will be able to speficy exact outputs for a given model and software version.

#### 5 `VEET` Data: Time of flight (distance)

In this last section, the distance data of the `VEET` device will be imported, analogous to the other modalities. The `TOF` modality contains information for up to two objects in a 8x8 grid of measurements, spanning a total of about 52° vertically and 41° horizontally. Because the `VEET` device can detect up to two objects in a given grid point, and there is a confidence value assigned to every measurement, each observation contains \(2\*2\*8\*8 = 256\) measurements.

```
Import VEET Spectral Sensor (TOF) data
```

```
path <- "data/01_VEET_L.csv.zip"
dataVEET3 <- import$VEET(path, 
                        tz = tz, 
1                        modality = "TOF",
2                        manual.id = "VEET"
                        )
```

1
:   `modality` is a parameter only the `VEET` device requires. If uncertain, which devices require special parameters, have a look a the import help page (`?import`) under the `VEET` device. Setting it to `TOF` gives us the distance modality.

2
:   As we are only dealing with one individual here, we set a manual `Id`

```
Successfully read in 304'195 observations across 1 Ids from 1 VEET-file(s).
Timezone set is US/Central.
The system timezone is Europe/Berlin. Please correct if necessary!

First Observation: 2024-06-04 15:00:36
Last Observation: 2024-06-12 08:29:43
Timespan: 7.7 days

Observation intervals: 
  Id    interval.time              n pct      
1 VEET  0s                         3 0.00099% 
2 VEET  1s                      2089 0.68673% 
3 VEET  2s                    299876 98.58051%
4 VEET  3s                      2213 0.72750% 
5 VEET  4s                         3 0.00099% 
6 VEET  6s                         1 0.00033% 
7 VEET  9s                         7 0.00230% 
8 VEET  109s (~1.82 minutes)       1 0.00033% 
9 VEET  59077s (~16.41 hours)      1 0.00033%
```

Figure 16: Overview plot of imported VEET data

```
Print the first 6 rows of the VEETs distance dataset
```

```
dataVEET3 |> head()
```

```
# A tibble: 6 × 261
# Groups:   Id [1]
  Id    Datetime            file.name     time_stamp modality conf1_0 conf1_1
  <fct> <dttm>              <chr>              <dbl> <chr>      <dbl>   <dbl>
1 VEET  2024-06-04 15:00:36 01_VEET_L.csv 1717531236 TOF           46      36
2 VEET  2024-06-04 15:00:38 01_VEET_L.csv 1717531238 TOF           29      22
3 VEET  2024-06-04 15:00:40 01_VEET_L.csv 1717531240 TOF            0       0
4 VEET  2024-06-04 15:00:42 01_VEET_L.csv 1717531242 TOF            0       0
5 VEET  2024-06-04 15:00:44 01_VEET_L.csv 1717531244 TOF            0       0
6 VEET  2024-06-04 15:00:46 01_VEET_L.csv 1717531246 TOF           59      40
# ℹ 254 more variables: conf1_2 <dbl>, conf1_3 <dbl>, conf1_4 <dbl>,
#   conf1_5 <dbl>, conf1_6 <dbl>, conf1_7 <dbl>, conf1_8 <dbl>, conf1_9 <dbl>,
#   conf1_10 <dbl>, conf1_11 <dbl>, conf1_12 <dbl>, conf1_13 <dbl>,
#   conf1_14 <dbl>, conf1_15 <dbl>, conf1_16 <dbl>, conf1_17 <dbl>,
#   conf1_18 <dbl>, conf1_19 <dbl>, conf1_20 <dbl>, conf1_21 <dbl>,
#   conf1_22 <dbl>, conf1_23 <dbl>, conf1_24 <dbl>, conf1_25 <dbl>,
#   conf1_26 <dbl>, conf1_27 <dbl>, conf1_28 <dbl>, conf1_29 <dbl>, …
```

In a first step, we condition the data similarly to the other `VEET` modalities. For computational reasons of the use cases, we remove the second object and set the interval to `10 seconds`. *Note that the next step still takes considerable computation time*.

```
Aggregate distance data to 5-second intervals and mark gaps
```

```
dataVEET3 <- 
  dataVEET3 |>
1  select(-contains(c("conf2_", "dist2_"))) |>
2  aggregate_Datetime(unit = "5 secs") |>
3  gap_handler(full.days = TRUE) |>
  add_Date_col(group.by = TRUE) |> 
  remove_partial_data(dist1_0, threshold.missing = "1 hour")
dataVEET3 |> summary_overview(dist1_0)
```

1
:   Remove the second object (for computational reasons)

2
:   Aggregate to 10-second bins

3
:   Explicit NA for any gaps

```
# A tibble: 4 × 4
  name                mean   min   max
  <chr>              <dbl> <dbl> <dbl>
1 Participants           1    NA    NA
2 Participant-days       6     6     6
3 Days ≥80% complete     6     6     6
4 Missing/Irregular      0     0     0
```

In the next step, we need to transform the `wide` format of the imported dataset into a `long` format, where each row contains exactly one observation for one grid-point.

```
Pivot distance grid from wide to long
```

```
dataVEET3 <- 
  dataVEET3 |> 
  pivot_longer(
    cols = -c(Datetime, file.name, Id, is.implicit, time_stamp, modality, Date),
    names_to = c(".value", "position"),
    names_pattern = "(conf1|conf2|dist1|dist2)_(\\d+)"
  )
```

In a final step before we can use the data in the analysis, we need to assign `x` and `y` coordinates based on the `position` column that was created when pivoting longer. Positions are counted from 0 to 63 starting at the top right and increasing towards the left, before continuing on the right in the next row below. `y` positions thus depend on the row count, i.e., how often a row of 8 values fits into the `position` column. `x` positions consequently depend on the `position` within each 8-value row. We also add an `observation` variable that increases by `+1` every time, the `position` column hits `0`. We then center both `x` and `y` coordinates to receive meaningful values, i.e., 0° indicates the center of the overall measurement cone. Lastly, we convert both confidence columns, which are scaled from 0-255 into percentages by dividing them by `255`. Empirical data from the manufacturer points to a threshold of about `10%`, under which the respective distance data is not reliable.

```
Calculate grid positions of spatial distance measurements
```

```
dataVEET3 <- 
  dataVEET3 |> 
  mutate(position = as.numeric(position),
1         y.pos = (position %/% 8)+1,
2         y.pos = scale(y.pos, scale = FALSE)*52/8,
3         x.pos = 8 - (position %% 8),
4         x.pos = scale(x.pos, scale = FALSE)*41/8,
5         observation = cumsum(position == 0),
6         across(starts_with("conf"), \(x) x/255)
         )
```

1
:   Increment the y position for every 8 steps in `position`

2
:   Center `y.pos` and rescale it to cover 52° across 8 steps

3
:   Increment the x position for every step in `position`, resetting every 8 steps

4
:   Center `x.pos` and rescale it to cover 41° across 8 steps

5
:   Increase an observation counter every time we restart with `position` at 0

6
:   Scale the confidence columns so that 255 = 100%

Now this dataset is ready for further analysis. We finish by visualizing the same observation time on different days. Note that we replace zero distance values with `Infinity`, as these indicate measurements outside the 5m measurement radius of the device.

```
Plot spatial distance grid for the same time point on each day
```

```
1extras <- list(
  geom_tile(),
  scale_fill_viridis_c(direction = -1, limits = c(0, 200),
                       oob = scales::oob_squish_any),
  scale_color_manual(values = c("black", "white")),
  theme_minimal(),
  guides(colour = "none"),
  geom_text(aes(label = (dist1/10) |> round(0), colour = dist1>1000),
            size = 2.5),
  coord_fixed(),
  labs(x = "X position (°)", y = "Y position (°)",
       fill = "Distance (cm)"))

2slicer <- function(x){seq(min((x-1)*64+1), max(x*64, by = 1))}

dataVEET3 |> 
3  slice(slicer(9530)) |>
4  mutate(dist1 = ifelse(dist1 == 0, Inf, dist1)) |>
5  filter(conf1 >= 0.1 | dist1 == Inf) |>
6  ggplot(aes(x=x.pos, y=y.pos, fill = dist1/10))+ extras +
7  facet_grid(~Datetime)
```

1
:   Set visualization parameters

2
:   Allows to choose an observation

3
:   Choose a particular observation

4
:   Replace 0 distances with Infinity

5
:   Remove data that has less than 10% confidence

6
:   Plot the data

7
:   Show one plot per day

Figure 17: Example observations of the measurement grid at 1:14 p.m. for each measurement day. Text values show distance in cm. Empty grid points show values with low confidence. Zero-distance values were replaced with infinite distance and plotted despite low confidence.

As we can see from the figure, different days have - at a given time - a vastly different distribution of distance data, and measurement confidence (values with confidence < 10% are removed)

##### 5.1 Removing non-wear times

Same as for the `ALS` modality, we can remove times of non-wear - we can even use the same `wear_data` set for it:

```
Removing non-wear observations from distance modality
```

```
dataVEET3 <- 
  dataVEET3 |> 
  group_by(Id) |> 
  add_states(wear_data)

dataVEET3 <- 
  dataVEET3 |> 
  mutate(dist1 = ifelse(wear, dist1, NA)) |> 
  group_by(Id, Date)
```

#### 6 Saving the Cleaned Data

After executing all the above steps, we have three cleaned data frames in our R session:

- **`dataCC`** – the processed `Clouclip` dataset (5-second intervals, with distance and lux, including NA for gaps and sentinel statuses).
- **`dataVEET`** – the processed `VEET` ambient light dataset (5-second intervals, illuminance in lux, with gaps filled).
- **`dataVEET2`** – the processed `VEET` spectral dataset (5-minute intervals, each entry containing a spectrum or derived spectral metrics).
- **`dataVEET3`** – the processed `VEET` distance dataset (5-second intervals, each entry containing the distance of up to two objects in the 8x8 grid).

For convenience and future reproducibility, we will save these combined results to a single R data file. Storing all cleaned data together ensures that any analysis can reload the exact same data state without re-running the import and cleaning (which can be time-consuming for large raw files).

```
Save preprocessed files
```

```
1if (!dir.exists("data/cleaned")) dir.create("data/cleaned", recursive = TRUE)
2save(dataCC, dataVEET, dataVEET2, dataVEET3, file = "data/cleaned/data.RData")
```

1
:   Create directory for cleaned data if it doesn’t exist

2
:   Save all cleaned datasets into one .RData file

The above code creates (if necessary) a folder `data/cleaned/` and saves a single RData file (`data.RData`) containing the three objects. To retrieve them later, one can use `<- load("data/cleaned/data.RData")`, which will return the objects into the environment. This single-file approach simplifies sharing and keeps the cleaned data together.

In summary, this supplement has walked through the full preprocessing pipeline for two example devices. We began by describing the raw data format for each device and then demonstrated how to import the data with correct time zone settings. We handled device-specific quirks like sentinel codes (for `Clouclip`) and multiple modalities with gain normalization (for `VEET`). We showed how to detect and address irregular sampling, how to explicitly mark missing data gaps to avoid analytic pitfalls, and how to reduce data granularity via rounding or aggregation when appropriate. Throughout, we used functions from LightLogR in a tidyverse workflow, aiming to make the steps clear and modular. By saving the final cleaned datasets, we set the stage for the computation of visual experience metrics such as working distance, time in bright light, spectral composition ratios, as presented in the main tutorial. We hope this detailed tutorial empowers researchers to adopt similar pipelines for their own data, facilitating reproducible and accurate analyses of visual experience.

#### Footnotes

1. A *sentinel value* is a special placeholder value used in data recording to signal a particular condition. It does not represent a valid measured quantity but rather acts as a marker (for example, “device off” or “value out of range”).↩︎
2. **tibble** are data.tables with tweaked behavior, ideal for a tidy analysis workflow. For more information, visit the documentation page for tibbles↩︎
3. **tibble** are data.tables with tweaked behavior, ideal for a tidy analysis workflow. For more information, visit the documentation page for tibbles↩︎
4. **tibble** are data.tables with tweaked behavior, ideal for a tidy analysis workflow. For more information, visit the documentation page for tibbles↩︎


###### Source Code

```
---
title: "Supplement 1"
subtitle: "Analysis of human visual experience data"
author:
  - name: "Johannes Zauner"
    id: JZ
    affiliation: 
      - Technical University of Munich, Germany
      - Max Planck Institute for Biological Cybernetics, Tübingen, Germany
    orcid: "0000-0003-2171-4566"
    corresponding: true
   
  - name: "Aaron Nicholls"
    affiliation: "Reality Labs Research, USA"
    orcid: "0009-0001-6683-6826"
  - name: "Lisa A. Ostrin"
    affiliation: "University of Houston College of Optometry, USA"
    orcid: "0000-0001-8675-1478"
  - name: "Manuel Spitschan"
    affiliation: 
      - Technical University of Munich, Germany
      - Max Planck Institute for Biological Cybernetics, Tübingen, Germany
      - Technical University of Munich Institute of advanced study (TUM-IAS), Munich, Germany
    orcid: "0000-0002-8572-9268"
doi: 10.5281/zenodo.16566014
format:
  html:
    toc: true
    self-contained: true
    include-in-header:
      - text: |
          <style>
          .quarto-notebook .cell-container .cell-decorator {
            display: none;
          }
          </style>
    number-sections: true
    code-tools: true
    html-math-method:
      method: mathjax
      url: "https://cdnjs.cloudflare.com/ajax/libs/mathjax/2.7.9/latest.js?config=TeX-MML-AM_CHTML"
    theme:
      - cosmo
      - brand
    css: styles.css
    link-external-newwindow: true
    # comments:
    #   hypothesis: true
    format:
    code-link: true
    code-line-numbers: true
    toc-location: left
    toc-depth: 3
    toc-expand: 2

bibliography: references.bib
lightbox: true
execute:
  warning: false
  error: false
  echo: true
---

## Abstract {.unnumbered}

This supplementary document provides a detailed, step-by-step tutorial on importing and preprocessing raw data from two wearable devices: `Clouclip` and `VEET`. We describe the structure of the raw datasets recorded by each device and explain how to parse these data, specify time zones, handle special sentinel values, clean missing observations, regularize timestamps to fixed intervals, and aggregate data as needed. All original R code from the [main tutorial](index.qmd) is shown here for transparency, with additional guidance intended for a broad research audience. We demonstrate how to detect gaps and irregular sampling, convert implicit missing periods into explicit missing values, and address device-specific quirks such as the `Clouclip`’s use of sentinel codes for “sleep mode” and “out of range” readings. Special procedures for processing the `VEET`’s rich spectral data (e.g. normalizing sensor counts and reconstructing full spectra from multiple sensor channels) are also outlined. Finally, we show how to save the cleaned datasets from both devices into a single R data file for downstream analysis. This comprehensive walkthrough is designed to ensure reproducibility and to assist researchers in understanding and adapting the data pipeline for their own visual experience datasets.

## Introduction

Wearable sensors like the `Clouclip` and the Visual Environment Evaluation Tool (`VEET`) produce high-dimensional time-series data on viewing distance and light exposure. Proper handling of these raw data is essential before any analysis of visual experience metrics. In the [main tutorial](index.qmd), we introduce an analysis pipeline using the open-source R package LightLogR [@Zauner2025JOpenSourceSoftw] to calculate various distance and light exposure metrics. Here, we present a full account of the data import and preparation steps as a supplement to the methods, with an emphasis on clarity for researchers who may be less familiar with data processing in R. @fig-fc_supplement shows the main pre-processing steps and how they relate.

{{< include _flowchart_supplement.qmd >}}

We use example datasets from a `Clouclip` device [@Wen2021ActaOphtalmol; @Wen2020bjophthalmol] and from a `VEET` device [@Sah2025OphtalmicPhysiolOpt] (both provided in the accompanying [repository](https://github.com/tscnlab/ZaunerEtAl_JVis_2026)). The `Clouclip` is a glasses-mounted sensor that records working distance (distance from eyes to object, in cm) and ambient illuminance (in lux) at 5-second intervals. The `VEET` is a head-mounted multi-modal sensor that logs ambient light and spectral information (along with other data like motion and distance via a depth sensor) - in this exemplary case at 2-second intervals. A single week of continuous data illustrates the contrast in complexity: approximately 1.6 MB for the `Clouclip`’s simple two-column output versus up to 270 MB for the `VEET`’s multi-channel output (due to its higher sampling rate and richer sensor modalities).

In the following sections, we detail how these raw data are structured and how to import and preprocess them using LightLogR. We cover device-specific considerations such as file format quirks and sensor range limitations, as well as general best practices like handling missing timestamps and normalizing sensor readings. All code blocks can be executed in R (with the required packages loaded) to reproduce the cleaning steps. The end result will be clean, regularized datasets (`dataCC` for Clouclip, `dataVEET` for VEET light data, `dataVEET2` for VEET spectral data, and `dataVEET3` for VEET distance) ready for calculating visual experience metrics. We conclude by saving these cleaned datasets into a single file for convenient reuse.

## `Clouclip` Data: Raw Structure and Import

The `Clouclip` device exports its data as a text file (not a true Excel file despite sometimes using an .xls extension), which is actually tab-separated values. See @fig-Clouclip_file for the raw file.

![Screenshot of the first rows of the `Clouclip` export file format as seen in a text editor](assets/Clouclip_fileformat.png){#fig-Clouclip_file width=35%}

Each record corresponds to one timestamped observation (nominally every 5 seconds) and includes two measured variables: **distance** and **illuminance**. In the sample dataset (`Sample_Clouclip.csv` provided), the columns are:

-   `Date` – the date and time of the observation (in the device’s local time, here one week in 2021).

-   `Dis` – the viewing distance in centimeters.

-   `Lux` – the ambient illuminance in lux.

For example, a raw data line might look like:

```         
2021-07-01 08:00:00 45  320
```

indicating that at *2021-07-01 08:00:00* local time, the device recorded a working distance of 45 cm and illuminance of 320 lx. The Clouclip uses special **sentinel values**[^1] in these measurement columns to denote certain device states. Specifically, a distance (`Dis`) value of **204** is a code meaning the object is *out of the sensor’s range*, and a value of **-1** in either `Dis` or `Lux` indicates the device was in *sleep mode* (not actively recording). During normal operation, distance measurements are limited by the device’s range, and illuminance readings are positive lux values. Any sentinel codes in the raw file need to be handled appropriately, as described below.

[^1]: A *sentinel value* is a special placeholder value used in data recording to signal a particular condition. It does not represent a valid measured quantity but rather acts as a marker (for example, “device off” or “value out of range”).

We will use `LightLogR`’s built-in import function for `Clouclip`, which automatically reads the file, parses the timestamps, and converts sentinel codes into a separate status annotation. To begin, we load the necessary libraries and import the raw `Clouclip` dataset:

```{r}
#| label: setup
#| output: false
#| filename: Load required packages
library(tidyverse) #<1>
library(LightLogR) #<2>
library(gt) #<3>
library(ggridges) #<4>
library(downlit) #<5>
library(magick) #<5>
```

1. For tidy data science
2. Wearable analysis package
3. For great tables
4. For ridgeline plots
5. These packages are not used, but are needed for dependencies

```{r}
#| label: fig-importCC
#| filename: Import of Clouclip data
#| fig-cap: "Overview plot of imported Clouclip data"
#| fig-height: 2
#| fig-width: 6
path <- "data/Sample_Clouclip.csv" #<1>
tz   <- "US/Central"   #<2>
dataCC <- import$Clouclip(path, tz = tz, manual.id = "Clouclip") #<3>
```

1. Define file path
2. Time zone in which device was recording (e.g., US Central Time)
3. Import Clouclip data

In this code, `import$Clouclip()` reads the tab-delimited file and returns a **tibble**[^2] (saved in the variable `dataCC`) containing the data. We specify `tz = "US/Central"` because the device’s clock was set to U.S. Central time; this ensures that the `Datetime`values are properly interpreted with the correct time zone. The argument `manual.id = "Clouclip"` simply tags the dataset with an identifier (useful if combining data from multiple devices).

[^2]: **tibble** are data.tables with tweaked behavior, ideal for a tidy analysis workflow. For more information, visit the documentation page for [tibbles](https://tibble.tidyverse.org/index.html)

During import, LightLogR automatically handles the `Clouclip`’s sentinel codes. The `Date` column from the raw file is parsed into a `POSIXct` date-time (`Datetime`) with the specified time zone. The `Lux` and `Dis` columns are read as numeric, but any occurrences of **-1** or **204** are treated specially: these are replaced with `NA` (missing values) in the numeric columns, and corresponding status columns `Lux_status` and `Dis_status` are created to indicate the reason for those `NA` values. For example, if a `Dis` value of *204* was encountered, that row’s `Dis` will be set to NA and `Dis_status` will contain `"out_of_range"`; if `Lux` or `Dis` was -1, the status is `"sleep_mode"`. We will set all other readings to  `"operational"` (meaning the device was functioning normally at that time) for visualisation purposes.

```{r}
#| filename: Print the first 6 rows of the Clouclip dataset
dataCC |> head()
```


After import, it is good practice to get an overview of the data. The import function by default prints a brief summary (and generates an overview plot of the timeline) showing the number of observations, the time span, and any irregularities or large gaps. In our case, the `Clouclip` summary indicates the data spans one week and reveals that there are **irregular intervals** in the timestamps. This means some observations do not occur exactly every 5 seconds as expected. We can programmatically check for irregular timing:

```{r}
#| filename: Check if data are on a perfectly regular 5-second schedule
dataCC |> has_irregulars()
```

If the result is `TRUE`, it confirms that the time sequence has deviations from the regular interval. Indeed, our example dataset has many small timing irregularities and gaps (periods with no data). Understanding the pattern of these missing or irregular readings is important. We can visualize them using a gap plot:

```{r}
#| label: fig-irregular
#| filename: Plot gaps and irregular timestamps for Clouclip data
#| fig-height: 12
#| warning: false
#| fig-cap: "Visualization of gaps and irregular data. Black traces show available data. Red shaded areas show times of missing data. Red dots show instances where observations occur off the regular interval from start to finish, i.e., irregular data."
y.label <- "Distance (cm)"
dataCC |> gg_gaps(Dis, #<1>
                  include.implicit.gaps = FALSE,  #<2>
                  show.irregulars = TRUE,         #<3>
                  y.axis.label = y.label,
                  group.by.days = TRUE, col.irregular = alpha("red", 0.03)
                  ) + labs(title = NULL)
```

1. Basing the gap-figure on the `Dis` distance variable
2. Only show missing observations (`NA` values)
3. Highlight irregular timing

In @fig-irregular, time periods where data are missing appear as red-shaded areas, and any off-schedule observation times are marked with red dots. The `Clouclip` example shows extensive gaps (red blocks) on certain days and irregular timing on all days except the first and last. These irregular timestamps likely arise from the device’s logging process (e.g. slight clock drift or buffering when the device was turned on/off). Such issues must be addressed before further analysis.

When faced with irregular or gapped data, we recommend a few strategies:

-   *Remove leading/trailing segments that cause irregularity.* For example, if only the first day is regular and subsequent days drift, one might exclude the problematic portion using date filters (see [filter_Date() / filter_Datetime()](https://tscnlab.github.io/LightLogR/reference/filter_Datetime.html) in LightLogR).

-   *Round timestamps to the nearest regular interval.* This relabels slightly off-schedule times back to the 5-second grid (using [cut_Datetime()](https://tscnlab.github.io/LightLogR/reference/cut_Datetime.html) with a 5-second interval), provided the deviations are small and this rounding won’t create duplicate timestamps.

-   *Aggregate to a coarser time interval.* For instance, grouping and averaging data into 1-minute bins with [aggregate_Datetime()](https://tscnlab.github.io/LightLogR/reference/aggregate_Datetime.html) can mask irregularities at finer scales, at the cost of some temporal resolution.

In this case, the deviations from the 5-second schedule are relatively minor. We choose to **round the timestamps to the nearest 5 seconds** to enforce a uniform sampling grid, which simplifies downstream gap handling. We further add a separate date column for convenience:

```{r}
#| filename: Regularize timestamps by rounding to nearest 5-second interval
dataCC <- dataCC |>
  cut_Datetime("5 secs", New.colname = Datetime) |>  #<1>
  add_Date_col(group.by = TRUE)                     #<2>
```

1. Round times to 5-second bins
2. Add a Date column for grouping by day

After this operation, all `Datetime` entries in `dataCC` align perfectly on 5-second boundaries (e.g. **08:00:00**, **08:00:05**, **08:00:10**, etc.). We can verify that no irregular intervals remain by re-running `has_irregulars()`:

```{r}
#| filename: Re-check if data are on a perfectly regular 5-second schedule
dataCC |> has_irregulars()
```

Next, we want to quantify the missing data. LightLogR distinguishes between **explicit missing values** (actual `NA`s in the data, possibly from sentinel replacements or gaps we have filled in) and **implicit missing intervals** (time points where the device *should* have a reading but none was recorded, and we have not yet filled them in). See @fig-gaps for a visual aid to these terms. Initially, many gaps are still implicit (between the first and last timestamp of each day). 

::: {#fig-gaps layout-ncol=2}

![Terminology of gaps and irregular data in LightLogR](assets/gap_terminology.png)

![Handling of gaps with LightLogR's `gap_handler()` function](assets/gap_handler.png)

Gaps and irregular data

:::

We can generate a gap summary table:

```{r}
#| label: tbl-gaps
#| tbl-cap: "Summary of missing and observed data for the Clouclip device"
#| filename: Summarize observed vs missing data by day for distance
dataCC |> gap_table(Dis, Variable.label = "Distance (cm)") |>
  cols_hide(contains("_n"))   #<1>
```

1. Hide absolute counts for brevity in output

This summary (@tbl-gaps) breaks down, for each day, how much data is present vs. missing. It reports the total duration of recorded data and the duration of gaps. After rounding the times, there are no irregular timestamp issues, but we see substantial **implicit gaps** — periods where the device was not recording (e.g., overnight when the device was likely not worn or was in sleep mode). Notably, the first and last days of the week have very little data (less than 1 hour each), since they probably represent partial recording days (the trial started and ended on those days).

To prepare the dataset for analysis, we will convert all those implicit gaps into explicit missing entries, and remove days that are mostly incomplete. Converting implicit gaps means inserting rows with `NA` for each missing 5-second slot, so that the time series becomes continuous and explicit about missingness (see @fig-gaps). We use `gap_handler()` for this, and then drop the nearly-empty days:

```{r}
#| filename: Convert implicit gaps to explicit NA gaps, and drop days with <1 hour of data
dataCC <- dataCC |> 
  mutate(across(c(Lux_status, Dis_status), ~ replace_na(.x, "operational"))) |> #<1>
  gap_handler(full.days = TRUE) |> #<2> 
  remove_partial_data(Dis, threshold.missing = "-1 hour") #<3>
```

1. First ensure that status columns have an "operational" tag for non-missing periods
2. Fill in all implicit gaps with explicit NA rows (for full days range)
3. Remove any day that has less than 1 hour of recorded data

After these steps, `dataCC` contains continuous 5-second timestamps for each day that remains. We chose a threshold of “-1 hours” to remove days with less than one hour of data, which in this dataset removes the first and last (partial) days. The cleaned `Clouclip` data now covers six full days with bouts of continuous wear.

It is often helpful to double-check how the sentinel values and missing data are distributed over time. We can visualize the distance time-series with status annotations and day/night periods:

```{r}
#| fig-height: 8
#| warning: false
#| filename: Visualize observations and sentinel states
#| label: fig-state
#| fig-cap: "Distance measurements across days. Blue, grey and yellow-colored bars at the bottom of each day show sentinel states of the device. Blue indicates an operational status, grey sleep mode (not recording), and yellow an out of range measurement. Shaded areas in the main plot show nighttime from civil dusk until dawn, which are calculated based on the recording date and geographic coordinates"
coordinates <- c(29.75, -95.36) #<1>
dataCC |> 
  fill(c(Lux_status, Dis_status), .direction = "downup") |> #<2>
  gg_day(y.axis = Dis, geom = "line", y.axis.label = y.label) |> #<3>
  gg_state(Dis_status, aes_fill = Dis_status, ymin = 0, ymax = 0.5, alpha = 1) |> #<4>
  gg_photoperiod(coordinates, alpha = 0.1) + #<5>
  theme(legend.position = "bottom")
```

1. Setting coordinates for Houston, Texas (recording location)
2. Retain status markers until a new marker arrives
3. Create the basic plot
4. Add the status times
5. Add the photoperiod (day/night)

In this plot, blue segments indicate times when the Clouclip was *operational* (actively measuring), grey segments indicate the device in *sleep mode* (no recording, typically at night), and yellow segments indicate *out-of-range* distance readings. The red outlined regions show nighttime (from civil dusk to dawn) based on the given latitude/longitude and dates. As expected, most of the grey “sleep” periods align with night hours, and we see a few yellow spans when the user’s viewing distance exceeded the device’s range (e.g. presumably when no object was within 100 cm, such as when the user looked into the distance). At this stage, the Clouclip dataset `dataCC` is fully preprocessed: all timestamps are regular 5-second intervals, missing data are explicitly marked, extraneous partial days are removed, and sentinel codes are handled via the status columns. The data are ready for calculating daily distance and light exposure metrics (as done in the [main tutorial](index.qmd)’s Results).

## `VEET` Data: Ambient Light (Illuminance) Processing

The **VEET** device [@Sah2025OphtalmicPhysiolOpt, @Sullivan2024] is a more complex logger that records multiple data modalities in one combined file. Its raw data file contains interleaved records for different sensor types, distinguished by a “modality” field. We focus first on the ambient light sensor modality (abbreviated **ALS**), which provides broad-spectrum illuminance readings (lux) and related information like sensor gains and flicker, recorded every 2 seconds. Later we will import the spectral sensor modality (**PHO**) for spectral irradiance data, and the time-of-flight modality (**TOF**) for distance data.

In the `VEET`’s export file, each line includes a timestamp and a modality code, followed by fields specific to that modality. Importantly, this means that the `VEET` export is not rectangular, i.e., tabular (see @fig-VEET_file). This makes it challenging for many import functions that expect the equal number of columns in every row, which is not the case in this instance. For the **ALS modality**, the relevant columns include a high-resolution timestamp (in Unix epoch format), integration time, UV/VIS/IR sensor gain settings, raw UV/VIS/IR sensor counts, a flicker measurement, and the computed illuminance in lux. For example, the ALS data columns are named: `time_stamp`, `integration_time`, `uvGain`, `visGain`, `irGain`, `uvValue`, `visValue`, `irValue`, `Flicker`, and `Lux`.

![Screenshot of the first rows of the `VEET` export file format as seen in a text editor](assets/VEET_fileformat.png){#fig-VEET_file}

For the **PHO (spectral) modality**, the columns include a timestamp, integration time, a general `Gain` factor, and nine sensor channel readings covering different wavelengths (with names like `s415, s445, ..., s680, s940`) as well as a `Dark` channel, a broadband channel `Clear`, and another broadband channel for flicker detection (`FD`). In essence, the `VEET`’s spectral sensor captures light in several wavelength bands (from \~415 nm up to 940 nm) rather than outputting a single lux value like the ambient light sensor does (*PHO*).

To import the `VEET` ambient light data, we again use the LightLogR import function, specifying the `ALS` modality. The raw VEET data in our example is provided as a zip file (`01_VEET_L.csv.zip`) containing the logged data for one week. We do the following:

```{r}
#| label: fig-VEET-overview
#| fig-cap: "Overview plot of imported VEET data"
#| fig-height: 2
#| fig-width: 6
#| filename: Import VEET Ambient Light Sensor (ALS) data
path <- "data/01_VEET_L.csv.zip"
tz   <- "US/Central"
dataVEET <- import$VEET(path, tz = tz, modality = "ALS", manual.id = "VEET") #<1>
```

1. In difference to the `Clouclip` file, we simply respecify the device type with `import$VEET(...)`, but must also provide a `modality` argument.

::: {.callout-note}

We get one warning as a single time stamp could not be parsed into a datetime - with \~300k observations, this is not an issue.

:::

This call reads in only the lines corresponding to the `ALS` modality from the `VEET` file. The result `dataVEET` is a tibble[^2] with columns such as `Datetime` (parsed from the `time_stamp`  to POSIXct in US/Central time), `Lux` (illuminance in lux), `Flicker`, and the various sensor gains/values. Columns we don't need for our analysis, like the modality code or file name, are also included but can be ignored or removed. From the import summary, we learn that the `VEET` light data, like the `Clouclip`, also exhibits irregularities and gaps. (The device nominally records every 2 seconds, but timing may drift or pause when not worn.)

```{r}
#| filename: Print the first 6 rows of the VEETs light dataset
dataVEET |> head()
```


To make the VEET light data comparable to the Clouclip’s and to simplify analysis, we choose to **aggregate the VEET illuminance data to 5-second intervals**. This slight downsampling will both reduce data volume and remove irregularities.

```{r}
#| filename: Aggregate VEET light data to 5-second intervals and mark gaps
dataVEET <- dataVEET |>
  aggregate_Datetime(unit = "5 seconds") |>  #<1>
  gap_handler(full.days = TRUE) |>          #<2>
  add_Date_col(group.by = TRUE) |>          #<3>
  remove_partial_data(Lux, threshold.missing = "1 hour") #<4>
```

1. Resample to 5-sec bins (e.g. average Lux over 2-sec readings)
2. Fill in implicit gaps with NA rows
3. Add Date column for daily grouping
4. Remove participant days with more than one hour of missing data

First, `aggregate_Datetime(unit = "5 seconds")` combines the high-frequency 2-second observations into 5-second slots. By default, this function will average numeric columns like `Lux` over each 5-second period (and have sensible defaults for strings or categorical data). All of these *data type handlers* can be changed with the function call. The result is that `dataVEET` now has a reading every 5 seconds (or an NA if none were present in that window). Next, `gap_handler(full.days = TRUE)` inserts explicit NA entries for any 5-second timestamp that had no data within the continuous span of the recording. Then we add a `Date` column for grouping, and finally we remove days with more than 1 hour of missing data (using a more strict criterion as we did for `Clouclip`). According to the gap summary (@tbl-gaps2), this leaves six full days of `VEET` light data with good coverage, after dropping the very incomplete start/end days.

We can inspect the missing-data summary for the `VEET` illuminance data:

```{r}
#| label: tbl-gaps2
#| tbl-cap: "Summary of missing and observed data for the VEET device, light modality"
dataVEET |> gap_table(Lux, "Illuminance (lx)") |> cols_hide(contains("_n"))
```

@tbl-gaps2 shows, for each retained day, the total recorded duration and the duration of gaps. The `VEET` device, like the `Clouclip`, was not worn continuously 24 hours per day, so there are nightly gaps of wear time (when the device was likely off the participant), but not of recordings. After our preprocessing, any implicit gaps are represented as explicit missing intervals. The `VEET`’s time sampling was originally more frequent, but by aggregating to 5 s we have ensured a uniform timeline akin to the `Clouclip`’s.

At this point, the `dataVEET` object (for illuminance) is cleaned and ready for computing light exposure metrics. For example, one could calculate daily mean illuminance or the duration spent above certain light thresholds (e.g. “outdoor light exposure” defined as \>1000 lx) using this dataset. Indeed, basic summary tables in the [main tutorial](index.qmd) illustrate the highly skewed nature of light exposure data and the calculation of outdoor light metrics. We will not repeat those metric calculations here in the supplement, as our focus is on data preprocessing; however, having a cleaned, gap-marked time series is crucial for those metrics to be accurate.

### Identifying non-wear times

Not all recorded time points of the `VEET` can be considered as `wear time`. `Non-wear` times should generally be labelled in the wearable time series, if possible. There are several ways how `Non-wear` can be classified [@Zauner2025npj]:

1. A device detects `non-wear` and does not record any measurements during non-wear times, or has a variable in the exported data to denote non-wear times. The `Clouclip` is an example here with the `sleep mode` sentinel state.

2. Record non-wear separately, e.g. with a wear-log or diary. This is easy to implement, but puts a higher burden on participants and research staff, and can be error prone: forgotten, or incorrect data entries can mis-classify wear and non-wear times. For an implementation on how to remove non-wear times based on such a log, see, e.g., this [tutorial](https://tscnlab.github.io/LightLogR_webinar/advanced_01_a_day_in_daylight-live.html). These information can also be derived from contextual information. For example, the `VEET` records charging times in a separate file `log.csv`, which can be considered as definitive non-wear times.

3. Instruct participants with a certain behavior when removing the device to ease automated detection of non-wear times in the data. These include putting a device in a black opaque bag, so that sensor readings for light are zero, or pressing an event button (if one exists). Up- and downsides are similar to the previous option, but the measures are usually harder to decipher in the data: when a buttonpress was forgotten, does that mess up the automated detection moving forward, which expects two presses per non-wear period (at beginning and at start). When there is zero lux during the day - was it in a black bag, or was this person just in a dark environment?

4. Automated detection by an algorithm. There are several possibilities to try to classify non-wear times based on the available data from a wearable. E.g., if there is an activity tracker, that can be used to determine periods of inactivity, which could indicate non-wear. If only light data is available, the standard deviation (or coefficient of variance) of light in a moving window of a few minutes can be used to determine low variance, which also indicates inactivity. Usually a histogram of values shows a spike in those periods of inactivity.

::: {.callout-note}

#### How to handle non-wear intervals

Whether classified non-wear intervals should be removed, i.e., observations set to `NA`, or not should be a deliberate decision by the researcher. If it can be assumed that non-wear times misrepresent the measurement environment of the participant, then these observations are best set to `NA`. If, on the other hand, they might be somewhat representative, then it might be enough to label these times but still include them in analyses. Usually, this requires special instructions to the participants for non-wear. E.g., instructions for bed-time could include setting the device on the night stand facing into the room (i.e., sensors not obstructed). While this would not represent distance well, it can be considered as sufficient for ambient light measurements. 

If datasets are large enough, i.e., collect data from a long enought period, `non-wear` has a diminishing effect, at least as daily metrics of light are concerned. E.g., in a monthlong study of 39 participants from Switzerland and Malaysia, [@Biller2025] found that based on a sensitivity anaylsis up to 6 hours of data could be missing **every day**, and still the daily metrics across that month would not significantly change. This assumption only holds when non-wear times are not systematic, i.e., at the same time each day.

:::

We will perform a simple variant of the third option, and look at the `VEET`'s `IMU` modality.

```{r}
#| fig-height: 2
#| fig-width: 6
#| filename: Import VEET activity (IMU) data
path <- "data/01_VEET_L.csv.zip"
tz   <- "US/Central"
dataIMU <- 
  import$VEET(path, tz = tz, modality = "IMU", manual.id = "VEET",
              remove_duplicates = TRUE, #<1>
              silent = TRUE) #<2>
```

1. Some rows in the data are duplicated. These will be removed during import
2. In this instance, we skip the import summary

To get a quick feeling for the data, we visualize the activity variable for the x axis in @fig-IMU

```{r}
#| fig-height: 2
#| label: fig-IMU
#| fig-width: 14
#| filename: Plotting the IMU timeline
#| fig-cap: "Timeline of the activity sensor (x direction)"
dataIMU |> 
  gg_days(ax, 
          aes_col = abs(ax) < 1, 
          group = consecutive_id(abs(ax) < 1),
          y.axis.label = "ax"
          )
```

`ax` seems to vary between ±10. In general, the periods of inactivity seem to lie within a band of ±1. However, simply choosing this band does not eliminate false detections during times of high movement. Some data transformations can make these more clear:

```{r}
#| label: fig-IMU2
#| fig-width: 14
#| filename: Plotting a transformed IMU timeline
#| fig-cap: "Timeline of the activity sensor (x direction) - transformed to distinguish times of low activity"
dataIMU |> 
  aggregate_Datetime(
    "5 mins",
    numeric.handler = sd
  ) |> 
  pivot_longer(cols = c(ax, ay, az)) |> 
  group_by(name) |> 
  gg_days(value, 
          aes_col = value < 0.05,
          group = consecutive_id(value < 0.05),
          y.axis.label = "activity axes"
          )
```

What @fig-IMU2 shows is that the 5-minute standard deviation of the activity channels (x, y, and z direction) allows a pretty stable distinction between periods of high and low activity, with a threshold of **0.05**. We can us this to create a wear column. As the choice of channel does not seem to matter much, we will use `ax`.

```{r}
#| filename: Calculating times of wear and non-wear
wear_data <- 
dataIMU |> 
  aggregate_Datetime("5 mins",numeric.handler = sd) |> #<1>
  mutate(wear = ax > 0.05) |> #<1>
  select(Id, Datetime, wear) |> #<1>
  extract_states(wear) #<2>
wear_data |> #<3>
  ungroup(Id) |> #<3>
  summarize_numeric(remove = c("start", "end", "epoch")) |> #<3>
  gt() #<3>
wear_data <- wear_data |> select(Id, wear, start, end) #<2>
```

1. Calculation of the wear-variable
2. Extracting the start and end times of wear and non-wear
3. Summarizing wear and non-wear times in a table

We see that most of the recorded timespan is actually non-wear (including sleep), with an average wear time of 2.6 hours.

Let's add this data to the `ALS` dataset and visualize it in @fig-nonwear

```{r}
#| fig-height: 4
#| label: fig-nonwear
#| fig-width: 14
#| filename: Plotting non-wear for light
#| fig-cap: "Wear times for light data, shown as red bars"
dataVEET <- 
dataVEET |> 
  group_by(Id) |> 
  add_states(wear_data)

dataVEET |> 
  gg_days(Lux) |> 
  gg_states(wear, ymax = 0, alpha = 1, fill = "red")
```

This looks sensible, e.g., when we look at noon of 7 June 2024. We can remove these observations based on the `wear` column to make our summaries more valid.

```{r}
#| filename: Removing non-wear observations from light modality
#| warning: false
dataVEET <- dataVEET |> mutate(Lux = ifelse(wear, Lux, NA)) |> group_by(Id, Date)
```

## `VEET` Data: Spectral Data Processing

### Import

In addition to broad-band illuminance and distance, the `VEET` provides **spectral sensor data** through its `PHO` modality. Unlike illuminance, the spectral data are not given as directly interpretable radiometric metrics but rather as raw sensor counts across multiple wavelength channels, which require conversion to reconstruct a spectral power distribution. In our analysis, spectral data allow us to compute metrics like the relative contribution of short-wavelength (blue) light versus long-wavelength light in the participant’s environment. Processing this spectral data involves several necessary steps.

First, we import the spectral modality from a second `VEET` file. This time we need to extract the lines marked as `PHO`. We will store the spectral dataset in a separate object `dataVEET2` so as not to overwrite the `dataVEET` illuminance data in our R session:

```{r}
#| label: fig-VEET-overview2
#| fig-cap: "Overview plot of imported VEET data"
#| fig-height: 2
#| fig-width: 6
#| filename: Import VEET Spectral Sensor (PHO) data
path <- "data/02_VEET_L.csv.zip"
dataVEET2 <- import$VEET(path, tz = tz, modality = "PHO", manual.id = "VEET")
```

```{r}
#| filename: Print the first 6 rows of the VEETs spectral dataset
dataVEET2 |> head()
```


After import, `data` contains columns for the timestamp (`Datetime`), `Gain` (the sensor gain setting), and the nine spectral sensor channels plus a clear channel. These appear as numeric columns named `s415, s445, ..., s940, Dark, Clear`. Other columns are also present but not needed for now. The spectral sensor was logging at a 2-second rate. It is informative to look at a snippet of the imported spectral data before further processing. @tbl-PHO shows three rows of `data` after import (before calibration), with some technical columns omitted for brevity:

```{r}
#| label: tbl-PHO
#| filename: Table overview of spectral sensor data
#| tbl-cap: "Overview of the spectral sensor import from the VEET device (3 observations). Each row corresponds to a 2-second timestamp (Datetime) and shows the raw sensor readings for the spectral channels (s415–s940, Dark, Clear). All values are in arbitrary sensor units (counts). Gain values and integration_time are also relevant for each interval, depending on the downstream computation."
dataVEET2 |> 
  slice(6000:6003) |> 
  select(-c(modality, file.name, time_stamp)) |> 
  gt() |> 
  fmt_number(s415:Clear) 
```

### Spectral calibration

Now we proceed with **spectral calibration**. The `VEET`’s spectral sensor counts need to be converted to physical units (spectral irradiance) via a calibration matrix provided by the manufacturer. For this example, we assume we have a calibration matrix that maps all the channel readings to an estimated spectral power distribution (SPD). The LightLogR package provides a function `spectral_reconstruction()` to perform this conversion. However, before applying it, we must ensure the sensor counts are in a normalized form. This procedure is laid out by the manufacturer. In our version, we refer to the [*VEET SPD Reconstruction Guide.pdf*](https://projectveet.com/wp-content/uploads/VEET-Spectral-Power-Distribution-Reconstruction-Guide.pdf){target="_blank"}, version *06/05/2025*. Note that each manufacturer has to specify the method of count normalization (if any) and spectral reconstruction. In our raw data, each observation comes with a `Gain` setting that indicates how the sensor’s sensitivity was adjusted; we need to divide the raw counts by the gain to get normalized counts. LightLogR offers `normalize_counts()` for this purpose. We further need to scale by integration time (in milliseconds) and adjust depending on counts in the `Dark` sensor channel.

```{r}
#| filename: Normalize spectral sensor counts
count.columns <- c("s415", "s445", "s480", "s515", "s555", "s590", "s630", #<1>
                      "s680", "s910", "Dark", "Clear") #<1>

gain.ratios <- #<2>
  tibble( #<2>
    gain = c(0.5, 1, 2, 4, 8, 16, 32, 64, 128, 256, 512), #<2>
    gain.ratio = #<2>
      c(0.008, 0.016, 0.032, 0.065, 0.125, 0.25, 0.5, 1, 2, 3.95, 7.75) #<2>
  ) #<2>

#normalize data:
dataVEET2 <-
  dataVEET2 |> 
  mutate(across(c(s415:Clear), \(x) (x - Dark)/integration_time)) |>  #<3> 
  normalize_counts( #<4>
    gain.columns = rep("Gain", 11), #<5> 
    count.columns = count.columns, #<6>
    gain.ratios #<7>
  ) |> 
  select(-c(s415:Clear)) |> #<8>
  rename_with(~ str_remove(.x, ".normalized")) #<8>
```

1.  Column names of variables that need to be normalized
2.  Gain ratios as specified by the manufacturer's reconstruction guide
3.  Remove dark counts & scale by integration time
4.  Function to normalize counts
5.  All sensor channels share the gain value
6.  Sensor channels to normalize (see 1.)
7.  Gain ratios (see 2.)
8.  Drop original raw count columns

In this call, we specified `gain.columns = rep("Gain", 11)` because we have 11 sensor columns that all use the same gain factor column (`Gain`). This step will add new columns (with a suffix, e.g. `.normalized`) for each spectral channel representing the count normalized by the gain. We then dropped the `raw` count columns and renamed the normalized ones by dropping `.normalized` from the names. After this, `dataVEET2` contains the normalized sensor readings for `s415, s445, ..., s940, Dark, Clear` for each time point time point.

Because we do not need this high a resolution, we will **aggregate it to a 5-minute interval** for computational efficiency. The assumption is that spectral composition does not need to be examined at every 2-second instant for our purposes, and 5-minute averages can capture the general trends while drastically reducing data size and downstream computational costs.

```{r}
#| filename: Aggregate spectral data to 5-minute intervals and mark gaps
dataVEET2 <- dataVEET2 |>
  aggregate_Datetime(unit = "5 mins") |>     #<1>
  gap_handler(full.days = TRUE) |>           #<2> 
  add_Date_col(group.by = TRUE) |>           #<3> 
  remove_partial_data(Clear, threshold.missing = "1 hour") #<4> 
```

1. Aggregate to 5-minute bins
2. Explicit NA for any gaps in between
3. Add a date identifier for grouping
4. remove days with more than one hour of data missing

We aggregate over 5-minute windows; within each 5-minute bin, multiple spectral readings (if present) are combined (averaged). We use one of the channels (here `Clear`) as the reference variable for `remove_partial_data` to drop incomplete days (the choice of channel is arbitrary as all channels share the same level of completeness).

::: callout-warning
Please note that `normalize_counts()` requires the `Gain` values according to the gain table. If we had aggregated the data before normalizing it, `Gain` values would have been averaged within each bin (5 minutes in this case). If the `Gain` did not change in that time, it is not an issue. Any mix of `Gain` values will lead to a `Gain` value that is not represented in the gain table. While outputs for `normalize_counts()` are not wrong in these cases, they will output `NA` if a `Gain` value is not found in the table. Thus we recommend to always normalize counts based on the raw dataset.
:::

### Spectral reconstruction

For spectral reconstruction, we require a calibration matrix that corresponds to the `VEET`’s sensor channels. This matrix would typically be obtained from the device manufacturer or a calibration procedure. It defines how each channel’s normalized count relates to intensity at various wavelengths. For demonstration, the calibration matrix was provided by the manufacturer and is specific to the make and model (see @fig-calibmtx). It should not be used for research purposes without confirming its accuracy with the manufacturer.

![Calibration matrix](assets/calibration_matrix.png){#fig-calibmtx}

```{r}
#| filename: Calibration matrica and reconstruction of spectral power distribution
calib_mtx <- # <1>
  read_csv("data/VEET_calibration_matrix.csv", # <1>
           show_col_types = FALSE) |> # <1>
  column_to_rownames("wavelength") # <1>

dataVEET2 <-
  dataVEET2 |> 
  mutate(
  Spectrum = 
    spectral_reconstruction( # <2>
      pick(s415:s910),   # <3>
      calibration_matrix = calib_mtx, #<4>
      format = "long" # <5>
  )
)
```

1. Import the calibration matrix and make certain `wavelength` is set a rownames
2.  The function `spectral_reconstruction()` does not work on the level of the dataset, but has to be called within `mutate()`(or provided the data directly)
3.  Pick the normalized sensor columns
4.  Provide the calibration matrix
5.  Return a long-form list column (wavelength, intensity)

Here, we use `format = "long"` so that the result for each observation is a **list-column** `Spectrum`, where each entry is a tibble[^2] containing two columns: `wavelength` and `irradiance` (one row per wavelength in the calibration matrix). In other words, each row of `dataVEET2` now holds a full reconstructed spectrum in the `Spectrum` column. The long format is convenient for further calculations and plotting. (An alternative `format = "wide"` would add each wavelength as a separate column, but that is less practical when there are many wavelengths.)

To visualize the data we will calculate the photopic illuminance based on the spectra and plot each spectrum color-scaled by their illuminance. For clarity, we reduce the data to observations within the day that has the most observations (non-`NA`).

```{r}
#| filename: Calculate photopic illuminance
data_spectra <- 
dataVEET2 |> 
  sample_groups(order.by = sum(!is.implicit)) |> #<1>
  mutate( 
    Illuminance = Spectrum |> #<2>
      map_dbl(spectral_integration, #<3>
              action.spectrum = "photopic", #<4>
              general.weight = "auto") #<5>
  ) |> 
  unnest(Spectrum) #<6>
data_spectra |> select(Id, Date, Datetime, Illuminance) |> distinct()
```

1.  Keep only observations for one day (with the lowest missing intervals)
2.  Use the spectrum,...
3.  ... call the function `spectral_integration()` for each,...
4.  ... use the brightness sensitivity function,...
5.  ... and apply the appropriate efficacy weight.
6.  Create a long format of the data where the spectrum is unnested

The following plot visualizes the spectra:

```{r}
#| label: fig-VEET-spectra-intensity
#| fig-cap: "Overview of the reconstructed spectra, color-scaled by photopic illuminance (lx)"
#| fig-height: 5
#| fig-width: 8
#| filename: Plot spectra
data_spectra |> 
  ggplot(aes(x = wavelength,group = Datetime)) +
  geom_line(aes(y = irradiance*1000, col = Illuminance)) +
  labs(y = "Irradiance (mW/m²/nm)", 
       x = "Wavelength (nm)", 
       col = "Photopic illuminance (lx)") +
  scale_color_viridis_b(breaks = c(0, 10^(0:3))) +
  scale_y_continuous(trans = "symlog", breaks = c(0, 1, 10, 50)) +
  coord_cartesian(ylim = c(0,NA), expand = FALSE) +
  theme_minimal()
```

The following ridgeline plot can be used to assess **when** in the day certain spectral wavelenghts are dominant:

```{r}
#| label: fig-VEET-spectra-time
#| fig-cap: "Overview of the reconstructed spectra by time of day, color-scaled by photopic illuminance (lx)"
#| fig-height: 5
#| fig-width: 8
#| filename: Plot spectra across the time of day
data_spectra |> 
  ggplot(aes(x = wavelength, group = Datetime)) +
  geom_ridgeline(aes(height = irradiance*1000, 
                     y = Datetime, 
                     fill = Illuminance), 
                 scale = 400, lwd = 0.1, alpha = 0.7) +
  labs(y = "Local time & Irradiance (mW/m²/nm)", 
       x = "Wavelength (nm)", 
       fill = "Photopic illuminance (lx)")+
  scale_fill_viridis_b(breaks = c(0, 10^(0:3))) +
  theme_minimal()
```


At this stage, the `dataVEET2` dataset has been processed to yield time-series of spectral power distributions. We can use these to compute biologically relevant light metrics. For instance, one possible metric is the proportion of power in short wavelengths versus long wavelengths. 

In the main analysis, we defined **short-wavelength (blue light) content** as the integrated intensity in the 400–500 nm range, and **long-wavelength content** as the integrated intensity in a longer range (e.g. 600–700 nm), then computed the *short-to-long ratio* (“sl ratio”). Calculating these metrics is the first step of spectrum analysis in the [main tutorial](index.qmd#spectrum).

```
dataVEET2 |> 
  select(Id, Date, Datetime, Spectrum) |>    # focus on ID, date, time, and spectrum
  mutate(
    short = Spectrum |> map_dbl(spectral_integration, wavelength.range = c(400, 500)),
    long  = Spectrum |> map_dbl(spectral_integration, wavelength.range = c(600, 700)),
    `sl ratio` = short / long   # compute short-to-long ratio
  )
```

*(The cutoff of 500 nm here is hypothetical for demonstration; actual definitions might vary.)* We would then have columns `short`, `long`, and `sl_ratio` for each observation, which could be averaged per day or analyzed further. The cleaned spectral data in `dataVEET2` makes it straightforward to calculate such metrics or apply spectral weighting functions (for melatonin suppression, circadian stimulus, etc., if one has the spectral sensitivity curves).

With the `VEET` spectral preprocessing complete, we emphasize that these steps – normalizing by gain, applying calibration, and perhaps simplifying channels – are **device-specific** requirements. They ensure that the raw sensor counts are translated into meaningful physical measures (like spectral irradiance). Researchers using other spectral devices would follow a similar procedure, adjusting for their device’s particulars (some may output spectra directly, whereas others, like `VEET`, require reconstruction.

::: {.callout-note}

Some devices may output normalized counts instead of raw counts. For example, the `ActLumus` device outputs normalized counts, while the `VEET` device records raw counts and the gain. Manufacturers will be able to speficy exact outputs for a given model and software version.

:::

## `VEET` Data: Time of flight (distance)

In this last section, the distance data of the `VEET` device will be imported, analogous to the other modalities. The `TOF` modality contains information for up to two objects in a 8x8 grid of measurements, spanning a total of about 52° vertically and 41° horizontally. Because the `VEET` device can detect up to two objects in a given grid point, and there is a confidence value assigned to every measurement, each observation contains $2*2*8*8 = 256$ measurements.

```{r}
#| label: fig-VEET-overview3
#| fig-cap: "Overview plot of imported VEET data"
#| fig-height: 2
#| fig-width: 6
#| filename: Import VEET Spectral Sensor (TOF) data
path <- "data/01_VEET_L.csv.zip"
dataVEET3 <- import$VEET(path, 
                        tz = tz, 
                        modality = "TOF", #<1>
                        manual.id = "VEET" #<2>
                        ) 
```

1.  `modality` is a parameter only the `VEET` device requires. If uncertain, which devices require special parameters, have a look a the import help page (`?import`) under the `VEET` device. Setting it to `TOF` gives us the distance modality.
2.  As we are only dealing with one individual here, we set a manual `Id`

```{r}
#| filename: Print the first 6 rows of the VEETs distance dataset
dataVEET3 |> head()
```


In a first step, we condition the data similarly to the other `VEET` modalities. For computational reasons of the use cases, we remove the second object and set the interval to `10 seconds`. *Note that the next step still takes considerable computation time*.

```{r}
#| label: VEET-cleaning
#| filename: Aggregate distance data to 5-second intervals and mark gaps
dataVEET3 <- 
  dataVEET3 |>
  select(-contains(c("conf2_", "dist2_"))) |> # <1>
  aggregate_Datetime(unit = "5 secs") |>     # <2>
  gap_handler(full.days = TRUE) |>           # <3>
  add_Date_col(group.by = TRUE) |> 
  remove_partial_data(dist1_0, threshold.missing = "1 hour")
dataVEET3 |> summary_overview(dist1_0)
```

1.  Remove the second object (for computational reasons)
2.  Aggregate to 10-second bins
3.  Explicit NA for any gaps

In the next step, we need to transform the `wide` format of the imported dataset into a `long` format, where each row contains exactly one observation for one grid-point. 

```{r}
#| label: VEET-long
#| filename: Pivot distance grid from wide to long
dataVEET3 <- 
  dataVEET3 |> 
  pivot_longer(
    cols = -c(Datetime, file.name, Id, is.implicit, time_stamp, modality, Date),
    names_to = c(".value", "position"),
    names_pattern = "(conf1|conf2|dist1|dist2)_(\\d+)"
  )
```

In a final step before we can use the data in the analysis, we need to assign `x` and `y` coordinates based on the `position` column that was created when pivoting longer. Positions are counted from 0 to 63 starting at the top right and increasing towards the left, before continuing on the right in the next row below. `y` positions thus depend on the row count, i.e., how often a row of 8 values fits into the `position` column. `x` positions consequently depend on the `position` within each 8-value row. We also add an `observation` variable that increases by `+1` every time, the `position` column hits `0`. We then center both `x` and `y` coordinates to receive meaningful values, i.e., 0° indicates the center of the overall measurement cone. Lastly, we convert both confidence columns, which are scaled from 0-255 into percentages by dividing them by `255`. Empirical data from the manufacturer points to a threshold of about `10%`, under which the respective distance data is not reliable.

```{r}
#| label: VEET-distance preparation
#| filename: Calculate grid positions of spatial distance measurements
dataVEET3 <- 
  dataVEET3 |> 
  mutate(position = as.numeric(position),
         y.pos = (position %/% 8)+1, #<1>
         y.pos = scale(y.pos, scale = FALSE)*52/8, #<2>
         x.pos = 8 - (position %% 8), #<3>
         x.pos = scale(x.pos, scale = FALSE)*41/8, #<4>
         observation = cumsum(position == 0), #<5>
         across(starts_with("conf"), \(x) x/255) #<6>
         )
```

1.  Increment the y position for every 8 steps in `position`
2.  Center `y.pos` and rescale it to cover 52° across 8 steps
3.  Increment the x position for every step in `position`, resetting every 8 steps
4.  Center `x.pos` and rescale it to cover 41° across 8 steps
5.  Increase an observation counter every time we restart with `position` at 0
6.  Scale the confidence columns so that 255 = 100%

Now this dataset is ready for further analysis. We finish by visualizing the same observation time on different days. Note that we replace zero distance values with `Infinity`, as these indicate measurements outside the 5m measurement radius of the device.

```{r}
#| label: fig-VEET-distance
#| fig-cap: "Example observations of the measurement grid at 1:14 p.m. for each measurement day. Text values show distance in cm. Empty grid points show values with low confidence. Zero-distance values were replaced with infinite distance and plotted despite low confidence."
#| fig-height: 5
#| message: false
#| fig-width: 12
#| filename: Plot spatial distance grid for the same time point on each day
extras <- list( #<1>
  geom_tile(), #<1>
  scale_fill_viridis_c(direction = -1, limits = c(0, 200), #<1>
                       oob = scales::oob_squish_any), #<1>
  scale_color_manual(values = c("black", "white")), #<1>
  theme_minimal(), #<1>
  guides(colour = "none"), #<1>
  geom_text(aes(label = (dist1/10) |> round(0), colour = dist1>1000), #<1>
            size = 2.5), #<1>
  coord_fixed(), #<1>
  labs(x = "X position (°)", y = "Y position (°)", #<1>
       fill = "Distance (cm)")) #<1>

slicer <- function(x){seq(min((x-1)*64+1), max(x*64, by = 1))} #<2>

dataVEET3 |> 
  slice(slicer(9530)) |>  #<3>
  mutate(dist1 = ifelse(dist1 == 0, Inf, dist1)) |> #<4>
  filter(conf1 >= 0.1 | dist1 == Inf) |> #<5>
  ggplot(aes(x=x.pos, y=y.pos, fill = dist1/10))+ extras + #<6>
  facet_grid(~Datetime) #<7>
```

1.  Set visualization parameters
2.  Allows to choose an observation
3.  Choose a particular observation
4.  Replace 0 distances with Infinity
5.  Remove data that has less than 10% confidence
6.  Plot the data
7.  Show one plot per day

As we can see from the figure, different days have - at a given time - a vastly different distribution of distance data, and measurement confidence (values with confidence \< 10% are removed)

### Removing non-wear times

Same as for the `ALS` modality, we can remove times of non-wear - we can even use the same `wear_data` set for it:

```{r}
#| filename: Removing non-wear observations from distance modality
dataVEET3 <- 
  dataVEET3 |> 
  group_by(Id) |> 
  add_states(wear_data)

dataVEET3 <- 
  dataVEET3 |> 
  mutate(dist1 = ifelse(wear, dist1, NA)) |> 
  group_by(Id, Date)
```


## Saving the Cleaned Data

After executing all the above steps, we have three cleaned data frames in our R session:

-   **`dataCC`** – the processed `Clouclip` dataset (5-second intervals, with distance and lux, including NA for gaps and sentinel statuses).

-   **`dataVEET`** – the processed `VEET` ambient light dataset (5-second intervals, illuminance in lux, with gaps filled).

-   **`dataVEET2`** – the processed `VEET` spectral dataset (5-minute intervals, each entry containing a spectrum or derived spectral metrics).

-   **`dataVEET3`** – the processed `VEET` distance dataset (5-second intervals, each entry containing the distance of up to two objects in the 8x8 grid).

For convenience and future reproducibility, we will save these combined results to a single R data file. Storing all cleaned data together ensures that any analysis can reload the exact same data state without re-running the import and cleaning (which can be time-consuming for large raw files).

```{r}
#| filename: Save preprocessed files
if (!dir.exists("data/cleaned")) dir.create("data/cleaned", recursive = TRUE) #<1>
save(dataCC, dataVEET, dataVEET2, dataVEET3, file = "data/cleaned/data.RData") #<2>
```

1. Create directory for cleaned data if it doesn't exist
2. Save all cleaned datasets into one .RData file

The above code creates (if necessary) a folder `data/cleaned/` and saves a single RData file (`data.RData`) containing the three objects. To retrieve them later, one can use `<- load("data/cleaned/data.RData")`, which will return the objects into the environment. This single-file approach simplifies sharing and keeps the cleaned data together.

In summary, this supplement has walked through the full preprocessing pipeline for two example devices. We began by describing the raw data format for each device and then demonstrated how to import the data with correct time zone settings. We handled device-specific quirks like sentinel codes (for `Clouclip`) and multiple modalities with gain normalization (for `VEET`). We showed how to detect and address irregular sampling, how to explicitly mark missing data gaps to avoid analytic pitfalls, and how to reduce data granularity via rounding or aggregation when appropriate. Throughout, we used functions from LightLogR in a tidyverse workflow, aiming to make the steps clear and modular. By saving the final cleaned datasets, we set the stage for the computation of visual experience metrics such as working distance, time in bright light, spectral composition ratios, as presented in the [main tutorial](index.qmd). We hope this detailed tutorial empowers researchers to adopt similar pipelines for their own data, facilitating reproducible and accurate analyses of visual experience.

## References
```
